# Selfing is the safest sex for *Caenorhabditis tropicalis*

**DOI:** 10.1101/2020.08.07.242032

**Authors:** Luke M. Noble, John Yuen, Lewis Stevens, Nicolas Moya, Riaad Persaud, Marc Moscatelli, Jacqueline Jackson, Gaotian Zhang, Rojin Chitrakar, L. Ryan Baugh, Christian Braendle, Erik C. Andersen, Hannah S. Seidel, Matthew V. Rockman

## Abstract

Mating systems have profound effects on genetic diversity and compatibility. The convergent evolution of self-fertilization in three *Caenorhabditis* species provides a powerful lens to examine causes and consequences of mating system transitions. Among the selfers, *C. tropicalis* is the least genetically diverse and most afflicted by outbreeding depression. We generated a chromosomal-scale genome for *C. tropicalis* and surveyed global diversity. Population structure is very strong, and islands of extreme divergence punctuate a genomic background that is highly homogeneous around the globe. Outbreeding depression in the laboratory is caused largely by multiple gene drive elements, genetically consistent with maternal toxin/zygotic antidote systems. Driver loci harbor novel and duplicated genes, and their activity is modified by mito-nuclear background. Segregating drivers dramatically reduce fitness, and simulations show that selfing limits their spread. Frequent selfing in *C. tropicalis* may therefore be a strategy to avoid drive-mediated outbreeding depression.

## Introduction

Sex and outcrossing are common, but costly, and taxa have repeatedly evolved mating systems to avoid them. This sets advantages of selfing, such as reproductive assurance, against long-term adaptability (Otto, 2009). Selfing has profound consequences for evolution due to changes in effective recombination, homozygosity, and migration, leading to a net reduction in effective population size. Mixed mating systems, combining some form of selfing with occasional outcrossing, are a frequent compromise (Chelo et al., 2019; Cutter, 2019; Escobar et al., 2011; Goodwillie et al., 2005; Igic & Kohn, 2006; Jarne & Auld, 2006).

Variation in mating systems is especially familiar in plants, but has also been one of the longstanding attractions of nematode biology (Nigon & Félix, 2017). Just within Rhabditidae, this aspect of life-history now spans systems with separate males and females (gonochorism), males and self-fertile hermaphrodites (androdioecy; (Kanzaki et al., 2017; Mayer et al., 2007)), males, females, and hermaphrodites (trioecy; (Chaudhuri et al., 2015; Kanzaki et al., 2017)), asexual reproduction where sperm does not contribute genetic material (parthenogenesis, gynogenesis; (Fradin et al., 2017; Grosmaire et al., 2019)), and alternating generations of hermaphroditism and dioecy (heterogony; (Kiontke, 2005)).

Within the *Caenorhabditis* genus, the androdioecious system of males and self-fertilizing hermaphrodites has evolved three times independently (Ellis, 2017). Hermaphrodites are morphologically female, but during larval development they generate and store sperm for use as adults. Hermaphrodites cannot mate with one another, and in the absence of males all reproduction is by self fertilization. Mating system evolution, together with the unknown biology and population genetics of outcrossing ancestral species, seems to have played out to different effect across the three species, *C. elegans, C. briggsae,* and *C. tropicalis. C. elegans* and *C. briggsae,* isolated in 1899 (Maupas, 1899, 1900) and 1944 (Briggs Gochnauer & McCoy, 1954; Dougherty & Nigon, 1949), respectively, are relatively cosmopolitan (Cutter, 2015; Frezal & Felix, 2015; Kiontke & Sudhaus, 2006). *C. briggsae* is the most broadly distributed, frequently sampled, and genetically diverse of the three. It shows strong latitudinal population structure differentiating temperate and tropical clades (Cutter et al., 2006; E. S. Dolgin et al., 2008; Ferrari et al., 2017; Prasad et al., 2011; Thomas et al., (2015), although representatives of these clades can also be found together at the local scale (Félix et al., 2013). *C. elegans* shows a clear preference for cooler climates (Crombie et al., 2019; Félix & Duveau, 2012; Kiontke et al., 2011), and genetic diversity is relatively rich in the vicinity of the Pacific. Uneven sampling and data across species make generalizations fraught, however. The historical view of *C. elegans* as a near-clonal invasive species, with little genetic diversity from the lab-adapted N2 reference but for a few outliers, has been upended by systematic sampling and sequencing efforts (Cook et al., 2017; Crombie et al., 2019; Lee et al., 2020).

*C. tropicalis* was first identified by Marie-Anne Félix in 2008 (Kiontke et al., 2011), and investigations of its reproductive biology have shed light on the mechanistic basis of transitions to selfing (Wei et al., 2014; Zhao et al., 2018). However, a comprehensive reference genome for the species is lacking and relatively little is known of its biology and ecology. Global sampling indicates a more restricted range than that of the other selfers (Felix, 2020), and the single study of *C. tropicalis* population genetics and reproductive compatibility found extremely low levels of genetic diversity at a handful of loci (Gimond et al., 2013). Crosses among, and sometimes within, locales revealed outbreeding depression, a result common to the selfers (Baird & Stonesifer, 2012; Elie S. Dolgin et al., 2007; Ross et al., 2011) but in stark contrast to outbreeding species, where estimated diversity is often orders of magnitude higher and inbreeding depression can be severe (Barriere et al., 2009; Elie S. Dolgin et al., 2007; Gimond et al., 2013). These effects were particularly acute in *C. tropicalis,* with frequent embryonic lethality and developmentally abnormal F_2_ progeny among certain hybrid crosses (Gimond et al., 2013). Male mating ability was also found to be generally poor, though highly variable, which together with low genetic diversity suggests an especially high rate of selfing.

To study the effects of selfing on genomes and population structure, and establish genetic and genomic resources for *C. tropicalis,* we assembled a chromosome-scale genome, oriented by linkage data from recombinant inbred lines. We show with short-read mapping against this reference that the population structure of a global sample of isolates is very strong, which is an expected side-effect of selfing. We find extreme heterogeneity in the distribution of genetic diversity across the genome, leading to a relatively weak association between recombination rate and nucleotide diversity. We also show, using a second highly contiguous assembly, that in the face of this heterogeneity reference-based mapping vastly underestimates the true levels of genetic diversity in the species, which at some loci approach current estimates for dioecious species. Finally, we investigate the causes of strong outbreeding depression, another expected side-effect of selfing, in a cross between divergent isolates. Widespread outbreeding depression observed in the three species of selfing *Caenorhabditis* has been interpreted as evidence for the well understood process of Dobzhansky-Muller epistasis. Here, we show that it is due largely to a different process in *C. tropicalis·.* maternal-effect gene drive haplotypes that kill offspring that do not inherit them. We hypothesize that gene drive selects for selfing in this species, in other words, frequent selfing may be a consequence of outbreeding depression as much as its cause.

## Results

### Heritable variation in outcrossing rates

To better understand variation in selfing and outcrossing in *C. tropicalis,* we tested a global sample of five isolates for their propensity or ability to outcross. For each of these strains, collected from Hawaii, Panama, French Guiana, Cape Verde, and Reunion Island, we assayed the probability that an individual would produce cross progeny when paired with a single individual of the other sex. In factorial crosses, we observed significant variation among strains in their crossing probability as males and as hermaphrodites (Figure 1A). We also observed interaction effects, where the average crossing probability of male and hermaphrodite strains was not predictive of the success of the strains in combination. At one extreme, the pairing of JU1373 hermaphrodites from Réunion with JU1630 males from Cape Verde, yielded no cross progeny from 22 trials, while at the other extreme, Hawaiian QG131 males and South American NIC58 hermaphrodites yielded cross progeny in each of 34 trials. Male crossing probability was much more variable than that of hermaphrodites (residual deviance of 85.2 vs. 179.2, null deviance 232.6, binomial linear model). Same-strain pairings were indistinguishable from inter-strain pairings (p=0.88, likelihood ratio test (LRT) of binomial linear models). This analysis shows that wild isolates of *C. tropicalis,* and males in particular, vary greatly in their propensity or ability to outcross.

**Figure 1.**
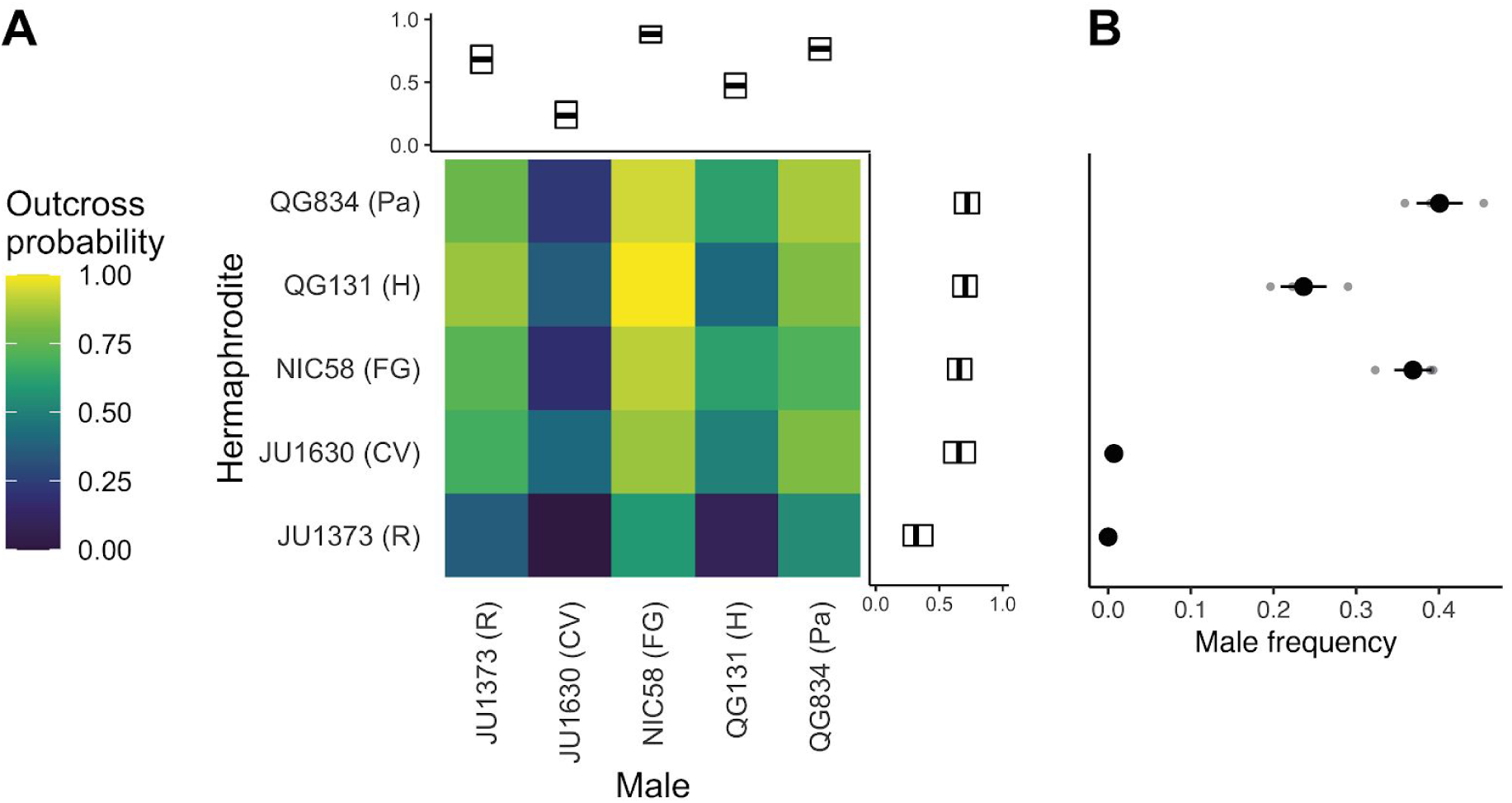
(**A**) Outcrossing probability in reciprocal crosses. Mating success was scored as a binary trait in 22–34 trials (biological replicates) per cross. Marginal means with bootstrap 99% confidence intervals are shown. (**B**) After 10 generations of passaging, strains vary in their male frequency (mean and standard error of three biological replicates). R: Reunion Island, CV: Cape Verde, FG: French Guiana, H: Hawaii, Pa: Panama. Data are in Figure 1 – source data 1 and 2.

Given extensive variation in outcrossing and the independence of male and hermaphrodite components of this trait, we asked whether strains could maintain males over time, or whether hermaphrodite selfing would drive them from populations. We founded single-strain populations with three hermaphrodites and five males, and passaged their descendants at large population size for ten generations. At the end, two strains, JU1630 and JU1373, had lost males and were reproducing solely by selfing; these are the strains with the lowest cross success in our single-worm pairings (Figure 1B). The other strains retained males at frequencies of 20–40%. This dichotomy resembles that seen among *C. elegans* strains, where some strains become exclusive selfers, while others can maintain high rates of outcrossing (Stewart & Phillips, 2002; Teotonio et al., 2006; Teotonio et al., 2012). We chose the strain with the most-outcrossing males, NIC58 from French Guiana, and the strain with the least-outcrossing hermaphrodites, JU1373 from Réunion Island, as focal strains for an investigation of the genetics, genomics, and population biology of *C. tropicalis.*

### Generation of a chromosomal genome for NIC58

To assemble a reference genome for *C. tropicalis* we used deep PacBio long-read sequencing of NIC58, and then applied genetic linkage data from recombinant inbred lines (RILs) to assess and orient the assemblies. RILs were derived from a cross between a JU1373 hermaphrodite and a NIC58 male, by selfing F_2_ hermaphrodites from a single F_1_ for 10 generations. We genotyped lines by shotgun sequencing, calling diallelic single nucleotide variants (SNVs), inferring parental ancestry by Hidden Markov Model (HMM), and using data for 119 RILs for genetic map estimation. During RIL construction, 12.2% of lines did not survive, consistent with effects of outbreeding depression. The genotypes of the surviving RILs revealed strong segregation distortion favoring JU1373 alleles at two loci on chromosomes III and V, which we return to later.

We used the RIL recombination data to evaluate five long-read assemblies from four assemblers (ra, (Vaser & Sikic, 2019); canu, (Koren et al., 2017); wtdb2, (Ruan & Li, 2019); and flye, (Kolmogorov et al., 2019)) and found that within-chromosome chimeras were common. The estimated genome size for *C. tropicalis* is around 80 Mb (Fierst et al., 2015) – by far the smallest of the selfers, though not the smallest in the genus (Stevens et al., 2019) – and assemblies varied in their naive contiguity from NG50 of 1.6 Mb for ra to 4.9 Mb for flye (half the expected genome size was in sequences of at least this length). One of two flye assemblies was fully concordant with the genetic data, with 36 sequences (>20 kb) spanning 81.8 Mb. We used information from junction-spanning sequences in other assemblies, then local mapping of long reads, to close gaps of estimated 0 cM genetic distance in this assembly. Five gaps of greater than 0 cM remain, which will require longer reads to resolve. The resulting nuclear genome comprised 81.3 Mb in 15 sequences. The X chromosome assembled as a single contig, and all other chromosomes were oriented genetically into pseudochromosomes. We assembled a 13,935 bp mitochondrial genome from short-reads, followed by circular extension with long-reads. To better assess genetic variation between the RIL founder strains, we also assembled draft nuclear (81 Mb span, 4.2 Mb NG50) and mitochondrial (13,911 bp) genomes for JU1373. We do not attempt to bring this assembly to pseudochromosomes here. We annotated nuclear genomes using mixed-stage short-read RNAseq data, calling 21,210 protein-coding genes for NIC58 and 20,829 for JU1373, and annotated the mitochondrial genomes by homology. Thus, this pipeline provided a high-quality genome assembly for NIC58 and a highly contiguous draft genome for JU1373.

To evaluate genome completeness, we tallied single copy orthologs and telomeric repeats for the NIC58 genome and four other *Caenorhabditis* species with chromosomal genomes: *C. elegans* (VC2010, genome reference strain for the canonical lab strain N2; (Yoshimura et al., 2019)), *C. briggsae* (Ross et al., 2011), *C. remanei* (Teterina et al., 2020), and *C. inopinata* (Kanzaki et al., 2018). Gene completeness was similar to that of the other Elegans group genomes, with 97.9% of 3131 nematode single copy orthologs present and complete in the *C. tropicalis* genome (97.9% for *C. inopinata* and *C. briggsae,* 98.4% for *C. remanei,* 98.5% for *C. elegans*; (Seppey et al., 2019)). Telomeres in *C. elegans* are built on repeats of the hexamer TTAGGC (Wicky et al., 1996), and total repeat length varies several-fold among *C. elegans* isolates around a median of 12.2 kb (Cook et al., 2015). Despite their small size, telomeres are difficult to assemble due to subtelomeric repeats (Yoshimura et al., 2019), and no chromosome-scale reference genome in the Elegans group possesses telomeric repeats (defined as chromosome-wide maxima) at all chromosome termini (see also Subirana & Messeguer (2017) for *C. nigoni).* In the NIC58 assembly, four of six chromosomes terminated in telomeric repeats at one end. All were unusually long, with a mean per-terminus repeat length four times that of the *C. elegans* VC2010 genome. Comparisons among assemblies are confounded by technical differences in data and assembly pipelines (total repeat length for the VC2010 assembly is more than six-fold that of the current N2 assembly, for instance), but the comparison to VC2010 is a fair one as both it and NIC58 are based on >250x PacBio long reads. Thus, differences in repeat length may reflect a real difference in telomere biology between *C. tropicalis* and its relatives.

### Recombination maps vary among androdioecious *Caenorhabditis* species

Recombination along the holocentric autosomes of other *Caenorhabditis* species is broadly structured into domains, with crossovers effectively absent from chromosome tips, rare in centers, and common on arms (Rockman & Kruglyak, 2009; Ross et al., 2011). The pattern of recombination in *C. tropicalis,* inferred from the genotypes of the RILs, was similar to that inferred from RILs in *C. elegans* and *C. briggsae* (Figure 2), though more extreme in at least two ways. First, autosomal recombination rate domain structure was more pronounced in *C. tropicalis,* due to longer tip and shorter arm domains in both absolute and relative terms (Figure 2B). *C. tropicalis* tips span 9.6 Mb in sum, versus 4.1 Mb in *C. elegans* and 3 Mb in *C. briggsae* (the latter potentially an underestimate due to genome incompleteness), or 12%, 4% and 3% of the genome, respectively. Assuming a single crossover per chromosome per meiosis (Hillers et al., 2017) and a similar per-domain crossover probability across species, this would cause differences in heterogeneity among species. Recombination in the centers of *C. tropicalis* autosomes was also less frequent on average, however. Second, while arm-center structure was less obvious on the X than on autosomes in all species, this was also most pronounced in *C. tropicalis* (Figure 2). Recombination on the left arm of the *C. tropicalis* X was more frequent than in the center, but the domain segmentation is uncertain given the relatively low contrast.

**Figure 2.**
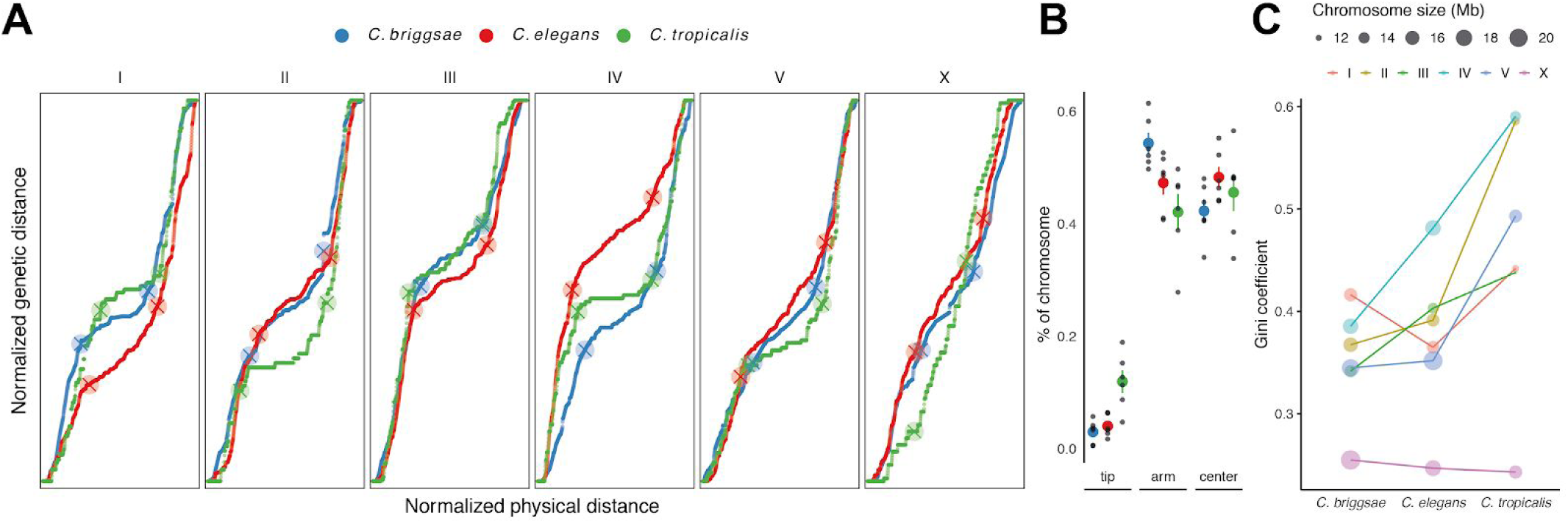
(**A**) Marey maps for the three androdioecious *Caenorhabditis* species are qualitatively similar. In these plots of genetic position as a function of physical position, the slope is an estimate of the local recombination rate. Recombination rate boundaries from a three-domain segmentation (excluding chromosome tips) are shown as crossed points for each species. (**B**) The size of non-recombining chromosome tips is larger on average in *C. tropicalis*, and arms are smaller, both relative to chromosome size and in absolute terms (not shown). Black points are per chromosome values, colored points are species means (± standard error). The variance among chromosomes in the relative sizes of recombination rate domains is also much smaller in *C. briggsae* and *C. elegans* (p < 0.01 for the among-species dispersion component of a double binomial linear model). (**C**) The distribution of recombination among chromosome arms and centers is more heterogeneous, as quantified by the Gini coefficient, a measure of heterogeneity that ranges from 0 (total uniformity) to 1 (all recombination in a single interval; (Kaur & Rockman, 2014)). Maps are based on 119 RILs for *C. tropicalis*, 236 for *C. elegans* (Rockman & Kruglyak, 2009) and 167 for *C. briggsae* (Ross et al., 2011). Data are in Figure 2 – source data 1 and 2.

Many aspects of *Caenorhabditis* DNA sequence and chromosome organization covary with recombination rate (Jovelin et al., 2013; Woodruff & Teterina, 2020). We show this for four other species with chromosome-scale assemblies – *C. briggsae, C. remanei, C. elegans,* and *C. inopinata -* and confirmed that it also holds for the assembled NIC58 genome. DNA sequence repetitiveness, gene density, and GC-content are all, on average, strongly associated with recombination (Figure 2 – figure supplement 1), though variably so across chromosomes and species. Notably, X chromosome differentiation is, as expected, very weak, and of the three species with genetically defined recombination-rate domains, the mean pattern of GC-content across chromosomes is inverted in *C. briggsae* relative to the other selfers.

**Figure 2- figure supplement 1.**
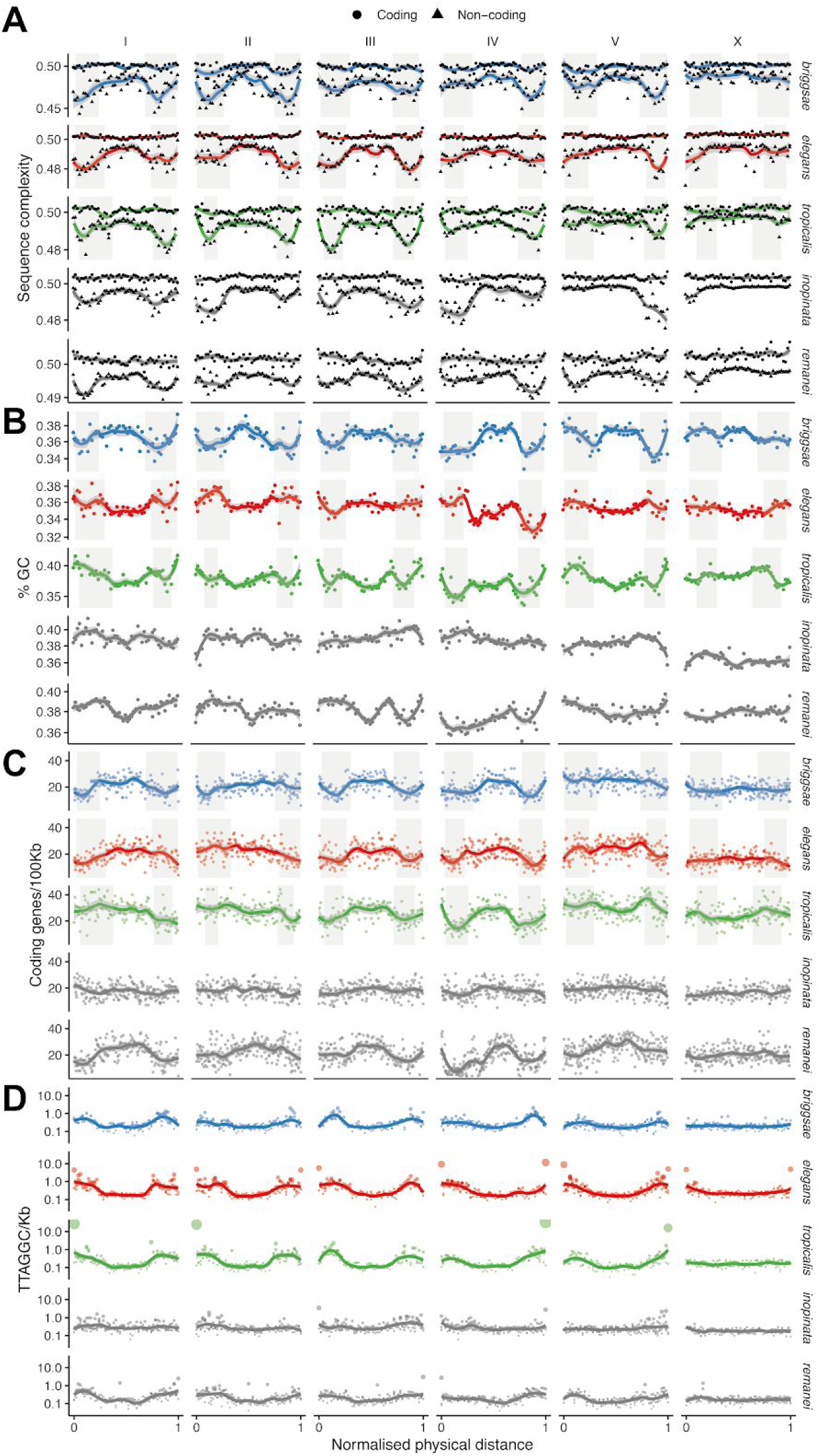
Covariation of chromosome organization with recombination. (**A**) DNA sequence complexity, measured as the maximum zlib compression ratio of 100 bp strings in coding and non-coding sequence. (**B**) GC-content (log scale). (**C**) Gene density. (**D**). Telomeric repeat density. Statistics in A-C are measured in 100 kb non-overlapping windows, then means (points) and a locally-weighted polynomial (LOESS) fit to the raw data are shown for normalized physical distance in 2% bins. Outliers beyond the 98th percentile in median deviation are excluded in A-C, and scales vary across species in A and B. Arm recombination rate domains are shaded where applicable. Data are in Figure 2 – source data 3.

### Macrosynteny is conserved despite extensive intrachromosomal rearrangement

The chromosome-scale genome allows testing of the generality of chromosomal evolution in the *Caenorhabditis* genus, including a dearth of large-scale structural variation that could contribute to outbreeding depression (Cutter et al., 2009; Fierst et al., 2015; Hillier et al., 2007; Jovelin et al., 2013; Kanzaki et al., 2018; Teterina et al., 2020). To examine patterns of macrosynteny, we established orthology relationships for canonical proteins across the five species (Figure 3). This analysis mostly recapitulated known patterns, in that changes in synteny occur predominantly through rearrangement between arms of the same chromosome, and structural evolution of the X chromosome is generally conservative relative to autosomes. To characterize rates of chromosomal rearrangement in *C. tropicalis* while accounting for phylogeny, we compared the number of changes that are unique to the *C. tropicalis* branch to those that are unique to the *C. briggsae* branch. There are just 12 cases of interchromosomal exchanges of one-to-one orthologs unique to the *C. tropicalis* branch (i.e., of genes that are found on the same chromosome in the other four species) versus 48 for *C. briggsae.* Within-chromosome movement across arms is also lower (1068 vs. 1196 cases of 5134 orthologs that are positionally conserved in both *C. remanei* and *C. inopinata).* Because the *C. briggsae* branch is shorter than the *C. tropicalis* branch (Figure 3), we can robustly interpret lower numbers of branch-specific changes in *C. tropicalis* than in *C. briggsae* as lower rates of change on the *C. tropicalis* lineage.

**Figure 3.**
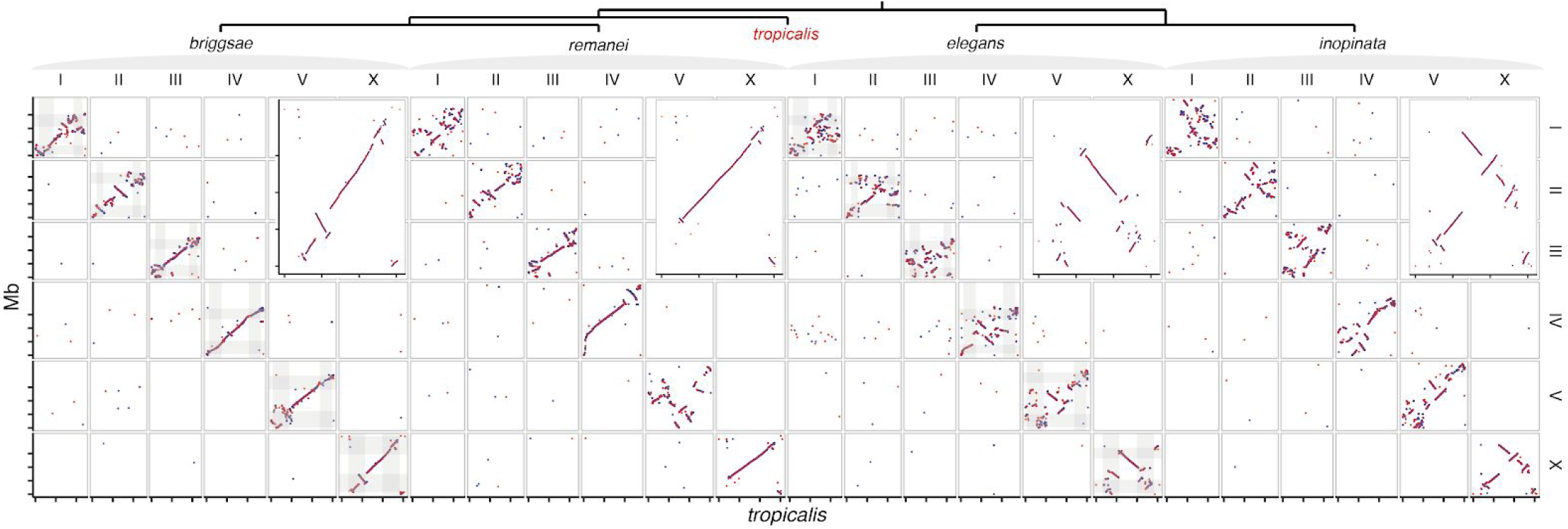
Macrosynteny based on 7983 one-to-one orthologs (colored by strand in the query species). Insets show the X chromosome, where the terminal tip domains are structurally divergent in *C. tropicalis* (query species on the y-axis, *C. tropicalis* on the x-axis). Arm recombination rate domains are shaded in the main panels for the three selfing species with genetic maps. Axes are in physical distance, with tick-marks every 5 Mb. Species relationships are shown in the cladogram above (Stevens et al., 2019). Data are in Figure 3 – source data 1.

**Figure 3- figure supplement 1.**
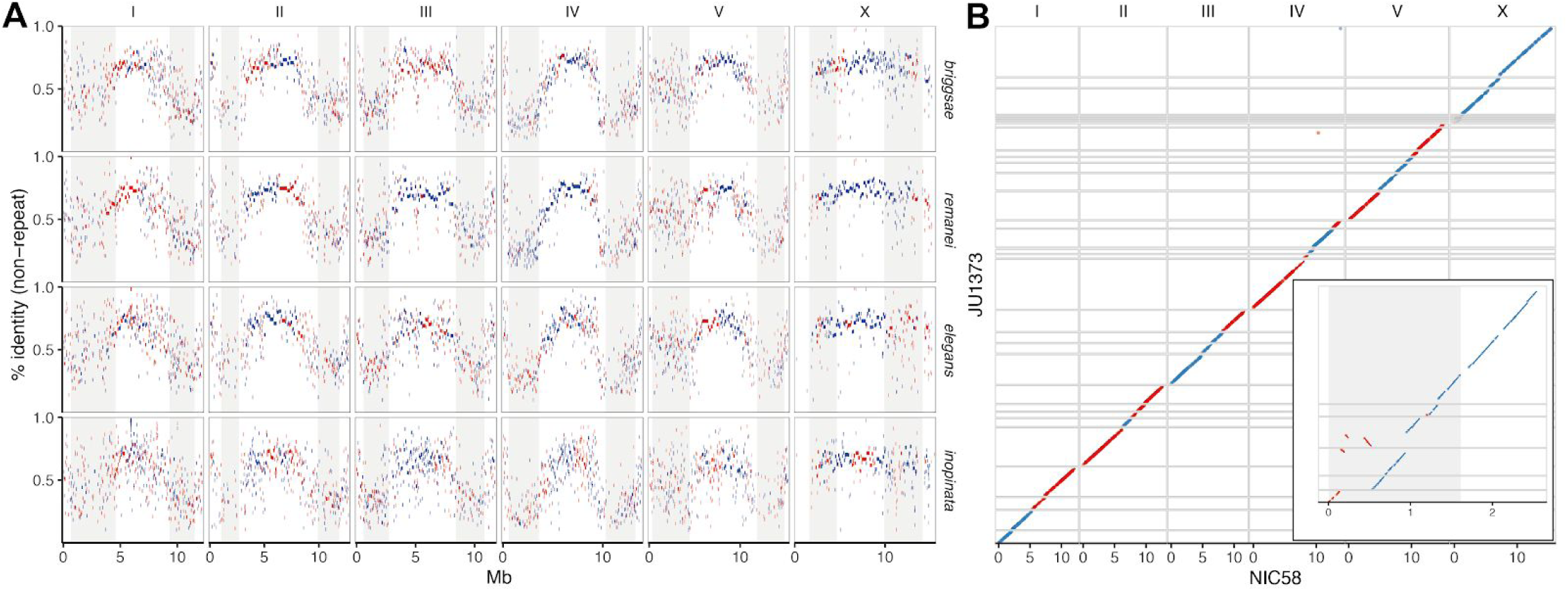
(**A**) Genome coverage and identity with *C. tropicalis* NIC58 as reference (Paten et al., 2011). Aligned blocks are plotted against block identity (percent aligned bases, excluding repetitive sequence) colored by strand in the query species, with arm recombination rate domains shaded. Block size is extended by 30 kb for visibility. Data are in Figure 3 – source data 2. (**B**) Genome alignment between NIC58 and the draft JU1373 assembly shows near-complete colinearity of assembled regions, with the exception of the left tip of the X chromosome (inset, with the genetically-defined tip domain shaded). Alignments (Marçais et al., 2018) are colored by JU1373 assembly strand, grid lines are drawn at sequence boundaries. Data are in Figure 3 – source data 3.

In contrast to the general trend of conservative *Caenorhabditis* X-chromosome evolution, the *C. tropicalis* X has undergone rearrangements in its tips, and hence these align particularly poorly across all species (Figure 3). This poor alignment is not limited to orthologs (i.e., is not due to duplication; see Figure 3 – figure supplement 1A), and represents a major rearrangement in the *C. tropicalis* branch. We found that X-chromosome tips are also polymorphic within *C. tropicalis,* with rearrangements and presence-absence variation observed between JU1373 and NIC58 (Figure 3 – figure supplement 1B). Segregating rearrangements within a tip domain are expected to be relatively free of the deleterious consequences associated with rearrangements elsewhere, as the tips do not recombine (Saito & Colaiacovo, 2017) and therefore would not generate gametes with deficiencies.

### Surveying genetic diversity worldwide

*C. tropicalis* is widely distributed within 25° of the Equator and absent outside this region (Félix, 2020). To begin a global survey of genetic diversity and population structure, we sequenced with short reads an additional 22 isolates that broadly represent the species’ global range. The collection spans Africa, Asia, and South and Central America, but large equatorial regions, notably in Central Africa and Southeast Asia, are not yet represented. Our sample includes 16 American isolates (eight from the Caribbean, eight from Central and South America), four from East Asia, three from Africa and one from the Central Pacific (Figure 5 – source data 1). We called variants against the NIC58 reference genome, hard-filtered to 794,676 biallelic SNVs on the nuclear genome (genotype set 1, Supplementary File 4; see Methods) and selected 397,515 sites with fully homozygous calls and no missing data (genotype set 2, Supplementary File 5) for exploration of population structure. We called mitochondrial variants separately, and filtered similarly, retaining 166 (of 197 hard-filtered) SNVs.

Previously Gimond et al. (2013) sequenced 5.9 kb across nine nuclear, protein-coding loci in 54 isolates (mostly from French Guiana in South America, but including African isolates from Cape Verde and Réunion Island, and our Pacific isolate from Hawaii), and found nine SNVs, equating to a per-site Watterson’s θ around 0.00034. Though not directly comparable, the genome-wide estimate of nucleotide diversity provided here is around three times higher (genotype set 1, median value across 20 kb windows = 0.00097), with mitochondrial diversity higher again (0.0038). These values likely underestimate species-wide variation because of short-read mapping bias – we adjust for missing data, but missing data may not, in fact, be missing from genomes. Indeed, we found rampant missingness in our data; up to 1.3% of the NIC58-alignable fraction of the genome lacks aligned reads among any single isolate, and 7.8% of the alignable genome lacks reads in at least one (considering only those with >25x mapped and paired reads, for which missing data and sequencing depth are uncorrelated, *r =* 0.13, p = 0.65).

### *C. tropicalis* genetic diversity is highly heterogeneous along the genome

Selfing has complex effects on population and genomic evolutionary dynamics. A general expectation is a reduction in effective population size *N_e_* proportional to the frequency of selfing, by a factor of up to two, and the strength of background selection, by potentially much more (Charlesworth, 2012; Nordborg & Donnelly, 1997). The strength of selection acting on genetic variation is proportional to the product of *N_e_* and the selection coefficient s. Selfing is therefore expected to both lower genetic diversity, making evolution more reliant on new mutations, and raise the threshold below which mutations are effectively neutral. Recombination plays a large hand in determining the reach of indirect selection, and is relatively homogenous within recombination rate domains in *Caenorhabditis* (Bernstein & Rockman, 2016; Kaur & Rockman, 2014). Patterns of genetic variation along chromosomes can therefore be potentially indicative of the nature of selection on alleles. We found that the distribution of SNV diversity along all chromosomes is extremely heterogeneous in *C. tropicalis.* Background diversity (median 0_W_) is lowest among the three selfers, and the median number of SNV differences between isolates (π) in 10 kb windows is just 3.2 on chromosome centers, and less than double that on arms (genotype set 2). Variance around the background is almost eight times that of *C. elegans* and more than 100 times that of *C. briggsae* (data in Figure 4A). We used kernel density smoothing of the binned distribution of θ_W_ to partition the genome into segments of very high diversity (the long right tail of divergent outlier regions) and segments with background levels of diversity (e.g., see Figure 4A). Heterogeneity is often highly localized: at 10 kb scale, 141 peaks fall to background within 30 kb or less (see Methods), and divergent regions cover just over 14% of the NIC58 genome in sum. We also found a positive, monotonic relationship between the degree of divergence and enrichment of polymorphism within genes – 64% of all SNVs fall within genes (which occupy 60.3% of the genome), but this rises to 75% for the 1% most divergent regions (Figure 4 – figure supplement 1). This is exacerbated, but not driven, by a strong, non-linear association between read depth and gene proximity. Genetic diversity in *C. tropicalis* is thus typified by a near-invariant background, suggesting very recent global shared ancestry, punctuated by regions of high divergence focused on genes, suggesting balancing selection.

**Figure 4.**
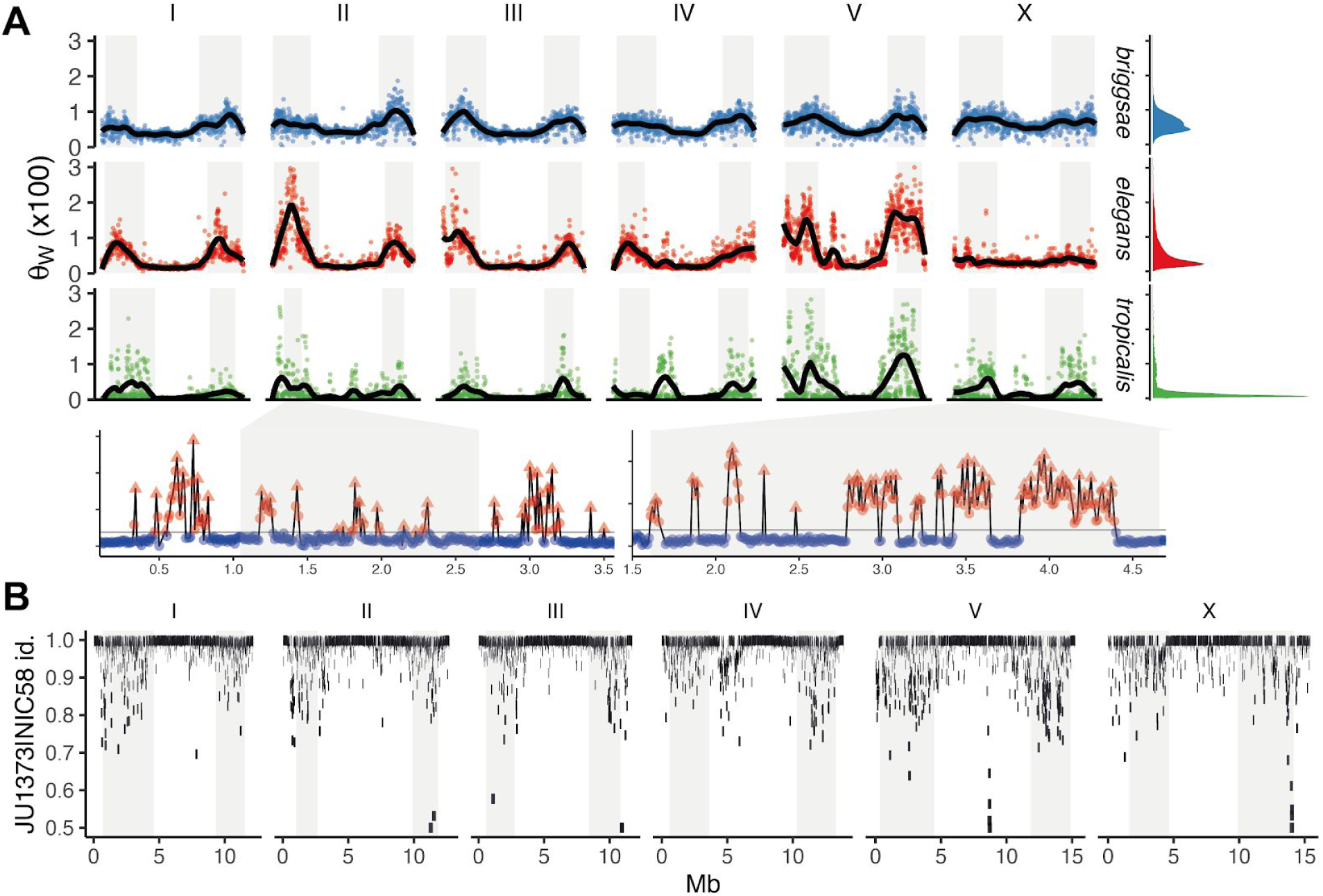
(**A**) Nucleotide diversity across chromosomes (Watterson’s θ in non-overlapping 20 kb windows X 100), based on 24 strains for *C. tropicalis*, 35 strains for *C. briggsae*, and 330 isotypes for *C. elegans*. Differences in heterogeneity across species are apparent from marginal density plots, and from dispersion around the locally-weighted polynomial (LOESS) fit to the data in black. Levels of variation at loci in *C. tropicalis* centers approach those of arms for chromosomes II, IV, V, and X. Arm recombination rate domains are shaded. Regions on the left arms of chromosome II and the X are magnified below, with triangles showing local peaks called at 10 kb scale by segmenting divergent regions (red) from background (blue) at the threshold shown by a grey line (see Methods), and y-axis tick marks as in the main plot. The denominator in Watterson’s estimator uses the mean number of strains with non-missing calls per window rounded to the nearest integer. 12 outliers for *C. elegans* are outside the plotted range. Data are in Figure 4 – source data 1. (**B**) Genetic diversity between JU1373 and the NIC58 reference genome. The measure of variation is the sum of all nucleotide differences in non-overlapping 10 kb windows for all SNV and indel variants, relative to the aligned length in NIC58. Windows are adjusted for missing data, and variation is emphasized: the plotted block size scales with identity (by adding 10^6^log_10_(1 /identity), which is thresholded at 0.5). Data are in Figure 4 – source data 2.

**Figure 4- figure supplement 1.**
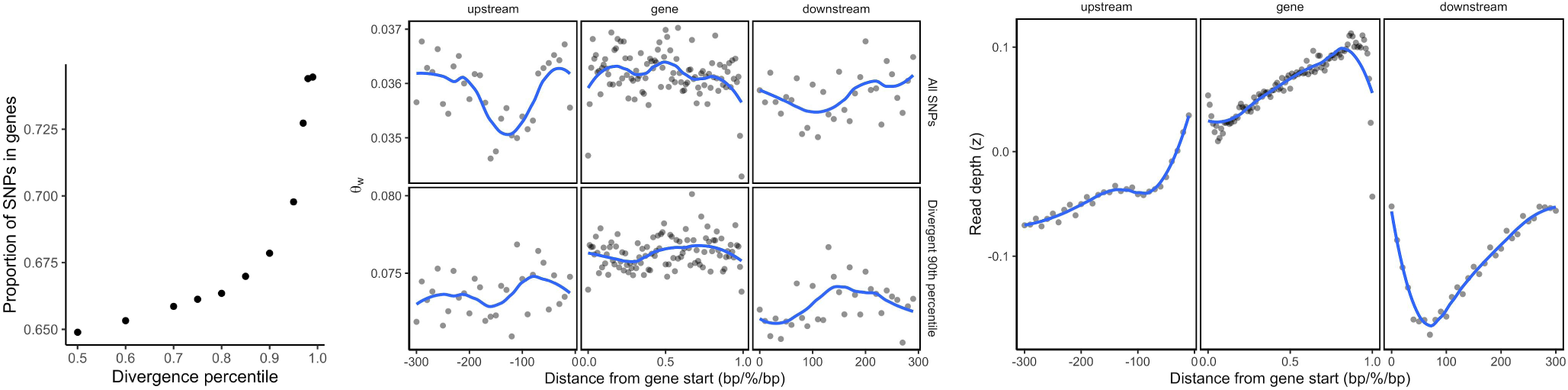
(**A**) Reference-based SNVs in regions of high divergence are concentrated in genes. Divergence is based on the number of SNVs in 10 kb non-overlapping windows. (**B**) The distribution of SNV diversity across genes is shown for normalized gene length (1% bins) and in physical distance for upstream and downstream regions (10 bp bins up to 300 bp). Points show mean values for θ (adjusted for missing data), blue lines are a locally-weighted polynomial (LOESS) fit. Divergent genes are enriched for SNVs relative to flanking background (compare upper row, for all SNVs in genotype set 2, to lower row, for the most divergent 10% of genes). (**C**) Missing data (mean per-strain z-score for aligned read depth) is strongly and non-linearly correlated with gene proximity. Data are based on Figure 4 – source data 3 and Supplementary File 4).

**Figure 4- figure supplement 2.**
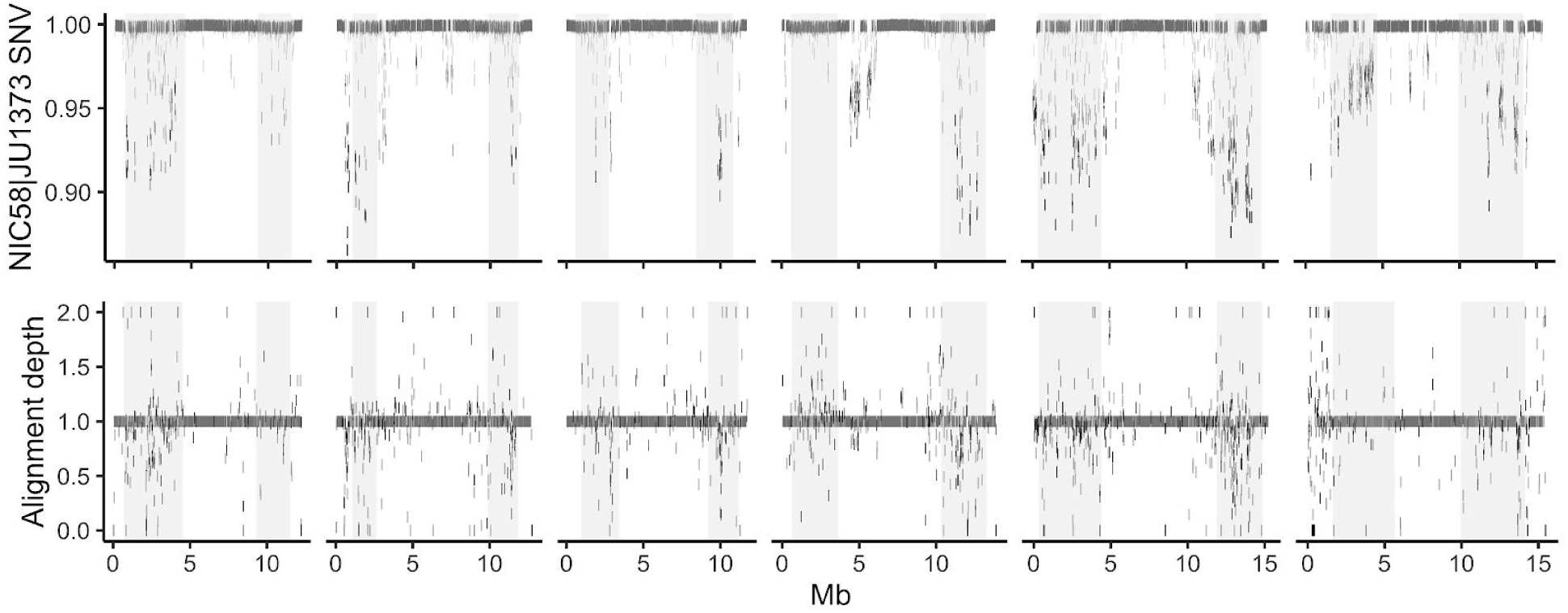
Genetic diversity from alignment of JU1373 to the NIC58 reference genome. Percent identity based on SNVs (upper). Variation is emphasized: plotted block size scales with identity (by adding 10^6^log_10_(1 /identity)) Alignment depth (lower) representing likely copy-number variation (maximum thresholded at 2). Blocks with estimated depth not equal to one are plotted as 50 kb. All statistics are for non-overlapping 10 kb windows adjusted for missing data. Data are in Figure 4 – source data 2.

The positive correlation between recombination rate and nucleotide diversity seen across taxa reflects the pervasive effect of indirect natural selection on diversity at linked sites. In the predominantly selfing *C. elegans* and *C. briggsae,* with streamlined genomes, a karyotype of six holocentric chromosomes, and a single crossover per meiosis, linked selection plays a particularly powerful role in shaping nucleotide diversity (Andersen et al., 2012; Cutter & Payseur, 2003; Graustein et al., 2002; Rockman et al., 2010). The heterogeneous distribution of diversity in *C. tropicalis* results in a much weaker association between recombination rate and diversity. While the recombination-rich chromosome arms are clearly more diverse than centers on average, recombination rate domain is far less predictive of autosomal diversity in *C. tropicalis* (McFadden’s pseudo-*r*^2^ = 0.04, against 0.31 for *C. briggsae* and 0.26 for *C. elegans,* quasibinomial linear models), and within-domain associations that are significant for the other species are not for *C. tropicalis.* Furthermore, in contrast to the weak apparent structuring of recombination rate on the X, nucleotide diversity is highly structured on the X chromosome in *C. tropicalis.* In *C. briggsae* and *C. elegans,* both the average levels of diversity and the variance along chromosomes differ significantly between autosomes and the X (p < 10^ࢤ25^ for higher mean diversity and lower variance on the *C. briggsae* X, p < 10^ࢤ100^ for lower mean and variance on the *C. elegans* X, quasibinomial double generalised linear models, 20 kb scale). In contrast, the *C. tropicalis X* is much more similar to autosomes in these respects (p = 0.13 for mean effect, p < 10^ࢤ5^ for lower variance on the X).

Finally, to gain a view of genetic diversity within *C. tropicalis* less subject to reference-mapping bias, we aligned the draft JU1373 genome against NIC58, calling variants and assessing copy number variation from alignment depth (H. Li, 2018). From 78.86 Mb of aligned bases (69.18 at single copy), we saw a 37% increase in SNVs over reference-based mapping, and a sum of 1.23 Mb in insertion-deletion variation including 388 variants of length greater than 1 kb. SNV divergence in 10 kb windows commonly exceeds 10% on the arms, and total divergence (the sum of variant length differences relative to NIC58) exceeds 30% in windows on every chromosome (Figure 5). Reference-based SNV calling thus dramatically underestimates the true levels of genetic diversity at divergent loci, which are comparable to current estimates for outcrossing species and to analogous patterns recently described in *C. elegans* and *C. briggsae* (Lee et al., 2020). Multiple long-read genomes and variant graph genome representation may be required to more fully describe species-wide variation (Garrison et al., 2018), and deeper population genetic data may allow better inference of foci within these loci.

**Figure 5.**
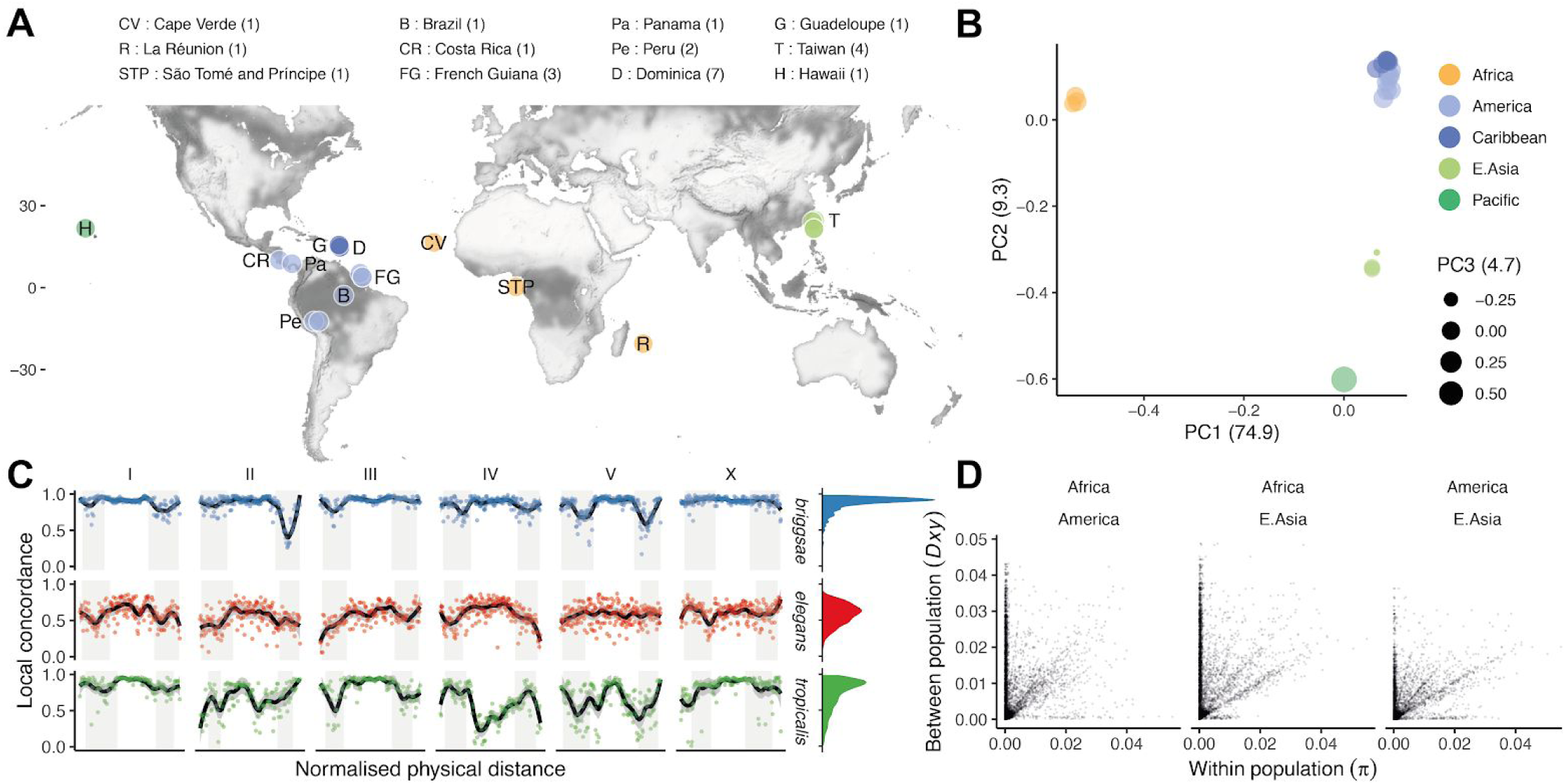
(**A**) The distribution of 24 isolates (numbers per locale are shown in the legend; Data in Figure 5 – source data 1), colored by groupings in (**B**), where principal component analysis of nuclear genomic similarity identifies largely discrete populations. (**C**) Despite strong structure at the genomic level, local ancestry is highly variable in *C. tropicalis*. Each point is the Pearson’s correlation coefficient between relatedness estimated for a 100 kb window (percent identity, including missing data and heterozygous calls) and relatedness at all sites (Stankowski et al. (2019)). (**D**) Genetic diversity is mostly, but not entirely, within populations. For three populations with at least two lines (<99% SNV identity), within population diversity (π) is plotted against between population diversity (Dxy; Nei & Li (1979); 10 kb scale). All correlations are significant, with r^2^ 0.07, 0.09 and 0.25, respectively. Data in B-D are based on Supplementary File 4.

**Figure 5 - figure supplement 1.**
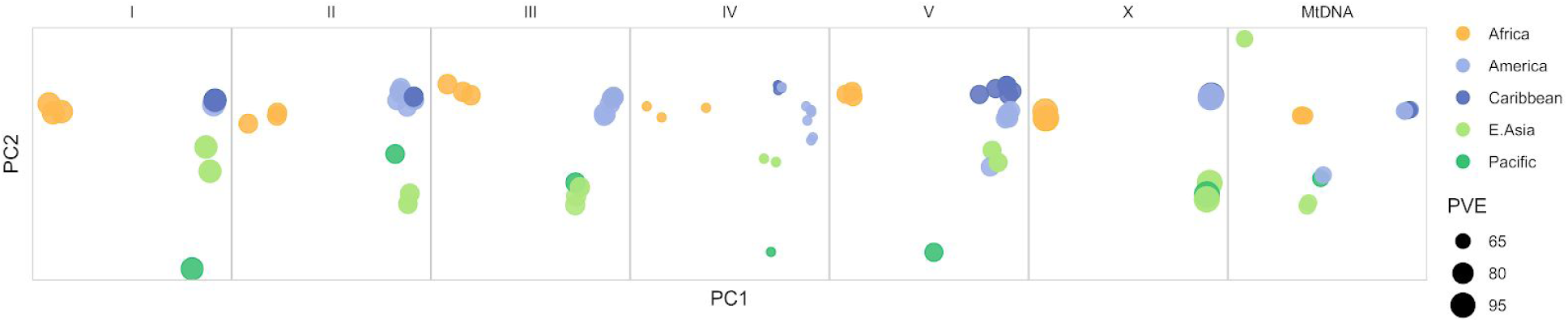
Chromosome and mitochondrial population structure. Point size scales with the percentage of variance explained (PVE) by the first two principal components of genetic relatedness for each chromosome/genome. Data are based on Figure 5 – source data 1.

**Figure 5-figure supplement 2.**
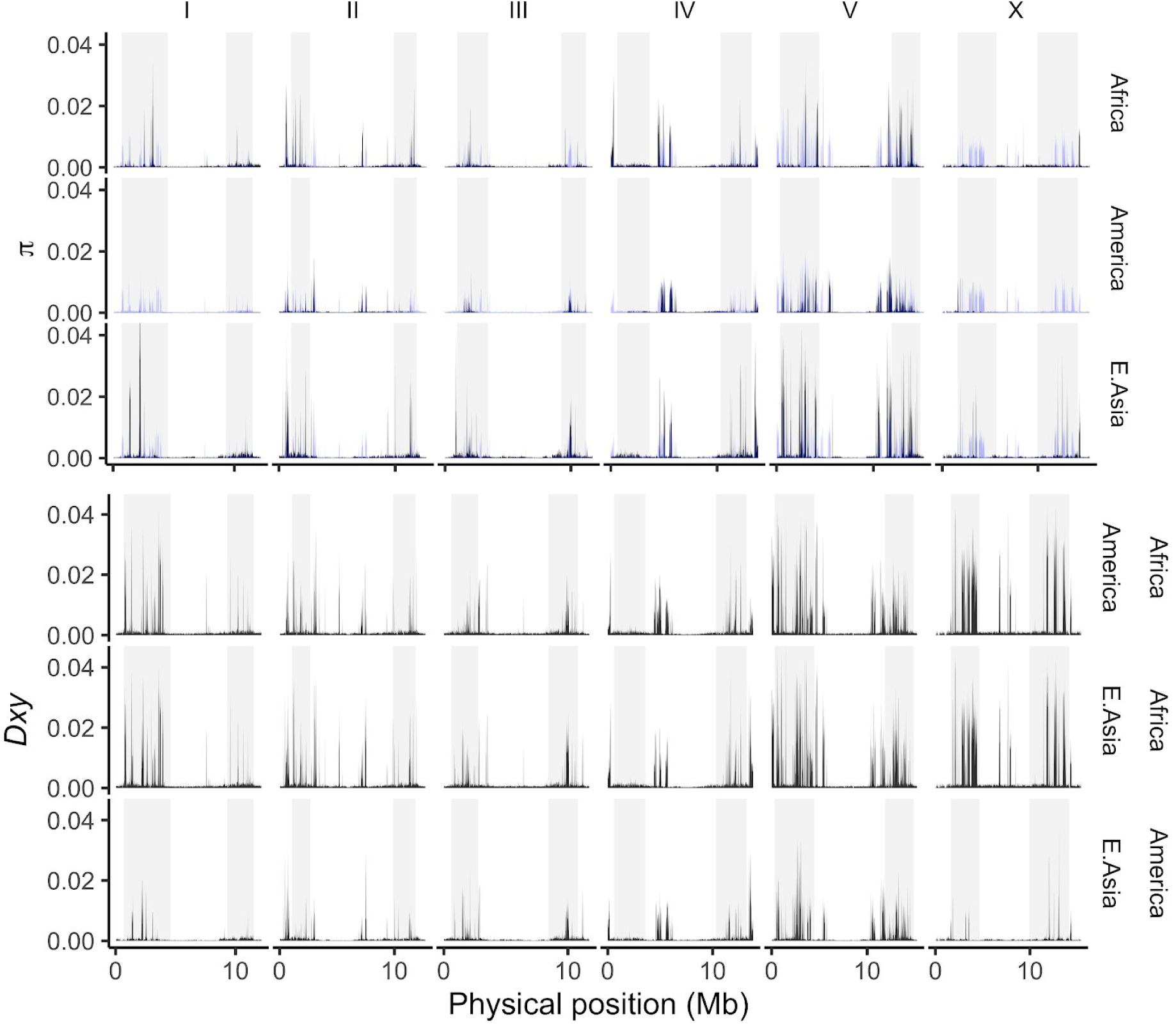
The average number of SNV differences among all pairwise comparisons within (jt; upper, with the global value for all pooled samples plotted in blue across each panel) and between (Nei and Li’s Dxy (1979); lower) populations (10 kb non-overlapping windows, adjusted for the mean fraction of missing data per window). We use the major population groupings defined by genome-wide PCA in Figure 5B, where multiple isolates are present (13 from Central and South America, 3 from Africa and 2 from East Asia after filtering to <99% identity). Data are based on Figure 5 – source data 1.

### *C. tropicalis* shows strong continental population structure with local heterogeneity

To examine population structure worldwide, we decomposed NIC58-reference-based genetic relatedness into its principal components. We observed strong structure, with the top axis differentiating three African isolates from all others and accounting for almost 75% of the nuclear genetic covariances (Figure 5). The close clustering of the three African lines, isolated across a transect spanning more than 9000 km from the Atlantic island of Cape Verde, off the Westernmost coast of continental Africa, to Réunion Island in the Indian Ocean, is remarkable given the large geographic distances separating these locales, though it is consistent with the occurrence of globally distributed haplotypes in *C. elegans* (Lee et al., 2020). PC2 differentiates Western Pacific samples (Hawaii, Taiwan) from all others, and PC3 largely differentiates two of four Taiwanese samples from Hawaii. These three dimensions account for 89% of the variance, which is of similar magnitude to the variance explained by the first three PCs in *C. briggsae* (based on 449,216 SNVs with no missing data among 34 lines). The genome-wide view masks heterogeneity at the chromosome level. Notably, chromosome IV shows more complex patterns of relatedness, and both chromosome V and the mitochondrial genome provide evidence of recent admixture, with strain QG834 from Panama clustering with strains from East Asia and the Pacific. Structuring of the X is particularly extreme, with essentially two haplotypes present in our sample – African and non-African – with this split accounting for 98% of genetic variation (Figure 5 – figure supplement 1 and 2).

While the population structure revealed by this analysis explains most of the genetic variance, we also observe variance within these groupings at divergent regions (Figure 5C,D). Examining local ancestry along chromosomes as the correlation between genome-wide SNV relatedness and that in a genomic window (after Stankowski et al. (2019)), we found that *C. briggsae* shows generally strong concordance, the much larger sample of *C. elegans* shows moderate concordance with higher variance, and *C. tropicalis* shows a particularly erratic relationship (variance around the median is 0.015, 0.063 and 0.074 for *C. briggsae, C. elegans,* and *C. tropicalis,* respectively; Figure 5C). While this statistic is expected to be sensitive to sample size and structure, the very different patterns seen for *C. briggsae* and *C. elegans* show that the results are not necessarily driven by these confounders. Genetic diversity and local discordance are strongly correlated across the three selfers genome-wide (Spearman’s ρ = 0.34, 0.37, 0.40 as ordered above, all p < 10^ࢤ25^). But the relationship across species diverges, near monotonically, with genetic divergence (e.g., **p** = 0.49, 0.04, 0.27, p < 10^ࢤ6^, p = 0.68, p = 0.015, above the 90^th^ divergence percentile for each species), consistent with differential structuring of divergent haplotypes among them (Lee et al., 2020). Looking across the *C. tropicalis* populations defined by genome-wide similarity, we see that within- and between-population genetic diversity are positively correlated in all pairwise comparisons, driven by a minority of genomic regions (Figure 5D). We also find a handful of highly divergent regions that vary only within populations, particularly Africa and the Americas. This includes loci on chromosomes IV and V that vary among seven isolates from a single collection on the small island nation of Dominica. The structure of *C. tropicalis* populations therefore mirrors that within genomes; featuring to a more extreme degree the strong differentiation seen between *C. briggsae* clades, the widespread homogeneity seen in *C. elegans* outside the Pacific, and the diversity seen at the local scale for both (Andersen et al., 2012; Barriere & Félix, 2007; Félix et al., 2013; Haber et al., 2005; Sivasundar & Hey, 2005).

### Quantitative genetics of outcrossing

We selected JU1373 and NIC58 because of their dramatic difference in outcrossing rate, as judged by single-worm and bulk-passaging assays. As a first step toward understanding the genetic basis of variation in outcrossing, we scored hermaphrodites from 118 RILs for their probability of producing cross offspring in matings with NIC58 males. We observed considerable variation among the RILs, including transgressive segregation (Figure 6A). Linkage mapping detected a significant effect of a locus on the center of the X chromosome (Figure 6B), which explained close to 15% of the variance in hermaphrodite outcrossing probability. Although the difference in equilibrium male frequency between JU1373 and NIC58 is likely mediated by their different outcrossing rates, we also observed differences in the rate of spontaneous male production due to X nondisjunction during hermaphrodite meiosis. Self progeny of NIC58 hermaphrodites were 0.8% male (21/2580), versus 0.06% in JU1373 (2/3088); given a characteristic brood size of 100–150, these numbers imply that most NIC58 self broods include a male, and most JU1373 self broods do not. These two strains thus differ heritably in male crossing ability, hermaphrodite crossing ability, equilibrium sex ratio, and spontaneous male production rate, providing multiple paths for the evolution of outcrossing rate.

**Figure 6.**
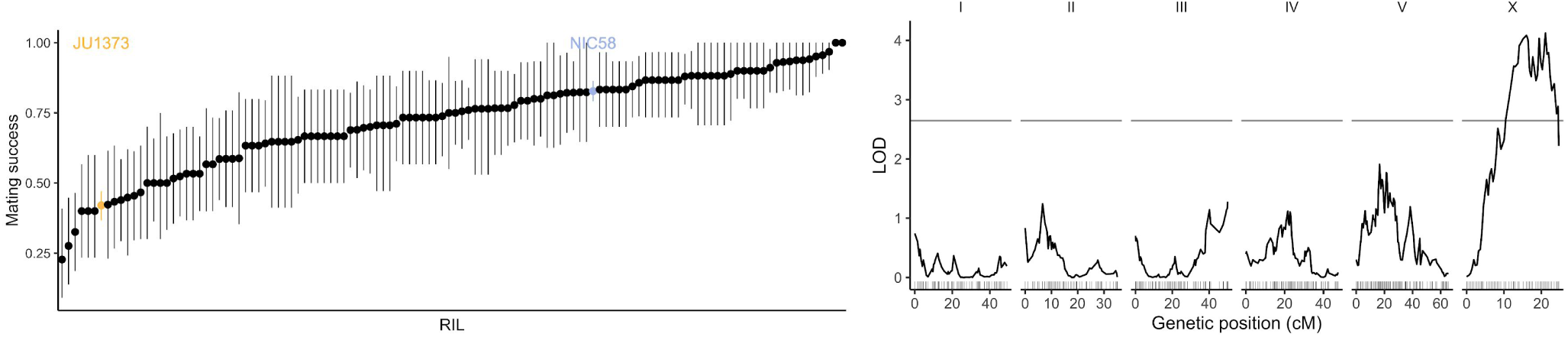
(**A**) RILs varied in their hermaphrodite crossing probability. Means and 95% bootstrap confidence intervals are shown for RILs and their parents. Data are in Figure 6 – source data 1. (**B**) Quantitative trait locus mapping for hermaphrodite crossing probability (genome-wide 0.05 significance threshold from 1000 phenotype permutations shown in grey, n=118 RILs). Data are based on Supplementary File 1.

### RIL segregation distortion and excess heterozygosity

The RIL genotypes displayed strong segregation distortion in two genomic regions: the left arm of chromosome III and the right arm of chromosome V (Figure 7). Both regions were strongly skewed toward JU1373 homozygotes, which reached a frequency of 97% on chromosome III and 78% on chromosome V. The chromosome V locus also showed an excess of heterozygotes (16% of RILs) compared to the theoretical expectation. RILs that retained heterozygosity on chromosome V also showed an enrichment of JU1373 genotypes on chromosome I (18.2–20.7 cM), which itself showed mild distortion in favor JU1373 (Fisher’s exact test p < 0.001, Figure 7 – source data 1). These data indicate that selection during RIL construction strongly favored the JU1373 allele chromosomes III and V, with complex selection at the locus on chromosome V favoring heterozygotes over JU1373 homozygotes under some conditions.

**Figure 7.**
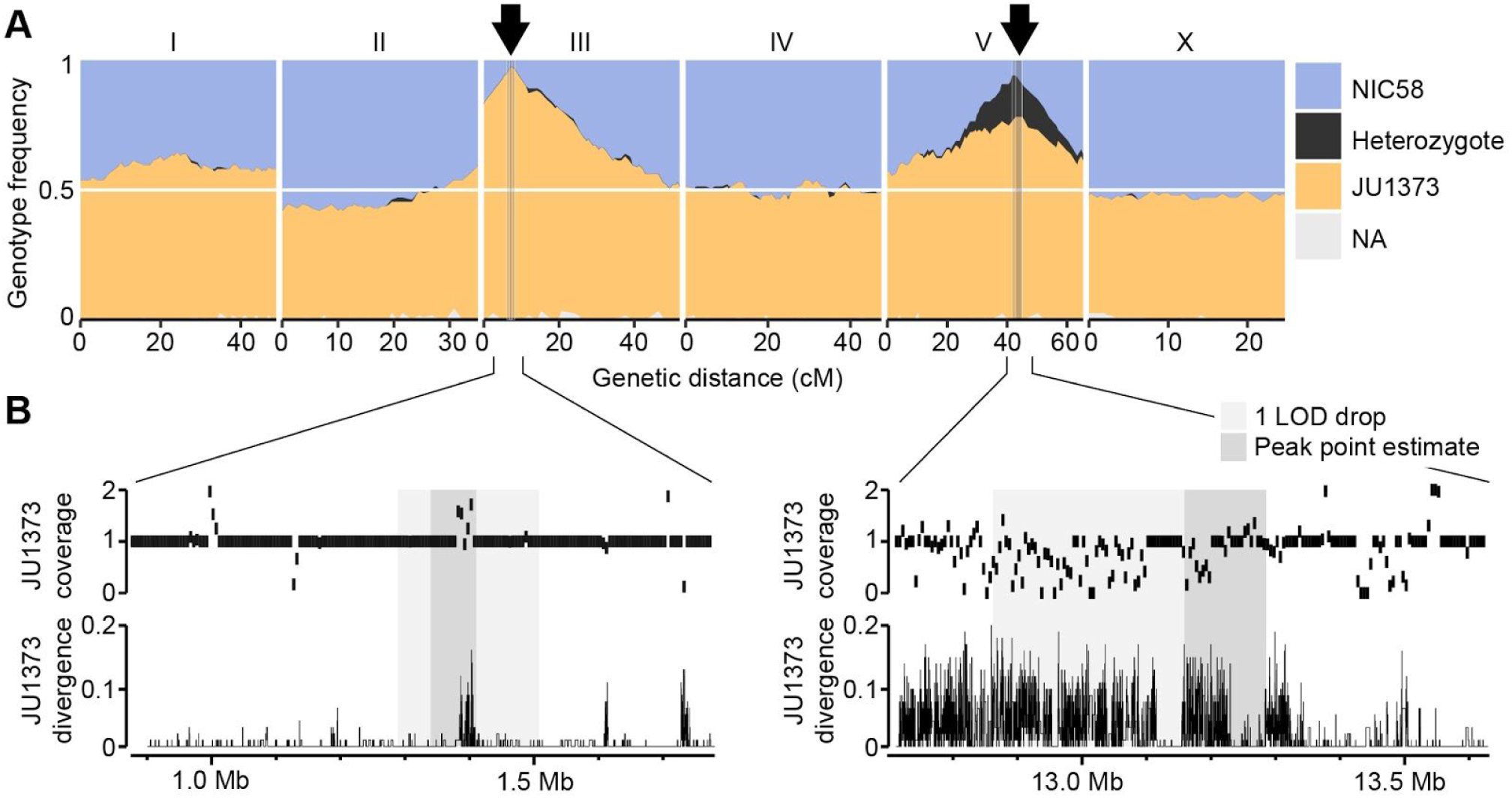
Two genomic regions show strong segregation distortion among RILs. (**A**) RIL genotype frequencies. Arrows are peaks of segregation distortion. Shaded areas are 1 LOD drop intervals and peak point estimates. Genome-wide data are based on Supplementary File 1, and multilocus segregation distortion genotype tables are in Figure 7 – source data 1. (**B**) Fold coverage and SNV divergence of JU1373 relative to NIC58. Fold coverage is in 5 kb windows, divergence is SNV identity in 100 bp windows. Data are based on Figure 4 – source data 2.

### Segregation distortion is not due to mitochondrial-nuclear incompatibilities

Strong selection during RIL construction is consistent with previous reports of extensive outbreeding depression (Gimond et al., 2013). Yet simple genetic incompatibilities between two nuclear loci are not expected to favor one parental allele to the exclusion of the other. Exclusion of one parental allele can occur, however, if an allele from the male parent (NIC58) is incompatible with the mitochondrial genome of the hermaphrodite parent (JU1373). Under this scenario, RILs homozygous for the male parent allele (NIC58) at loci showing segregation distortion should be sub-viable or sub-fertile. We examined RILs of such genotypes and found that their growth characteristics were superficially normal, with 93.1–98.3% (n = 151–679) of embryos developing into adults with parental developmental timing (Figure 9A). This pattern shows that mitochondrial-nuclear incompatibilities were not the cause of segregation distortion among RILs.

### Segregation distortion and excess heterozygosity are caused by drive loci

An alternative explanation for segregation distortion among RILs is that the distorted loci independently experienced drive-like dynamics, similar to those seen in *C. elegans* at the *zeel-1/peel-1* and *pha-1/sup-35* loci (Ben-David et al., 2017; Seidel et al., 2008). At these loci, a maternal- or paternal-effect locus loads a toxin into eggs or sperm that poisons zygotic development; subsequent zygotic expression from the same locus provides an antidote (Ben-David et al., 2017; Seidel et al., 2011). This model predicts that the JU1373 genome encodes two independent driver alleles, each encoding a toxin-antidote pair (Figure 7 – figure supplement 1). Under this model, all F_2_ progeny from a NIC58 x JU1373 cross will be exposed to toxins, but only some will inherit antidotes; animals not inheriting both antidotes will suffer the effects, manifesting as embryonic or larval arrest, sterility, or some other phenotype that would have prevented them from contributing to the RILs. The proportion of F_2_s showing such phenotypes is expected to be ~44% (7/16), assuming that arrest by each toxin is fully penetrant, and that arrest can be rescued by a single copy of the corresponding antidote.

**Figure 7- figure supplement 1.**
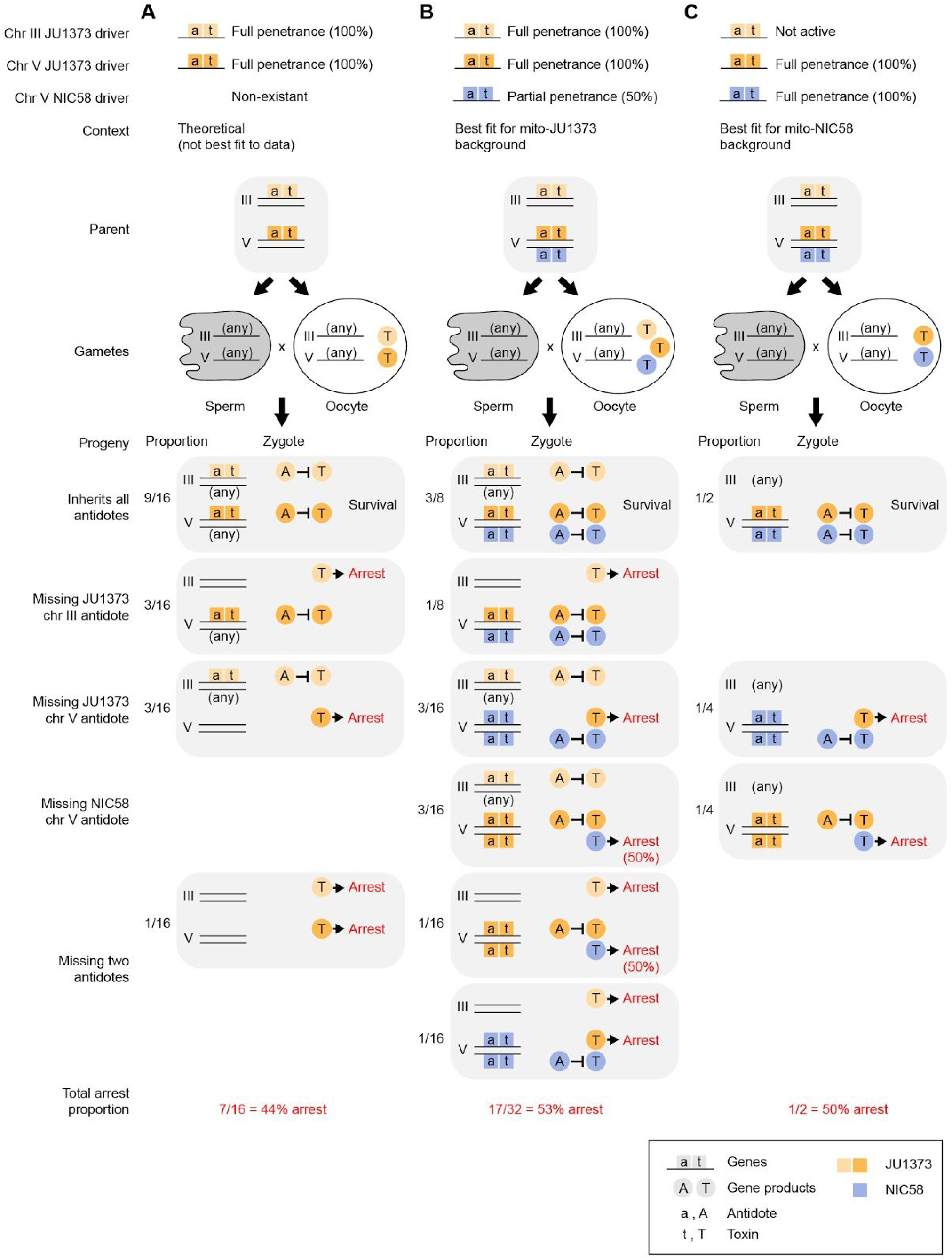
Models of gene drive and expected F_2_ arrest proportions. Models assume that each driver is composed of two genes, a maternally expressed toxin and zygotically expressed antidote. (**A**) Two JU1373 drivers, one on chromosome III and one on chromosome V. This model is not the best fit for observed genotype frequencies among wild-type F_2_ animals. (**B**) A JU1373 driver on III and antagonistic drivers (JU1373 and NIC58) on V. The NIC58 driver is assumed to be 50% penetrant, true penetrance likely depends on genetic background. This model is the best fit for observed genotype frequencies among wild-type F_2_ animals in a mito-JU1373 background. (**C**) Inactive JU1373 driver on III and antagonistic drivers (JU1373 and NIC58) on V. This model is the best fit for observed genotype frequencies among wild-type F_2_ animals in a mito-NIC58 background.

To test whether F_2_ populations from a NIC58 x JU1373 cross showed phenotypes consistent with two driver loci, we made reciprocal crosses, allowed F_1_ hermaphrodites to self-fertilize, and followed F_2_ progeny from embryo to adulthood. We observed that only 41–45% (n = 329–1283) of F_2_ embryos developed into adults within the normal developmental time, versus 98–99% (n = 1046–1093) for parental strains. Terminal phenotypes among abnormal F_2_ animals included failure to hatch (9%), early larval arrest (39–41%), late larval arrest (5–9%), and abnormally small, thin adults (5%; n = 329–381); these phenotype frequencies did not differ according to the direction of the cross that generated the F_1_ worm (Fisher’s exact test, p = 0.39). These data show that F_2_ populations experienced widespread developmental arrest, at proportions slightly higher than expected under a model of two driver loci. One interpretation is that the total arrest proportion among F_2_ individuals reflects the effects of two driver loci (~44% of F_2_ animals), plus additional background incompatibilities (~12–15% of F_2_ animals).

The two-driver model predicts that developmental arrest among F_2_ animals will preferentially affect animals homozygous for non-driver (NIC58) alleles. To test this prediction, we repeated reciprocal NIC58 x JU1373 crosses and genotyped F_2_ progeny at markers tightly linked to the segregation distortion peaks on chromosomes III and V. We observed that alleles at both loci were transmitted to progeny in Mendelian proportions, showing that meiosis and fertilization are not affected by the drive loci. Certain genotypes were associated with developmental arrest, but these differed depending on the direction of the cross (i.e., mitochondrial genotype). For the chromosome III locus, NIC58 homozygotes underwent complete developmental arrest in the mito-JU1373 cross but very little developmental arrest, compared to other genotypes, in the mito-NIC58 cross (Figure 8A). For the chromosome V locus, NIC58 homozygotes experienced highly penetrant developmental arrest in both crosses (Figure 8A). Chromosome V JU1373 homozygotes also experienced developmental arrest in both crosses, but with a lower penetrance, especially in the mito-JU1373 cross (Figure 8A). This pattern supports a model of two drive loci, but shows that the drive loci interact with mitochondrial genotype: for the locus on chromosome III, the driver allele (JU1373) is active in its own mitochondrial background but inactive or very weakly active in the opposite mitochondrial background; for the locus on chromosome V, both alleles (JU1373 and NIC58) act as drivers and are effectively antagonistic – JU1373 acts as a strong driver in both mitochondrial backgrounds, whereas NIC58 acts as a strong driver in its own mitochondrial background and a weaker driver in the opposite mitochondrial background (Figure 8C). Importantly, this finding of antagonistic drive at the chromosome V locus provides a simple explanation for the retention of heterozygosity at this locus among the RILs (Figure 7).

**Figure 8.**
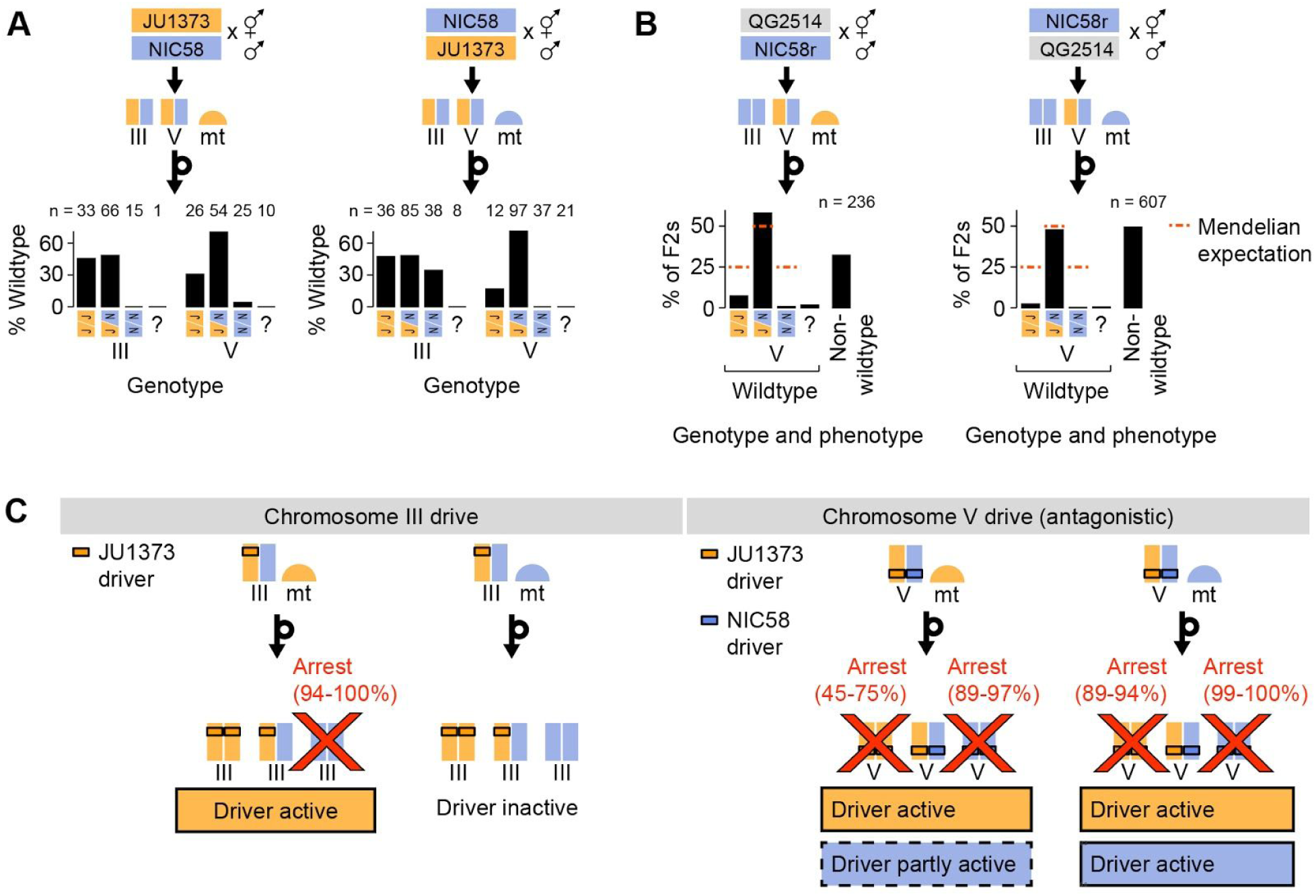
Drive genetics. (**A**) Percentage of F_2_ progeny from reciprocal NIC58 x JU1373 crosses showing wild-type development. Genotypes reflect markers tightly linked to the peaks of segregation distortion on chromosomes III and V. Question mark, genotyping failure. (**B**) Genotype and phenotype frequencies among F_2_ progeny from reciprocal crosses between NIC58 and RIL QG2514. Only wild-type F_2_ progeny were genotyped. Data are in Figure 8 – source data 1. (**C**) Schematic of drive activity for the loci on chromosomes III and V. Percentages are estimates for the proportion of animals undergoing developmental arrest, compared to heterozygous siblings. Estimates were derived by comparing observed genotype frequencies among wild-type F_2_ progeny to Mendelian expectations. This method avoids bias introduced by genotyping failures being more common among arrested versus wild-type animals. The following reciprocal crosses were used to estimate arrest proportions: NIC58 x JU1373, NIC58 x RIL QG2479 (not shown), and NIC58rx RIL QG2514. NIC58r expresses a red fluorescent transgene (see Methods).

**Figure 9.**
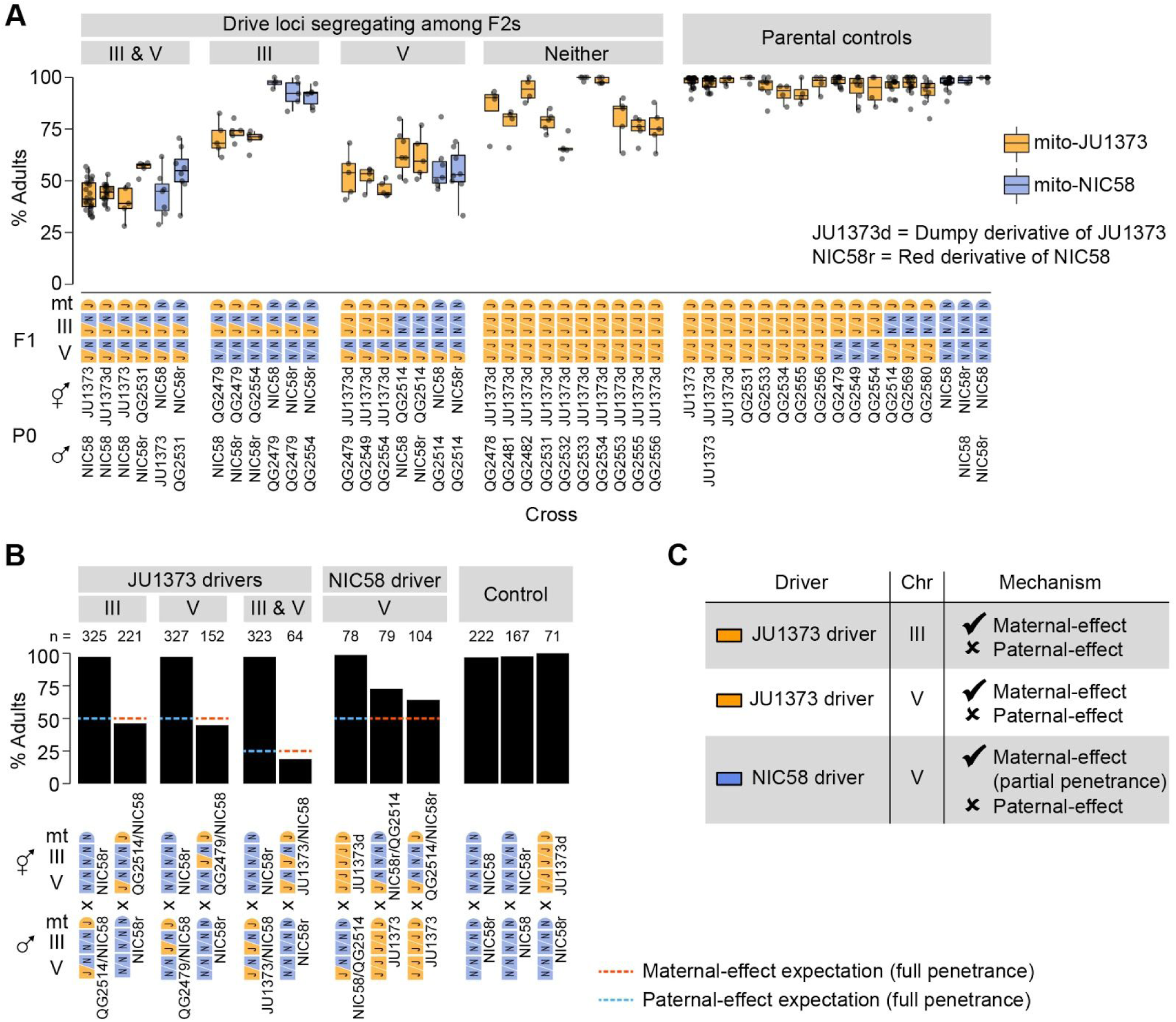
Drive loci act independently and by maternal effect. (A-B) Percentages of F_2_ or backcross progeny that reached adulthood within 72 hours of egg laying. Strains beginning with “QG” are RILs. JU1373d is a Dumpy mutant. NIC58r expresses a red fluorescent transgene (see Methods. Each point is a cross plate, with a median of 80 worms scored per plate. Data in Figure 9 – source data 1). (**A**) Crosses testing whether drive loci act independently. (**B**) Crosses testing whether drive loci act via maternal or paternal effect. Maternal- and paternal-effect expectations are under a model that either a maternal- or paternal-effect toxin causes fully penetrant developmental arrest for progeny not inheriting the driver haplotype. (**C**) Interpretation of maternal- and paternal-effect crosses. Partial penetrance of the NIC58 driver is consistent with this driver showing partial penetrance among F_2_ progeny.

**Figure 9- figure supplement 1.**
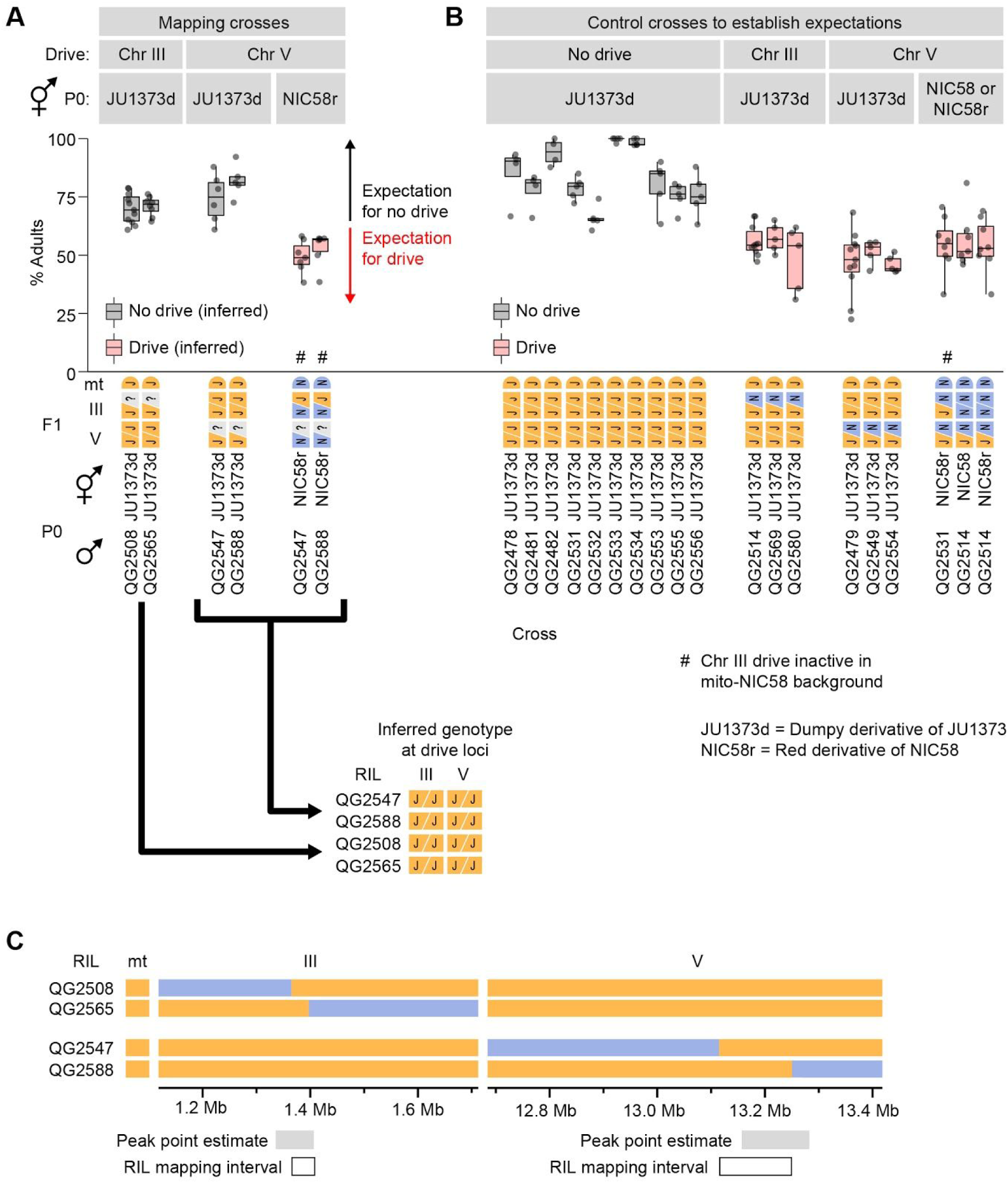
Mapping drive loci using RILs. (A-B) Percentages of F_2_ progeny that reached adulthood within 72 hours of egg laying. Strains beginning with “QG” are RILs. JU1373d is a Dumpy mutant. NIC58r expresses a red fluorescent transgene (see Methods, Data in Figure 9 – source data 1). (**A**) Crosses to determine whether RILs with recombination breakpoints near drive loci exhibit drive activity in crosses to JU1373d and NIC58r. (**B**) Crosses to establish expectations for mapping crosses. Some of the data here is duplicated from Figure 10 for ease of comparison. (**C**) RIL genotypes and intervals to which drivers were mapped.

### Antagonistic drive does not reflect mitochondrial-nuclear incompatibility

The reciprocal crosses described above showed that developmental arrest of chromosome V JU1373 homozygotes was more penetrant in a mito-NIC58 background than in a mito-JU1373 background. One possible contributor to this arrest is mitochondrial-nuclear incompatibility between the JU1373 haplotype of the chromosome V locus and the mitochondrial genome of NIC58; alternatively, the NIC58 driver on chromosome V may be more active in one mitochondrial background compared to the other. To test for mitochondrial-nuclear incompatibility, we attempted to recover lines homozygous for the JU1373 driver in a mito-NIC58 background. These lines were recovered by crossing males of RIL QG2514 to NIC58, producing an F_2_ population segregating at the chromosome V locus but fixed for the NIC58 haplotype at the chromosome III locus (Figure 8B). We recovered 10 F_2_ animals homozygous for the JU1373 V allele in a mito-NIC58 background. These animals produced progeny that were superficially wild-type, with typical brood sizes and developmental rates. This finding excludes mitochondrial-nuclear incompatibility as a contributor to developmental arrest of chromosome V JU1373 homozygotes. We conclude that differences in developmental arrest according to mitochondrial genotype reflect the NIC58 driver on chromosome V being more active in a mito-NIC58 background than in a mito-JU1373 background. We also conclude that activity of the NIC58 driver in the mito-JU1373 background is likely dependent on additional nuclear background factors, thus accounting for heterozygosity in the RILs being preferentially retained in certain genetic backgrounds (e.g., in animals with JU1373 alleles on the left arm of chromosome I).

### Drive loci act independently

Our model predicts that the drive loci act independently and are thus genetically separable. To test this prediction, we examined RILs with opposite parental genotypes at the two loci. Each RIL was crossed to an appropriate parental strain to generate F_2_ animals segregating opposite alleles at one drive locus but fixed at the other, which we scored for development. To provide a control for (non-drive) background incompatibilities, we repeated this analysis for F_2_ populations fixed for JU1373 alleles at both drive loci. These were generated by crossing 10 RILs carrying JU1373 alleles at both drive loci to a JU1373-derived strain carrying a Dumpy mutation, which allows us to distinguish cross progeny from self progeny, a constant issue for JU1373 with its low rate of crossing by hermaphrodites (Figure 1).

We observed that crosses segregating only the chromosome III driver in a mito-JU1373 background showed median rates of normal development consistent with drive activity at a single locus (68–75%, n = 218–240); crosses in a mito-NIC58 background showed no drive activity (Figure 9A). The arrested progeny in the mito-JU1373 background largely consisted of chromosome III NIC58 homozygotes, as evidenced by these animals being severely depleted among wild-type progeny (26:64:2 JJ:JN:NN genotypes), while the wild-type progeny of the reciprocal cross carried chromosome III genotypes at their expected Mendelian proportions (30:57:31). These data show that the chromosome III driver is active in the absence of drive at the chromosome V locus, and confirm that the chromosome III driver requires a mito-JU1373 genetic background.

Crosses segregating only at the chromosome V locus showed median rates of normal development that were similar across mitochondrial backgrounds and consistent with antagonistic drive at a single locus (54–61%, n = 203–321, in a mito-JU1373 background; 52–53%, n = 313–345, in a mito-NIC58 background; Figure 9A). Arrested progeny largely consisted of NIC58 homozygotes and, to a lesser extent, JU1373 homozygotes, as evidenced by these genotypes being depleted among wild-type progeny (Figure 9B). This result shows that antagonistic drive at the chromosome V locus occurs in the absence of drive at the chromosome III locus. For comparison, crosses segregating no drivers showed median rates of normal development that were usually higher than ~75%, as expected for crosses lacking drive activity (Figure 9A); nonetheless, rates of normal development in these crosses were highly variable and sometimes low, with medians ranging from 65–100% (n = 163–213), suggesting a contribution from non-drive background incompatibilities. We conclude that the chromosome III and V drive loci act independently of one other, and that additional background incompatibilities are widespread and diffuse in the NIC58 x JU1373 cross.

### Drive loci act via maternal effect

The drive loci we identified in *C. tropicalis* produce segregation distortion via an interaction between parent and offspring genotypes. To distinguish between maternal and paternal effects, we scored rates of normal development among progeny in which driver alleles segregated from mothers or fathers. For tests of JU1373 drivers, progeny were derived from reciprocal crosses between a derivative of NIC58 expressing red fluorescent protein from an integrated transgene and animals heterozygous for the drivers. For tests of the NIC58 driver, progeny were derived from reciprocal crosses between JU1373 (or a Dumpy derivative of JU1373) and animals heterozygous for the driver. For a maternal effect (and no paternal effect), ~50% of cross-progeny lacking the driver were expected to undergo developmental arrest when the driver was present in the mother, assuming full penetrance, but develop normally when the driver was present in the father. In the case of a paternal effect (and no maternal effect), the opposite pattern was expected. Crosses testing the JU1373 drivers showed that both acted via maternal, and not paternal, effect (Figure 9B). Crosses testing the NIC58 driver ruled out paternal effect (Figure 9B); crosses testing a maternal effect were confounded by the poor mating ability of JU1373 males, but were consistent with a maternal effect (Figure 9B). We therefore infer a maternal effect for the NIC58 driver, and conclude that all three driver alleles act via maternal, and not paternal, effect (Figure 9C). The overall model is presented in Figures 9C and Figure 7 – figure supplement 1.

### Incompatibilities are associated with duplicated and novel genes

To understand the genetic basis for drive activity, we mapped the JU1373 drivers using RILs with recombination breakpoints near the peaks of segregation distortion. RILs were crossed to derivatives of parental strains, and drive activity was assessed by scoring the rate of normal development among F_2_ progeny. This analysis mapped drivers to intervals of around 33 kb on chromosome III and 69 kb on chromosome V (Figure 9 – figure supplement 1) that overlapped the peaks of segregation distortion (Figure 7). The chromosome III interval encompasses a locus of locally elevated sequence divergence between NIC58 and JU1373, in an otherwise well-conserved region. A clear driver candidate is an insertion of sequence in JU1373 that includes seven predicted genes (Figure 10A): six are tandem duplications of neighboring genes in NIC58; the seventh is a duplication of a gene located 0.68 Mb to the right in both NIC58 and JU1373. Five of the seven genes have no detectable protein or nucleotide homology outside *C. tropicalis.* The remaining two are homologous to *C. elegans* genes *F44E2.8* and *F40F8.11,* which share NADAR and YbiA-like superfamily protein domains. In addition to these seven genes, JU1373 also carries an eighth gene inserted within the original copy of the duplicated sequence (Figure 10A, arrowhead). This gene is novel and has no detectable homology to any gene in NIC58 or in any other species. Thus, the JU1373 driver on chromosome III contains a total of eight additional genes compared to NIC58: one unique to JU1373, and five with no homology outside *C. tropicalis.*

**Figure 10.**
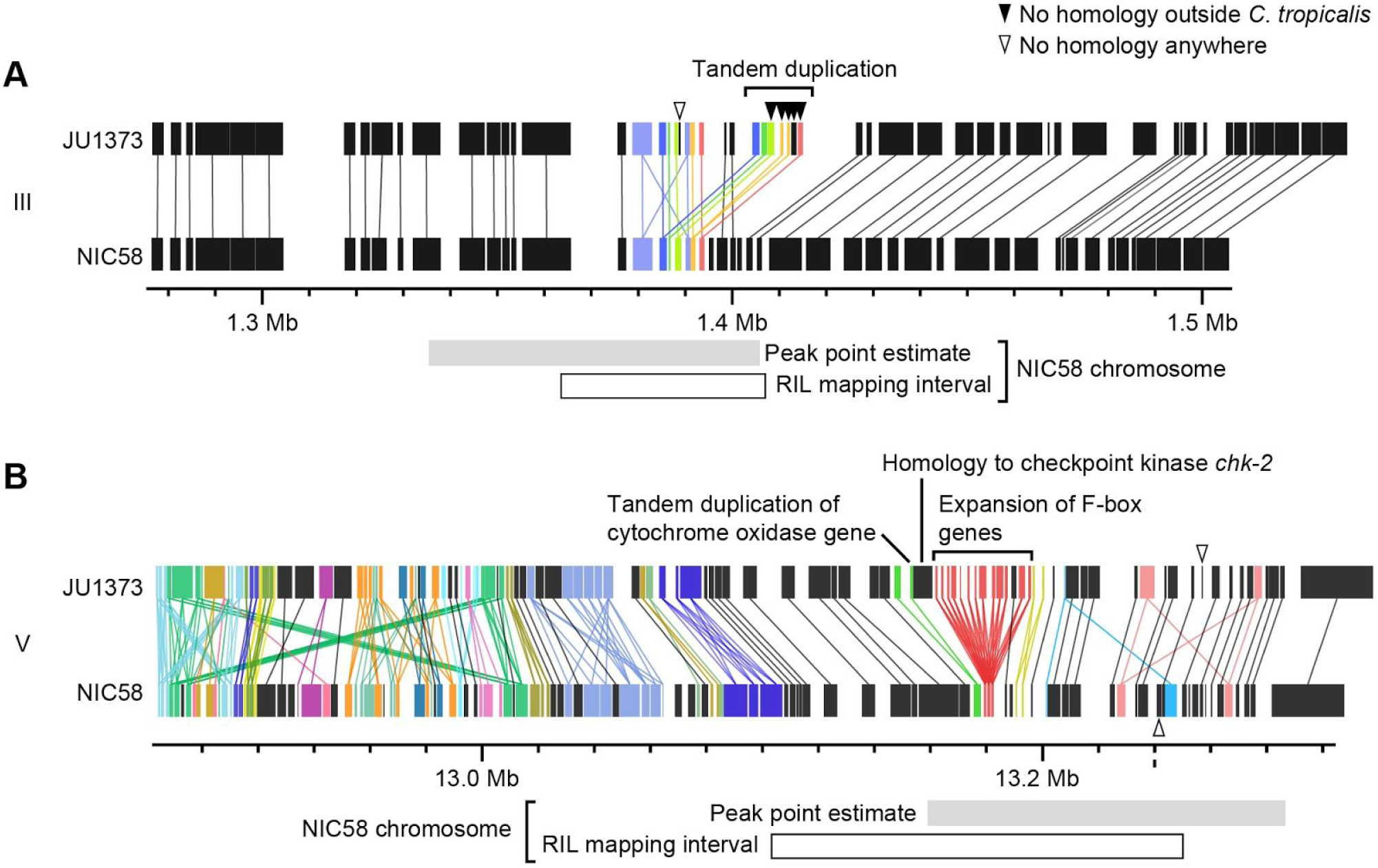
Genomic internals surrounding drive loci on chromosome III (**A**) and V (**B**). Windows span segregation distortion 1 LOD drop intervals. Rectangles, predicted genes. Lines connect homologs. Colors indicate the union of homologs within the interval. Homology relationships to genes outside the intervals are not shown. Data are based on Supplementary File 2 and Supplementary File 6.

**Figure 10- figure supplement 1.**
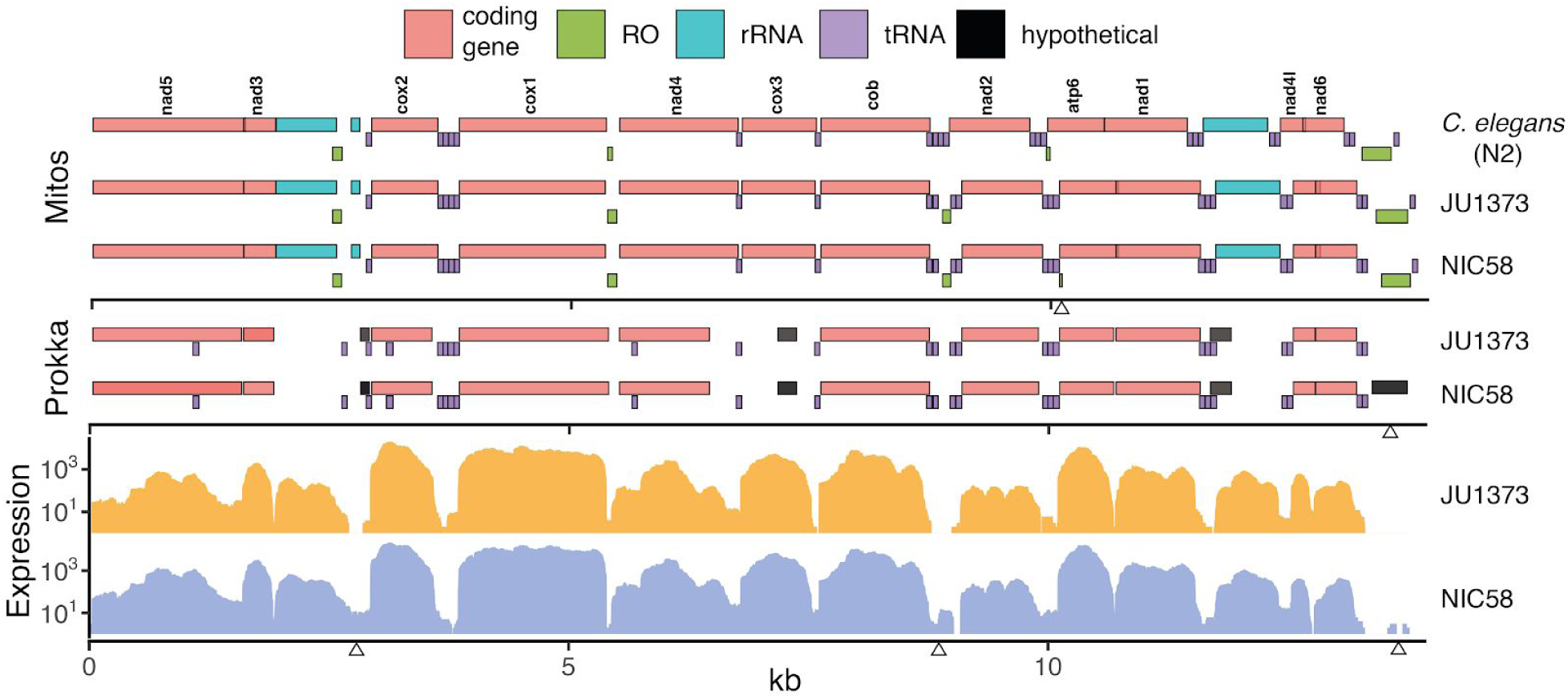
Mitochondrial genomes. Annotations are shown for two homology-based methods, Mitos (Bernt et al., 2013) and Prokka (Seemann, 2014), over expression from short-read RNAseq data (per base read depth). RO: potential replication origin, including the large D-loop non-coding region at 13.5 kb. Some obvious differences of unknown significance are highlighted with arrowheads along the three x-axes including (top to bottom, left to right): the presence of a small, low-scoring RO in NIC58 called by Mitos; the presence of a novel open reading frame, called by Prokka, overlapping the D-loop in NIC58 and predicted to encode a 122-amino acid transmembrane protein; and three regions of clear differential expression overlapping a 16S ribosomal RNA fragment, a tRNA and RO cluster, and the D-loop region. Data are based on Supplementary File 2.

The chromosome V locus lies in a region of high divergence between NIC58 and JU1373, extending well beyond the mapped interval (Figure 7B). This region is home to a number of dynamically evolving gene families (Figure 10B). A major structural difference between JU1373 and NIC58 haplotypes is an expansion of divergent F-box-domain-encoding genes, expanded from three homologs in NIC58 to 13 in JU1373. Immediately flanking this expansion in JU1373 is a duplication of a gene located 0.6 Mb away (and present in both NIC58 and JU1373), which encodes a homolog of the checkpoint kinase *chk−2.* Adjacent to the *chk−2* homolog is a tandem duplication of a nuclear gene encoding a mitochondrial ubiquinol-cytochrome c oxidoreductase complex protein, perhaps accounting for the interaction of drive activity with mitochondrial genotype. Additional differences between haplotypes include a gene encoding a small and novel protein in JU1373 and a novel gene in NIC58. We also examined the mitochondrial genomes of JU1373 and NIC58, and found that the core functional complement is superficially identical; all 12 protein-coding, 2 ribosomal RNA and 22 tRNA genes are called as present and intact in both (Bernt et al., 2013; Lemire, 2005; R. Li et al., 2018). There are, however, several differences of unknown significance, including 12 missense variants in six of the core protein coding genes, the presence in NIC58 of a small non-coding region and a potentially novel (lowly expressed) gene encoding a 2-pass transmembrane protein, and differential expression at two additional loci (Figure 10 – figure supplement 1). Thus, the JU1373 driver on chromosome V resides within a larger genomic window of elevated sequence divergence, containing an expansion of F-box genes, a novel gene, as well as a duplicated gene encoding a mitochondrial protein. The inferred antagonistic driver in NIC58 may also involve novel protein-coding genes, and there are a number of candidate variants in mitochondrial genomes, including a potentially novel protein-coding gene, that might underlie nuclear-cytoplasmic interaction.

### Incompatibilities consistent with drive activity are widespread among wild isolates

To examine the distribution of putative drive activity among wild strains, we crossed 14 isolates to a Dumpy derivative of JU1373, to NIC58, or to both, and scored development among the F_2_. Crosses were classified as having putative drive activity if rates of normal development were below ~75% (the expectation for segregation of a single driver). We observed that crosses to JU1373 showed putative drive activity for 12 of 13 isolates, and the strength of drive activity was associated with geographic origin, and haplotype at the chromosome III and V drive loci (which are themselves associated due to population structure). The two African isolates showed the least drive activity and had haplotypes similar to JU1373 at both drive loci; five American isolates showed consistently higher drive activity and carried haplotypes dissimilar to JU1373 at both drive loci; isolates from other areas were more variable, but drive activity was lowest for isolates carrying haplotypes similar to JU1373 at the chromosome V locus (NIC535 and NIC773) or somewhat similar to JU1373 at both loci (QG131; Figure 11). Nonetheless, the correlation between drive activity, geographic origin, and haplotype was imperfect (e.g., JU1630 versus JU3170), and some isolates showed median rates of normal development consistent with segregation of three or more drivers (NIC517 and QG834, 20–28%, n= 257–276). This pattern suggests that the JU1373 drivers may be fixed within Africa but polymorphic or absent elsewhere, and that some of the non-African isolates may contribute additional drivers of their own.

**Figure 11.**
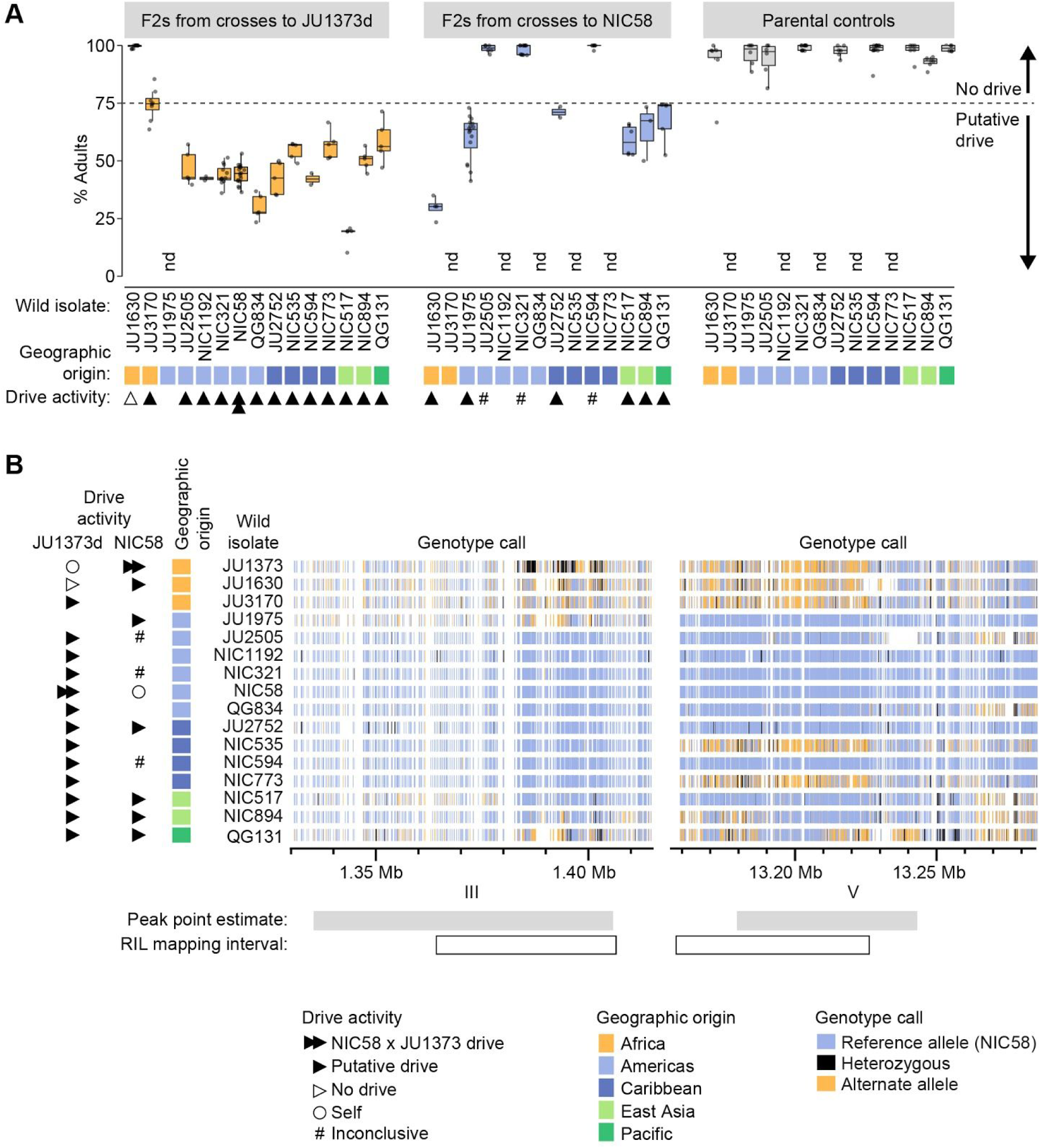
Wild isolate phenotypes and haplotypes. (**A**) Percent of F_2_ progeny reaching adulthood within the normal developmental time (∼72 hrs), for crosses between wild isolates and a Dumpy derivative of JU1373 (JU1373d) or NIC58. #, inconclusive because we cannot be certain that F_1_ animals were cross progeny, nd, not determined. Putative drive, crosses in which the median percent of F_2_ animals reaching adulthood was less than ∼75%. Each plotted point is a plate (2–16 per cross, median 6), with a median 90 animals scored per plate. Data in Figure 11 – source data 1. (**B**) Wld isolate haplotypes at the drive loci on chromosomes III and V. Heterozygous calls likely reflect duplication and divergence. Data are based on Supplementary File 4.

Among crosses to NIC58, putative drive activity was observed in six of nine crosses (Figure 11). Five of these crosses showed drive activity that was moderate in strength and largely overlapping, despite these isolates having different haplotype combinations at the chromosome III and chromosome V drive loci (Figure 11). The sixth cross (JU1630) showed much higher drive activity, consistent with this isolate alone having JU 1373-like haplotypes at both drive loci (Figure 11). The three remaining crosses showed no drive activity and had haplotypes similar to NIC58 at both loci; we conservatively interpret these crosses as inconclusive because we cannot be certain of paternity of F_1_ animals (crosses to NIC58 did not include a marker to distinguish cross- from self-progeny). Nonetheless, these data as a whole indicate that putative drive activity is widespread. Drivers sometimes segregate within a geographic region, and some crosses may segregate drivers in addition to those identified in the NIC58 x JU1373 cross.

### Drive dynamics in partial selfers

Populations segregating drive loci like those we discovered on chromosomes III and V are expected to evolve suppressors to alleviate the reduced fitness of animals whose progeny are killed by the toxins. A general resistance mechanism for suppressing this lethality is selfing, which reduces the prevalence of heterozygotes and hence of animals expressing drive-associated lethality. To better understand how the mating system of *C. tropicalis* influences the spread of maternal-effect toxin / zygotic-effect antidote drive elements, we simulated evolution under varying levels of selfing and outcrossing. These simulations include the idiosyncratic features of *Caenorhabditis* androdioecy, including the facts that hermaphrodites cannot mate with one another and that males can arise spontaneously by nondisjunction of the X chromosome (see Methods). We found that in a large population, high rates of selfing can dramatically slow the rate of spread of a drive element, and indeed can reduce its efficacy below the threshold required for selection to act on it (Figure 12A). In small populations, elements introduced at 5% frequency are rapidly fixed under random mating but lost to drift under high rates of selfing (Figure 12B). These simulations also reveal that once a driver is at high frequency, selfing actually hastens fixation of the driver (compare partial selfing at S = 0.25 to obligate outcrossing at S = 0). This occurs because once a driver is at high frequency, heterozygotes that self will always expose their progeny to the effects of drive, but heterozygotes that outcross will usually mate with an antidote-carrying male, slowing the fixation of the driver.

**Figure 12.**
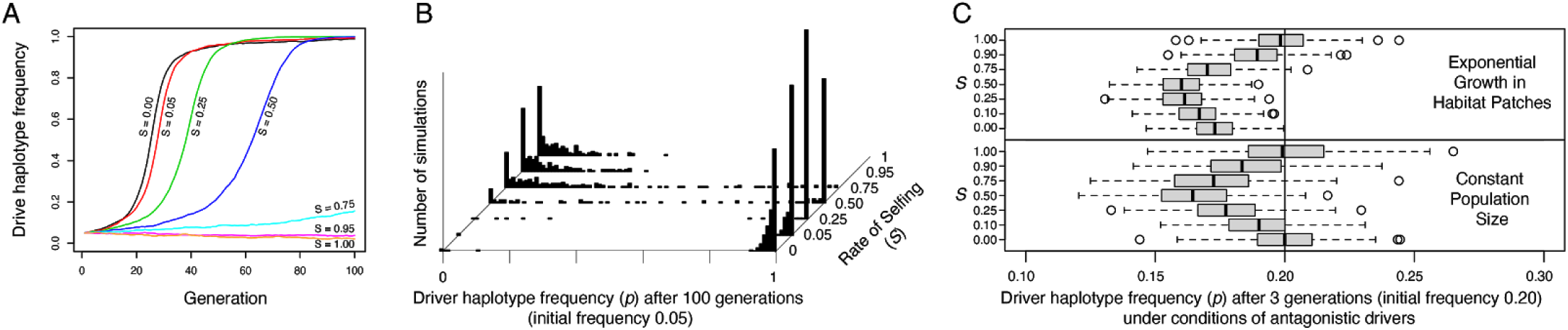
Selfing reduces the efficacy of maternal-toxin/zygotic-antidote drive elements. (**A**) Representative allele frequency trajectories of a drive haplotype under different rates of selfing (S). Population size is 20,000 in each case, the initial drive allele frequency is 0.05, and initial genotype frequencies and sex ratios are those expected at neutral equilibrium given the selfing rate. (**B**) Distribution of drive allele frequencies after 100 generations in populations of size 1000. Each histogram shows the outcome of 250 simulations with initial drive allele frequency 0.05. Drive alleles are often lost under high selfing rates. (**C**) Antagonistic drivers induce positive frequency dependent selection, acting against the rarer driver, when selfing rates are intermediate, when populations undergo exponential growth in ephemeral habitat patches, or both. Each boxplot represents the results of 250 simulations of three generations of evolution starting from allele frequency 0.2, with initial population size 1000. In the patchy environment, those 1000 individuals are distributed among 250 separate patches, and population growth is unbounded within each. Source code is available from [github](https://github.com/lukemn/tropicalis/tree/master/driver_simulations).

**Figure 12- figure supplement 1.**
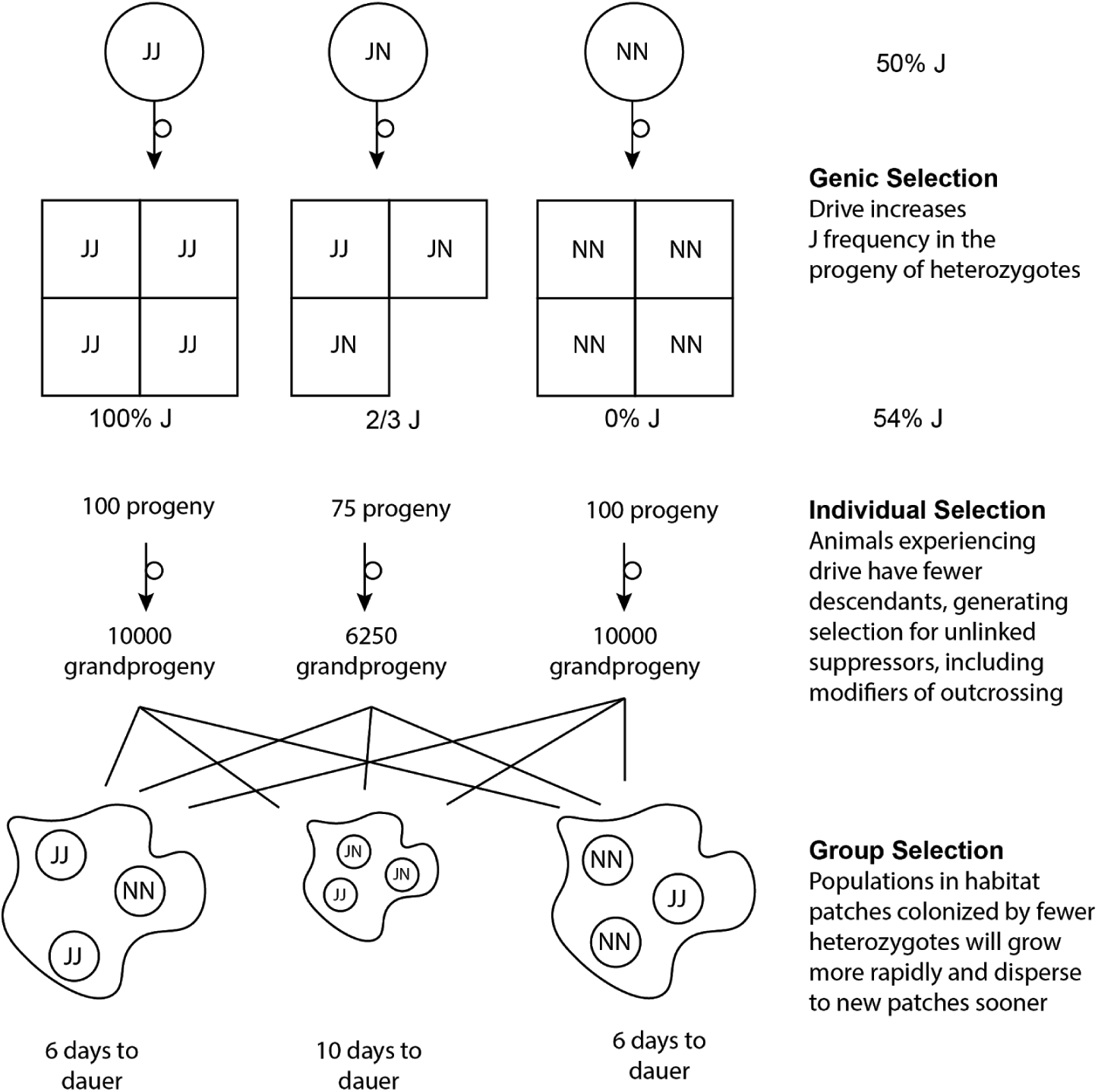
In a single-driver scenario, genic and group selection affect the frequencies of drive haplotypes, and drive-induced deaths generate individual-level selection for drive suppression.

**Figure 12- figure supplement 2.**
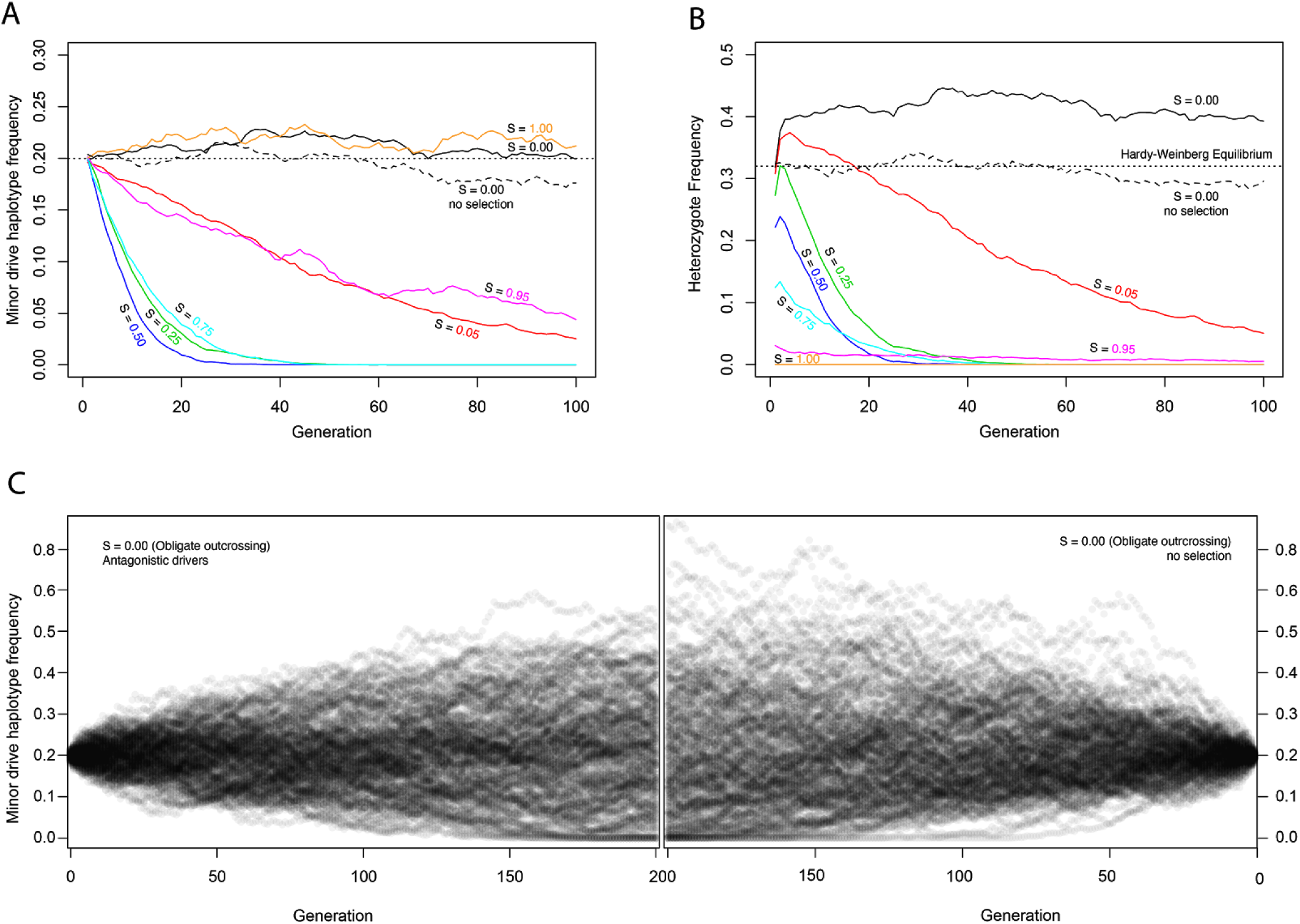
Antagonistic drivers are subject to drift or positive frequency-dependent selection, depending on the selfing rate. (**A**) At a locus with two haplotypes, each containing a maternal-effect toxin gene and a zygotic-effect antidote gene, allele frequencies change overtime as a function of selfing rate (S). At intermediate selfing rates, positive frequency-dependent selection acts to remove the rarer haplotype from the population. This effect is explained by the disproportionate impact of segregation in heterozygous selfers on the rarer haplotype. Heterozygous selfers kill each haplotype in equal proportions (% of progeny are homozygous for each haplotype), while outcrossers kill each as a function of their population frequency; the result is that the rare allele suffers more from heterozygote selfing. Intermediate levels of outcrossing result in the highest frequency of heterozygous selfer and so the most efficient selection against the rare allele. Under complete selfing (S = 1), there are no heterozygotes and the drivers have no effect. Under obligate outcrossing, the haplotypes are subject to drift and show similar dynamics to obligate selfers or outcrossers without drivers. The pattern in obligate outcrossers reflects the maternal-zygotic interaction character of the drivers. While the drivers kill homozygotes, generating overdominance and heterozygote excess, the drivers only have this effect in heterozygous mothers. The fitness cost to these heterozygous mothers represents underdominance, favoring homozygotes, and the over- and under-dominance balance each other. This figure shows the results of representative simulations with fixed population size of 20,000 breeding individuals, initial frequency 0.2, and 95% penetrance of each of the two drivers. (**B**). Heterozygote frequencies (from the simulations shown in A) vary over evolution. Obligate outcrossers maintain substantially elevated heterozygote frequencies. (**C**) Surprisingly, despite the strong selection against homozygotes and the elevated frequency of heterozygotes in obligate outcrossers, allele frequencies evolve with drift-like dynamics. Results of 200 simulations are shown, each with a fixed population size of 1,000 breeding individuals and initial frequency of 0.2. The left panel illustrates the spread of allele frequencies across simulations in the case of antagonistic drivers, and the right shows results with no drivers. In both cases, frequencies drift and some populations lose the lower-frequency allele. After 200 generations, the allele frequency variances are not different between the two cases (Levene’s test, p = 0.13). Source code is available from [github](https://github.com/lukemn/tropicalis/tree/master/driver_simulations).

We next considered antagonistic drivers, under the simplistic scenario of perfect linkage and equal penetrance of the two drivers. We found that intermediate levels of selfing cause frequency-dependent selection against the minor haplotype (Figure 12C, lower panel, and Figure 12 – figure supplement 2). Surprisingly, allele frequencies of the antagonistic drivers drift as though neutral under both obligate selfing and obligate outcrossing, albeit for different reasons. With obligate selfing, there are simply no heterozygotes and the alleles are literally neutral. Under obligate outcrossing, the killing of both homozygote classes generates overdominance, but the costs of these deaths are borne by heterozygote mothers, generating offsetting underdominance, and the net effect is that the alleles experience drift despite the enormous selective cost.

Finally, we considered the metapopulation biology of *Caenorhabditis* nematodes, which involves colonization of ephemeral habitat patches, rapid population expansion, and then generation of new dispersal morphs (dauers) when the population exhausts its patch (Cutter, 2015; Felix & Braendle, 2010; Ferrari et al., 2017; Richaud et al., 2018). If a patch exists for a limited time, patches whose populations expand most rapidly will contribute more descendents to the overall gene pool when the patches expire. Chance genetic differences among patches can strongly influence the rate of population expansion (Figure 12 – figure supplement 2). Within each patch, allele frequencies experience the same forces as in a population of fixed size, but among patches we expect that those founded solely by homozygotes (experiencing no selective deaths from drivers) will have grown the largest after a few generations, increasing the representation of their alleles in the overall gene pool. As the rarer allele at a locus will be overrepresented in heterozygotes, and most homozygotes will carry only the common allele, this mode of population regulation should increase the frequency of the more common allele. This is a kind of “Haystack Model,” well studied in the context of sex ratios, where patchy environments favor a female bias because of the more rapid population expansion it allows (Bulmer & Taylor, 1980; Wilson & Colwell, 1981). These group-selection models are sensitive to parameters, including the number of individuals that found a patch, the number of offspring per parent, and the number of generations of growth within each patch. For each selfing rate, we modeled a situation in which each patch is colonized by four individuals, with sex ratio fixed to exclude its effects but genotypes drawn from the equilibrium frequencies given the allele frequency and selfing rate. The population then grows without any constraints on size until the patch reaches its expiration date, after three generations, at which time we calculate the global allele frequency, summing across the entire collection of patches. Under these conditions, we confirm a strong negative relationship between patch heterozygote frequency and patch growth rate. For a single driver, the effects of growth in a patchy environment on allele frequency are very modest, and the genic selection within each patch allows drivers to spread despite their costs. With antagonistic drivers, the reduction in heterozygote brood size is greater and the directional effects of single-driver genic selection are absent. In these conditions, group selection is sufficient to generate a global change in allele frequency (Figure 12C). There are more all-homozygote patches for the more common haplotype, and the result is positive frequency-dependent selection, decreasing the frequency of the less common haplotype. Overall, the joint effects of selfing and group selection on antagonistic drivers are therefore to eliminate the rarer allele. Selection achieves this result most efficiently at intermediate selfing rates. Although the drivers stick around at higher selfing rates, they impose very little genetic load, and at very high selfing rates they are entirely neutral.

## Discussion

We have brought genetic and genomic resources to the most recently discovered androdioecious *Caenorhabditis* species, *C. tropicalis,* facilitating studies of the varied effects of mating system transitions on genomes and metapopulation genetics, and their interactions with ecology, as well as comparative quantitative genetics. Aided by a chromosome-scale reference genome, our data confirm that *C. tropicalis* is indeed the most genetically homogeneous of the three selfers, on average, which implies a correspondingly high rate of selfing, low rate of effective recombination, and small effective population size. We also show that the average obscures extreme variance in the distribution of genomic diversity. This mirrors recent findings for *C. elegans* and *C. briggsae* (Lee et al., 2020). Within two highly divergent regions, we find some striking biology: three gene drive systems segregating in a single cross. Genetically similar drive elements have also been found in *C. elegans* (Ben-David et al., 2017; Seidel et al., 2008), but they must be especially common in *C. tropicalis.* They thus represent a potent and surprisingly widespread form of genetic incompatibility underlying outbreeding depression, and a potential cause of the species’ high effective selfing rate. Wth the addition of the third genome and genetic map for *Caenorhabditis* selfers, several other correlates of selfing rate, including the structuring of recombination, are apparent. Comparative analysis will be greatly furthered by more extensive sampling, in time, and in geographic and genomic space, for all selfing species and, as for *C. briggsae* and its outcrossing sister-species *C. nigoni* (Yin et al., 2018), study of *C. tropicalis’* closely related sister *C. wallacei* (Felix et al., 2014).

### Selfing and population genetics

Our early view of selfing *Caenorhabditis* species was of widespread, weedy lineages depauperate of genetic diversity relative to their outcrossing ancestors. This view, limited by mostly opportunistic sampling of isolates, and sequencing technology, was based on the apparent global expansion of *C. elegans* associated with human activity (Andersen et al., 2012). Better sampling has led to a more complete picture of strong population and genomic structure for all three species (Crombie et al., 2019; Thomas et al., 2015), though tropical areas remain particularly undersampled.

Most recently, a large survey of *C. elegans* genomes, including 15 assembled from long-reads, found regions of high-diversity spanning up to 20% of the reference genome (Lee et al., 2020). Similar heterogeneity was found in a smaller sample of *C. briggsae* genomes. This finding was presaged by efforts to build a complete genome for the divergent Hawaiian isolated CB4856 using 2^nd^ generation sequencing (Thompson et al., 2015), but has been greatly enabled by more contiguous assemblies that circumvent the reference mapping bias plaguing study of all genetically diverse species. A promising hypothesis for the presence of hyperdivergent regions in genomes is the action of balancing selection across the species’ range, leading to preservation of some of the abundant genetic diversity of outcrossing ancestral species. This hypothesis is supported in *C. elegans* by the enrichment of genes encoding environmentally responsive sensory factors, which are themselves enriched for differential expression and quantitative trait loci for response to microbes isolated from natural habitats. Alternative hypotheses include introgression, a common source of islands of divergence in other animal taxa (Hedrick, 2013), but the presence of multiple distinct divergent haplotypes in *C. elegans* (Lee et al., 2020) argues strongly against it. Balancing selection can act at many levels, from local adaptation via directional selection, at the global scale, to various frequency-dependent phenomena at the local scale, and overdominance at the molecular scale. While some environmental associations clearly play a role in the biogeography of all three androdioecious species, they seem not to be a definitive factor in structuring divergent haplotypes. The high migration rate that comes with the microscopic nematode form, coupled with the reproductive assurance afforded by selfing, may effectively counter adaptation to global environmental variation. Our results suggest that population structuring of divergent regions varies across the three selfers, but this will need to be revisited with *C. elegans-*scale sampling of *C. briggsae* and *C. tropicalis.*

Taxa with mixed selfing and outcrossing, very large population sizes, and broad, diverse ranges, may occupy a population genetic space particularly well-suited to the detection and localisation of balancing and local selection. The ability to detect targets of strong balancing selection scales with total population size (assuming symmetric migration) and recombination rate, and the homogenising effect of partial selfing is to increase the signal of balanced peaks against a background continually swept of diversity by indirect selection (Charlesworth et al., 1997; Nordborg et al., 1996). The particularly high selfing rate of *C. tropicalis,* together with changes in recombination between chromosome arms and centers, may be especially favorable, and its preferred habitat may also allow for more rapid evolution than the temperate-dwelling *C. elegans.* Detection of the targets of local adaptation depends on population size, migration rate and selection intensity (Charlesworth et al., 1997), which are largely unknown. Although gene drive elements have the potential to generate balancing selection, we find that most divergent regions between NIC58 and JU1373 do not carry gene drive elements, or, at least, not ones that drive independent of environment and genetic background. These, and other highly divergent haplotypes in *C. tropicalis,* may harbor loci under positive selection in different conditions, a conjecture made more plausible by the analysis of divergent gene content in *C. elegans.*

### Population dynamics of drivers

Genetic drive elements like those we discovered in *C. tropicalis* will reliably spread in randomly mating populations (Wade & Beeman, 1994). Highly penetrant killing of homozygous larvae is an exceptionally potent selective mechanism to drive allele frequency change, and in our short RIL construction pedigree we saw two haplotypes sweep nearly to fixation. In nature, things are likely quite different, as both the mating system and natural history of *C. tropicalis* conspire to render drive loci selectively inert. *C. tropicalis* shows strong geographic structure, presumably exacerbated by habitat fragmentation, such that encounters between driver and sensitive haplotypes may be rare. Upon an encounter between a hermaphrodite and a male, outcrossing rates, measured under benign laboratory conditions, are relatively low on average. When divergent isolates do cross, drivers may find themselves inactive due to dependence on genetic background, including mitochondrial genotype, and, potentially, dependence on environmental factors. Most importantly, *C. tropicalis* reproduces primarily by self fertilization, and drive elements are unable to gain traction when heterozygotes are infrequent. At the same time, the patchy, ephemeral microhabitat of *C. tropicalis* – rotting fruits and flowers on the forest floor – provides a perfect substrate for group selection. Small numbers of dispersing larvae colonize each patch and undergo exponential population growth for a small number of generations. Although drivers will increase in frequency in patches with heterozygotes, population growth in patches without heterozygotes can be so much greater as to overwhelm the countervailing effects of drivers on allele frequency.

The patterns we observe on chromosome V implicate tightly linked drivers, one on each of the alternative haplotypes, a phenomenon also discovered by Ben-David et al. (2020). Surprisingly, we found by simulation that antagonistic maternal toxin-zygotic antidote drivers do not generate balancing selection, at least under the symmetric-effects scenario we modeled. They nevertheless impose a strong segregation load under outcrossing, which should select for suppressors. The antagonistic drivers themselves evolve by drift at high selfing rates, and at intermediate selfing rates frequency-dependent selection eliminates the rarer haplotype. At the same time, the chromosome V drivers occur on ancient haplotypes, evidenced by extreme divergence between NIC58 and JU1373 (Figure 7; Figure 10). These haplotypes may encode unique toxin-antidote pairs that arose independently, or they may encode toxin-antidote pairs that co-evolved from a common ancestor but are no longer cross-compatible. Competition among driver haplotypes is known to occur for Segregation Distorter in *D. melanogaster,* but in this case, driver haplotypes compete for slots in a balanced equilibrium with non-drivers (C. L. Brand et al., 2015; Presgraves et al., 2009). Our simulations suggest that antagonistic drive is unlikely to play a role in the ancient balancing selection at the chromosome V locus, and we note that the two haplotypes are sufficiently different in gene content that effects on phenotypes other than drive are likely.

### Molecular mechanisms of drive

The drivers we have discovered in *C. tropicalis* are analogous to the *sup−35/pha−1* maternal-effect driver (Ben-David et al., 2017) and the *zeel-1/peel-1* paternal-effect driver in *C. elegans* (Seidel et al., 2008, 2011), and four maternal-effect *Medea* drivers in *Tribolium* (Beeman et al., 1992; Beeman & Friesen, 1999). Additionally, several maternal-effect drivers have been independently identified in *C. tropicalis* by Ben-David et al. (2020). Similar inheritance patterns have also been reported for two loci in mice (Peters & Barker, 1993; Weichenhan et al., 1996, 1998; Winking et al., 1991). The causal genes underlying the JU1373 and NIC58 drivers remain to be identified, but likely include one or more of the multiple genes unique to driver haplotypes, as seen for *C. elegans* where toxin and antidote functions are encoded by genes present on the driver haplotype and absent (or pseudogenized) on the non-driver haplotype (Ben-David et al., 2017; Seidel et al., 2008). Similarly, the single *Medea* driver whose genetic basis is known maps to a transposable element insertion absent from non-driver haplotypes (Lorenzen et al., 2008). A common pattern emerging from these systems is that maternal-effect drivers (and the single example of a paternal-effect driver) are encoded by dispensable genes with dedicated functions, rather than genes acquiring toxin or antidote activity while retaining an ancestral non-drive function.

Why are these drivers so prevalent in *C. tropicalis* (and to some extent in *C. elegans*) but mostly absent elsewhere? One option is ascertainment bias – maybe similar elements are taxonomically more widespread, but we simply haven’t looked for them. A second option is that mechanisms of translational control in the *Caenorhabditis* germline may make it easy for maternal-effect toxins to arise. Early embryogenesis in *Caenorhabditis* is largely controlled by maternal regulators, and a common expression pattern for these regulators is ubiquitous expression of mRNA in the oocyte but no translation until embryogenesis (Evans & Hunter, 2005; Robertson & Lin, 2015). A gene whose protein is generally cytotoxic could become a maternal-effect toxin by acquiring the (common) regulatory elements specifying this expression module; tight linkage to a zygotically expressed antidote would create a driver. A third, non-mutually exclusive option is that a selfing reproductive mode may promote cycles of compensatory substitutions that help create or strengthen toxin-antidote pairs. Partially penetrant toxins might arise in a background lacking an antidote but become locally fixed, despite their deleteriousness, due to the tiny effective population size of a local selfing population. If outcrossing is rare, toxin-free haplotypes will not be re-introduced or decoupled by recombination and instead, the population might restore its fitness via compensatory evolution of an antidote; if the antidote is linked to the toxin, a driver is born. This third explanation surmises that toxin-antidote pairs arise in populations that have already transitioned to selfing or, for outcrossing species, have small effective population sizes due to high levels of inbreeding. Whether the toxin-antidote elements in *C. tropicalis* and *C. elegans* arose before the transition to selfing is unclear, although the level of divergence between opposite haplotypes at drive loci is suggestive of sampling from outcrossing ancestors. A closer examination of outcrossing species in the Elegans group is needed to determine whether toxin-antidote elements are specific to, or quantitatively different in, selfers. It will also be interesting to see the strong outbreeding depression in other taxa with mixed-mating dissected genetically, such as the “cryptic biological species” complexes in arctic *Draba* (Grundt et al., 2006).

### Strategies to combat drivers

Gene drive systems create a selective environment favoring the evolution of suppressors. Suppressors of gene drive have been well documented in many species, especially when obligate outcrossing continually exposes individuals to the costs of drive (Courret et al., 2019; Lindholm et al., 2016; Lyttle, 1991; Price et al., 2019). Suppressors of drive in selfing species are more rare, which has been interpreted as evidence that many drivers in selfing species did not evolve as drivers per se but instead evolved through non-drive mechanisms (Sweigart et al., 2019), such as balancing selection maintaining alternate homozygous genotypes (Seidel et al., 2008). Our data show that unlinked modifiers affect driver activity in *C. tropicalis,* though whether these modifiers evolved as suppressors is equivocal. In the case of maternal-effect drivers, mitochondrial suppressors are special: selection for mitochondrial suppressors may be especially strong because mitochondria cannot segregate away from drivers via inheritance in sperm. This selective environment may explain why two of the drivers we discovered in *C. tropicalis* (the chromosome III JU1373 driver and the chromosome V NIC58 driver) were differentially active according to mitochondrial genotype – the mitochondrial genotypes non-permissive for drive may have evolved as suppressors. Alternatively, drivers may have arisen in mitochondrial backgrounds that were permissive for drive activity, with little selection for or against alternate mitochondrial genotypes. Ultimately, the data provide little conclusive evidence that the drive loci experience selection in nature that is due to their drive activity.

Our data suggest that crosses between geographically distant *C. tropicalis* isolates will typically reveal multiple drive loci (Figure 11). Segregation of multiple drivers magnifies the cost of outcrossing and reduces the possibility of suppression by a common molecular mechanism, in a manner analogous to the role of multidrug therapy in preventing the evolution of drug resistance. The difficulty that organisms face in evolving suppressors to multiple drive elements at once has emerged as an important consideration for gene drive strategies for controlling disease vectors (Burt, 2003; Champer et al., 2018). In such cases, organisms can adapt by altering their population biology, increasing their rates of inbreeding and selfing (Bull, 2017; Bull et al., 2019; Drury et al., 2017), and thus reducing the heterozygosity required for all driver activity.

The costs of selfing as a defense against gene drive are inbreeding depression; reduced ability to adapt to new conditions; and reduced genetic variation and hence niche breadth. Androdioecious *Caenorhabditis* appear to have mechanisms for dealing with each of these costs. Selfing *Caenorhabditis* are typically found in nature as totally inbred lines, consistent with having purged recessive deleterious variants in their history (Anderson et al., 2010). While outcrossing plays an important adaptive role in selfing *Caenorhabditis* (Chelo et al., 2019; Morran et al., 2009; Teotonio et al., 2006; Teotonio et al., 2012), populations can transiently increase male frequency to achieve adaptation before returning to a primarily selfing mode of reproduction (Anderson et al., 2010; Shi et al., 2017). Finally, the preservation of genetic diversity at large numbers of ancient haplotypes by balancing selection allows these species to occupy a wide range of habitats despite very low levels of baseline genetic variation (Lee et al., 2020).

We have shown that *C. tropicalis* harbors abundant heritable variation in outcrossing rate, with nondisjunction, male mating ability, and hermaphrodite mating ability all providing avenues for genetic fine-tuning of the outcrossing rate. Other data also show that *C. tropicalis* has mechanisms that promote selfing over outcrossing. Ting et al. (2014) found that *C. tropicalis* hermaphrodites are uniquely resistant to the deleterious effects of interspecific matings, and they interpret their findings as evidence for reduced activity of sperm guidance cues in *C. tropicalis* hermaphrodites. Shi et al. (2017) showed that male longevity is reduced in *C. tropicalis* when male pheromone is present, creating a negative feedback that tamps down male frequencies, but not in obligately outcrossing *Caenorhabditis.* These findings are consistent with selection favoring high selfing rates in *C. tropicalis.*

Selfing is often considered a factor that favors the evolution of incompatibilities and outbreeding depression, just as the variably independent evolution of species or subspecies often leads to incompatibilities revealed by hybridization (Fishman & Sweigart, 2018; Maheshwari & Barbash, 2011; Presgraves, 2010). Selfing reduces the effective recombination rate, allowing unlinked loci to evolve together. When outcrossing reshuffles these co-evolved loci, it creates new combinations of alleles untested by selection. Incompatibilities between these alleles manifest as outbreeding depression (equivalently, rearrangements can fix within selfing lineages, rendering outbred progeny deficient). Our findings suggest that we should also consider causation running in the opposite direction. Incompatibilities in *C. tropicalis* appear to mostly represent interactions between tightly linked loci acting in different individuals (mothers and offspring), rather than interactions between unlinked loci; thus, the outbreeding depression caused by these incompatibilities is mostly not mediated by recombination. *C. tropicalis* can escape these incompatibilities and restore fitness by inbreeding. Thus, in contrast to the usual pattern of selfing leading to incompatibility, in this species incompatibility may also lead to increased selfing.

## Methods

### Strain maintenance

Strains were maintained using standard protocols for *C. elegans* (Brenner, 1974; Stiernagle, 2006), with the addition of 1.25% agarose to NGM-agar (NGMA) plates to discourage burrowing, and a 25C incubation temperature. This temperature is characteristic of substrate temperatures where we have collected *C. tropicalis,* and is the standard rearing temperature in previous work on this species (Gimond et al., 2013).

### Genome sequencing

Long-read data for NIC58 and JU1373 were around 250x expected coverage, given an estimated genome size of roughly 80 Mb (Fierst et al., 2015), from a PacBio Sequel at the Duke University Center for Genomic and Computational Biology. DNA was extracted from twelve 10 cm NGMA plates of nematodes spotted with OP50 using the Qiagen MagAttract HMW DNA kit as per (Lee et al., 2020).

JU1373 and NIC58 short-read data were around 25x and 40x expected coverage 100 bp paired-end reads (TruSeq libraries, HiSeq 2000, NYU Center for Genomics and Systems Biology Genomics Core), and another 40x coverage for NIC58 (150 bp paired-end reads, TruSeq library, NovaSeq6000, Novogene).

We sequenced 129 RILs from a cross between JU1373 and NIC58 to a median depth of 2.1x (NextEra libraries using the protocol of Baym et al. (2015), NextSeq 500, paired end 75 and 150 bp reads, NYU Center for Genomics and Systems Biology Genomics Core). DNA was isolated by proteinase-K digestion followed by phenol/chloroform/isoamyl alcohol purification.

An additional 22 wild isolates were sequenced to a median depth of 29x (NextEra libraries, NextSeq 500, paired end 75 bp reads; or Bioo libraries, HiSeq 2000, paired end 100 bp reads; NYU Center for Genomics and Systems Biology Genomics Core). DNA was extracted by salting out (Sunnucks & Hales, 1996). Isolates and associated metadata are in Figure 5 – source data 1.

All sequencing reads used in this project are available from the NCBI Sequence Read Archive under accession PRJNA662844.

### Genome assembly

Our reference genome is NIC58. We generated initial assemblies for evaluation with genetic linkage data, including a Canu hybrid assembly (Koren et al., 2017) and long-read only assemblies from flye (Kolmogorov et al., 2019), ra (Vaser & Sikic, 2019) and wtdbg2 (Ruan & Li, 2019). All were initially run with default parameters. Flye produced a highly contiguous assembly with this data set, and initial genetic evaluation showed few errors, so we varied parameters (minimum overlap length 4–10 kb, initial assembly depth 40–180x) and selected the two most contiguous assemblies for closer evaluation (the genetically concordant assembly used -m 10 kb --asm-coverage 120x). A draft assembly for JU1373 was made with flye using default parameters (44 contigs and scaffolds, NG50 4.2 Mb, 81 Mb span).

Both assemblies were polished with short-reads using Pilon (-fix bases mode) before further use (Walker et al., 2014). Mitochondrial genomes were initially assembled from long reads mapping to contigs identified as partially homologous to *C. elegans* sequence. *De novo* assemblies using Unicycler (Wick et al., 2017) to produce a polished circular sequence showed homology to all 12 *C. elegans* proteins for both NIC58 and JU1373, but total length and sequenced identity were sensitive to input read length (using all data, or only reads of length 10–15 kb, which spans the range of full-length *Caenorhabditis* mitochondrial genomes in GenBank) and mapping quality. We instead used fragmented, high-coverage contigs from lllumina *de novo* assemblies (Platanus 1.2.4; (Kajitani et al., 2014)) with homology to the long-read assemblies as bait to extract short reads for reassembly, which produced single sequences of length 13565 and 13091 bp for NIC58 and JU1373, respectively. After circular polishing with long-reads (Unicycler), sequences were 14394 and 14027 bp. We rotated these with five copies of the *C. elegans* mitochondrial genomes to optimise linear homology using MARS (Ayad & Pissis, 2017).

### Genetic map construction

RILs were derived by crossing a NIC58 male and JU1373 hermaphrodite, and inbreeding the F_2_ offspring of a single F_1_ hermaphrodite for 10 generations by selfing. We used the RIL data to evaluate assemblies based on 1) interchromosomal consistency and 2) concordance between genetic and physical order. A SnakeMake pipeline (Koster & Rahmann, 2012) implementing this procedure is on [github](http://github.com/lukemn/tropicalis). and we omit full details of software versions and parameters here.

Using short-read mapping to the NIC58 assemblies, we called variants distinguishing the parental lines, filtered them to homozygous diallelic SNVs (depth within 1/3 of the median, > 10bp from an insertion/deletion, quality > 50, then pruned to remove any SNVs in 20 bp windows with more than one SNV), and genotyped the RILs at these sites (H. Li, 2011; H. Li et al., 2009; H. Li & Durbin, 2009; Vasimuddin, Md et al., 2019).

Parental ancestry was inferred by HMM (Andolfatto et al., 2011), sampling one variant per read, with transition probabilities defined by homozygous priors, recombination rate (r= per base pair rate given an expected 6 recombination events per RIL genome), physical distance between markers in the reference genome *(d*) and a scaling factor (*rfac=* 10^ࢤ11^), parameterised as −10^ࢤrWac^, and emission probabilities set by parental genotyping error rate (10 ^ࢤ4^) and base quality scores. Markers for map construction were constructed by filtering on posterior probability > 0.5, binning up to 50 SNVs, and merging the sparse RIL marker inferences, interpolating missing positions across consistent uniparental flanking bins. Bins with both parental genotypes were considered as missing data.

Marker filtering and map construction was carried out in R/qtl (Broman et al., 2003). After dropping identically informative genotypes, two lines that were outliers for heterozygosity, and one of each of eight pairs of lines with >99% similarity, linkage groups (LGs) were formed (maximizing the number of markers in six linkage groups), and markers were ordered within LGs by likelihood from 100 iterations of greedy marker ordering.

Where genetic and physical ordering conflicted, the physical order was tested by likelihood and accepted if the change in LOD was > −1. Taking the genetic data as ground truth, we compared assemblies on the number of sequences spanning more than one LG, and on the number and sum of negative LOD scores for any remaining discordance in within-LG genetic/physical marker order. On these metrics, we selected a flye assembly, spanning 81.83 Mb in 36 contigs and scaffolds > 20 kb with an N50 of 4.795 Mb. The genetic map based on the assembly incorporated 33 of these sequences and spanned 81.3 Mb. We then did two rounds of manual stitching, considering only junctions with estimated genetic gaps of 0 cM. First, we accepted 10 joins where sequences from another assembly spanned a junction (>5 kb of MQ=60 alignment on either flank; minimap2 (H. Li, 2018)). Second, we accepted 8 joins where at least one read consistent with the genetic orientation spanned a junction (alignment >2 kb of MQ>20 on each 50 kb flank; minimap2). We took the consensus sequence (bcftools), or in two cases the read sequence, and converted the now fully-oriented assembly of 15 sequences into pseudochromosomes, with 50 bp N gaps at the remaining junctions. Chromosomes were named and oriented based on *C. elegans* homology, by summing aligned lengths per chromosome and strand (minimap2 -x asm20 mapping quality > 30). Chromosome preference was unequivocal (>60-fold bias toward a single homolog). Strand preference was relatively strong for chromosomes II and X (>3.2-fold bias), but less so for the others (1.8-fold bias for IV, 1.4 for I, 1.3 for III, and 1.1 for V), from 0.400–1.9 Mb of aligned sequence per homologous chromosome. The inferred orientations were consistent with strand bias from 1:1 orthologs in chromosome centers in all cases except chromosome I. Finally, we did one further round of short-read polishing (pilon -fix bases mode, making 5044 changes), and re-estimated the genetic map.

### Annotation

#### Mixed-stage RNA preparation

We collected three samples each for NIC58 and JU1373: well fed mixed-staged (L1-adults), well fed male-enriched, and starved (including dauers) plates. Strains were passaged by chunking every 2 days to maintain a well-fed mixed-stage population. Some plates were allowed to starve, and the presence of dauer larvae along with other developmentally arrested larvae was confirmed by visual inspection. Crosses were set up on single-drop OP50-seeded plates with 15–20 males and a few hermaphrodites to establish a male-enriched population. Following successful mating, worms were chunked to 10 cm OP50-seeded plates for sample collection.

Each sample was collected from one 10 cm plate, flash frozen in 100 μl S-Basal in liquid nitrogen, and stored at −80C until RNA extraction. Total RNA was extracted using Trizol reagent (Invitrogen) following the manufacturer’s protocol, except that 100 μl acid-washed sand (Sigma) was added during the initial homogenization step. RNA was eluted in nuclease-free water, purity was assessed by Nanodrop (ThermoFisher), concentration was determined by Qubit (ThermoFisher), and integrity was assessed by molecular weight on a Bioanalyzer (Agilent). Following quality control, 1.5 μg of total RNA from each sample was pooled, further purified using the RNA MinElute Cleanup kit (Qiagen), and again subject to the above quality control analyses.

#### Library preparation and sequencing

Libraries for JU1373 and NIC58 were prepared simultaneously from mRNA isolated from 1 μg of pooled total RNA using the NEBNext Poly(A) mRNA Magnetic Isolation Module (New England Biolabs). RNA fragmentation, first and second strand cDNA synthesis, and end-repair processing were performed with the NEBNext Ultra II RNA Library Prep with Sample Purification Beads (New England Biolabs).

Adapters and unique dual indexes in the NEBNext Multiplex Oligos for lllumina (New England Biolabs) were ligated, and the concentration of each library was determined using Qubit dsDNA BR Assay Kit (Invitrogen). Libraries were pooled and qualified by Bioanalyzer 2100 (Agilent, at Novogene, CA, USA), and 150 bp paired-end reads were sequenced on a single lllumina NovaSeq 6000 lane.

#### Annotation

Prior to predicting genes, we identified repetitive sequences in the NIC58 and JU1373 genomes de novo using RepeatModeler (Smit et al., n.d.) and classified these using the RepeatClassifier tool from RepeatModeler and the Dfam database (Hubley et al., 2016). We removed unclassified repeats and soft-masked the genome assemblies with RepeatMasker using the classified repeat library. We aligned short RNAseq reads to the soft-masked genomes with STAR in two-pass mode (Dobin et al., 2013), and used the BRAKER pipeline to annotate genes (Hoff et al., 2019). We extracted protein sequences from the BRAKER annotation using the getAnnoFasta.pl script from AUGUSTUS (Stanke et al., 2006), and assessed biological completeness using BUSCO (Seppey et al., 2019). We annotated mitochondrial genomes by homology using the MITOS2 server (Bernt et al., 2013) and Prokka (Seemann, 2014).

### Macrosynteny and orthology

We downloaded sequence and annotation data for other *Caenorhabditis* species with chromosomal genomes from WormBase (Harris et al., 2020) and the CGP: *C. elegans* N2 and VC2010 (WS277; (Yoshimura et al., 2019)), *C. inopinata* NK74SC (WS277; (Kanzaki et al., 2018)) and *C. briggsae* AF16 (WS235; (Ross et al., 2011), and for *C. remanei* PX506 (GCA_010183535.1; (Teterina et al., 2020)) from NCBI.

Orthologs were identified with OrthoFinder v. 2.4.0 (Emms & Kelly, 2019), using canonical proteins where annotated. Where not annotated (*C. remanei*) we removed proteins from alternative transcripts with cd-hit (-c 0.85 -G 0 -aL 0.2 -aS 0.2; (W. Li et al., 2001)). Examining predicted protein-coding genes under driver intervals, we define novelty as an orphan orthologous group unique to a single *C. tropicalis* strain among the studied species. We used blastp against the NCBI nr and [CGP](http://caenorhabditis.org/) databases (word size 3, expect value < 1e^ࢤ2^) to confirm novelty more broadly, and InterPro to search for known protein domains (Hunter et al., 2009).

Reference-free genome alignments for all species were from cactus v. 1.0.0 (Paten et al., 2011), reference-based alignments for *C. tropicalis* used Mummer v. 4.0.0 (Malyais et al., 2018), and minimap2 v. 2.17-r954 (-x asm20) and paftools to call variants (-I 1000 -L 1000; (H. Li, 2018)).

### Population genetics

#### Variant calling

A SnakeMake pipeline implementing variant calling and filtering is available from [github](https://github.com/lukemn/tropicalis) (Koster & Rahmann, 2012). In brief, we mapped reads to the NIC58 reference genome with bwa mem2 (Vasimuddin, Md et al., 2019), aligned and normalized indels with bcftools (H. Li, 2011), called variants jointly with GATK (DePristo et al., 2011; McKenna et al., 2010), and hard filtered diallelic SNVs (median absolute deviation in total depth < 99th percentile, QD > 4, MQ > 30, BaseQRankSum > −3, ReadPosRankSum > −4, SOR < 5). We also applied per-sample depth filtering (local depth in 1 kb windows < 2x against a LOESS polynomial fit for each chromosome, span=0.33), keeping SNVs in windows where at least 22/24 samples passed. A total of 880,599 diallelic SNVs were called, 794,676 passed filtering (genotype set 1), and we used the fully homozygous subset of these with no missing data, comprising 397,515 SNVs (genotype set 2).

#### Population structure

Principal component analysis was carried out on the additive genetic relationship matrix (base R ‘prcomp’) constructed from homozygous diallelic SNVs with no missing data (genotype set 2).

#### Recombination rate domain definition and genomic analysis

Arm/center junctions were estimated from a three-segment linear regression of cumulative physical distance on genetic distance, excluding tip domains, using the R package strucchange v. 1.5-1 (Zeileis et al., 2002). Tips were assigned based on terminal genetic positions. The same approach was applied to genetic maps for *C. elegans* (Rockman & Kruglyak, 2009) and *C. briggsae* (Ross et al., 2011) for the sake of consistency, resulting in minor differences with previous segmentations. Except where noted otherwise, subsequent comparisons across the three species were on maps standardized by linear interpolation to 1000 markers per chromosome.

Variation in recombination density across species was quantified as the Gini coefficient, or 1 minus twice the area under the curve described by recombination per Mb, sorted in ascending order and cumulated, and corresponding cumulative physical distance (Kaur & Rockman, 2014). Summary statistics across chromosomes were taken in 100 kb non-overlapping bins (Quinlan & Hall, 2010): GC-content, gene and exon density and spacing, mean zlib compression (level nine) ratio for non-coding 100mers (an approximation of Kolmogorov complexity), Watterson’s estimator (θ_W_) and π based on diallelic SNVs from our sequencing of 24 wild isolates for *C. tropicalis,* the *C. elegans* Natural Diversity Resource (330 isotypes in CeNDR release 20180527; (Cook et al., 2017)), and data from 35 sequenced *C. briggsae* lines (Thomas et al., 2015), kindly provided by A. Cutter.

To test the relationship between recombination rate and nucleotide diversity we used nested quasibinomial linear models and likelihood ratio tests. Data were binned genetically to the mean number of unique genetic positions in the observed genetic maps (547) to avoid pseudoreplication.

#### Divergent regions

We thresholded divergent regions using kernel density smoothing of the empirical distribution of θ_W_ across genomic windows (10 kb), taking the first positive value of the first derivative, after the minimum, as the threshold value (Duong, 2020). Regions were enumerated based simply on contiguous runs of the sign of the second derivative of θ_W_, that is, all local peaks in nucleotide diversity are treated as independent. This makes the unrealistic assumptions of uniform ancestral diversity and effective recombination, and is sensitive to sample size and binning. Deeper and broader population genetic data will be required to obtain more confident estimates of the number, size and local structure of divergent regions, ideally with multiple high-quality genome assemblies to minimize confounding by reference mapping bias.

### Statistical analysis, data wrangling, plotting

We used R (R Core Team, 2018) with packages data.table (Dowle & Srinivasan, 2019), dglm (Dunn & Smyth, 2016), dplyr (Wickham et al., 2020), ggmap (Kahle & Wickham, 2013), ggplot2 (Wickham, 2016), ggh4x (T. van den Brand, 2020), ggrepel (Slowikowski, 2020), Ime4 (Bates et al., 2015), and tidyr (Wickham & Henry, 2020).

### Mating trials among isolates and RILs

Mating trials were initiated with one L4 hermaphrodite and one L4 male worm on a 6 cm NGM agarose plate seeded with 50 μΙ_ of OP-50 *E. coli.* Plates were scored 72 hours later, with success defined as the presence of multiple males in the F_1_ generation.

We scored hermaphrodite cross probability in RILs by crossing NIC58 males to L4 RIL hermaphrodites. The number of RIL trials ranged from 16 to 75, with a median of 30, and trials took place over 116 days. A total of 338 JU1373 and 412 NIC58 control crosses were done on 107 of these days.

To estimate equilibrium male frequency, we scored sex ratio after 10 generations of passaging at large population size. Three L4 hermaphrodites and 5 L4 males were placed on a 6 cm agarose plate. Three days later, 3 mL of M9 buffer was pipetted onto the plate and 50 μL of worms was transferred to a 10 cm plate. 50 μL of worms was subsequently transferred to a 10 cm agarose plate every ~72 hours for 10 generations, at which point a sample of 267 ± 27 worms were sexed per strain. We performed three replicates of this passaging experiment.

Phenotypes for RIL QTL mapping were best linear unbiased predictions (BLUPs) from a binomial linear mixed-effects model (R package Ime4; (Bates et al., 2015)). Mapping in R/qtl (Broman et al., 2003) used a ‘normal’ model, and 1000 permutations of the phenotype values to establish genome-wide significance. The variance explained by the single significant QTL was estimated from variance components by refitting the linear model to the raw data with a random effect of genotype within RIL.

### Genetic analysis of drive activity

We used a standardized assay of the proportion of F_2_ embryos that develop to adulthood according to wild-type schedule. P_0_ males and hermaphrodites were paired as L4s, and the following day each hermaphrodite that bore a copulatory plug was transferred to a fresh plate to lay embryos. Two days later, when these embryos had developed to L4 stage, we isolated F_1_ hermaphrodites overnight. The following day, hermaphrodites were singled to new plates and left to lay embryos for 8 hours. The hermaphrodites were then removed and the embryos on each plate counted. Three days later, when wild-type animals have reliably reached adulthood, we counted the adults on each plate by picking. In some cases, slow-developing animals that had reached L3 or L4 were counted separately. The majority of driver-affected animals arrest as L1s and are difficult to see, so we typically estimated the number of arrested larvae as the number of embryos initially observed minus the number of adults counted three days later. In a small number of broods (~3%), the count of progeny at adulthood exceeded the count of embryos laid (by at most two extra adults, from broods containing ~35–55 total embryos laid). We made the assumption that this discrepancy reflected undercounting of embryos rather than overcounting of adults, given that embryos are hard to see. Thus, for such broods, we adjusted the embryo count upward to match the count of adults. All conclusions are robust to this adjustment.

Because of the low mating efficiency of many *C. tropicalis* genotypes, matings did not always produce cross progeny. To distinguish self and cross progeny, we employed several approaches. In some experiments, we depleted hermaphrodites of sperm by transferring them to fresh plates on each of the first four days of adulthood, until they ceased reproduction. These sperm-depleted hermaphrodites could then be crossed to males, and resulting progeny inferred reliably to be cross offspring. This method is not suitable for all experiments because the sperm-depleted hermaphrodites have small broods and generally show age-associated decrepitude. As an alternative, we developed visible marker strains that allow us to distinguish self and cross progeny. We isolated a spontaneous Dumpy mutant in the JU1373 background and established strain QG2413, *Ctr-dpy (qg2).* Control experiments confirmed that this semidominant mutation allows for clean discrimination between Dpy and semi- or non-Dpy animals, and that the mutation is unlinked to the drive loci on chromosomes III and V. Next, we generated a NIC58 derivative carrying a fluorescent reporter. Strain QG3501 (*qgls5*) carries pCFJ104 *[Pmyo-3::mCherry::unc-54utr]* (Fr_0_kjaer-Jensen et al., 2008). The transgene was introduced by microinjection into NIC58, integrated by UV, and backcrossed to NIC58 seven times. These animals express bright red fluorescence in muscle, visible under the dissecting scope from mid embryogenesis. Control experiments show that this transgene is unlinked to the drive loci.

Inheritance at drive loci was tracked using PCR to amplify insertion/deletion markers near the segregation distortion peaks:

LG3.1336F TTAGAGCCGCTTGAAGTTGG
LG3.1336R TCCGATGGACTAGGTTTCGT
LG5.2017F TAACGCAATGGCCTCCTATC
LG5.2017R GTTTGCTGGGTGGCCTAGTA

### Simulations

We used simulations to investigate the effects of selfing on the spread of a maternal-toxin/zygotic antidote haplotype in populations with the distinctive androdioecious mating system of *C. tropicalis.* Each simulated individual has a drive locus genotype and a sex. The drive locus has haplotypes D and d. The former carries a maternal-effect toxin and zygotic-effect antidote, and the latter carries neither. We initiate a population with *N* individuals and genotype frequencies and sex ratio that are at equilibrium given the population’s selfing rate S and frequency *p* of the drive haplotype D. The equilibrium inbreeding coefficient 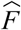 is S/(2-S) and the male frequency is 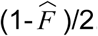.

For simulations in Figure 12A and B, with fixed population sizes, we generated starting populations using the equilibrium frequencies below.

| Genotype | Frequency |
| --- | --- |
| DD herm | $((1-F)p^2 + pF)(1+F)/2$ |
| Dd herm | $(1-F)2p(1-p)(1+F)/2$ |
| dd herm | $((1-F)(1-p)^2 + (1-p)F)(1+F)/2$ |
| DD male | $((1-F)p^2 + pF)(1-F)/2$ |
| Dd male | $(1-F)2p(1-p)(1-F)/2$ |
| dd male | $((1-F)(1-p)^2 + (1-p)F)(1-F)/2$ |

For simulations in Figure 12C, assessing the effects of patch dynamics on haplotype frequencies of antagonistic drivers, we draw genotype frequencies from a multinomial according to the equilibrium frequencies, but we assign sexes deterministically to ensure every patch receives hermaphrodite founders.

We modeled the *C. tropicalis* androdioecious mating system, with self-fertile hermaphrodites that are incapable of mating with one another, and males that can cross-fertilize hermaphrodite oocytes. If no male is present in a population, each hermaphrodite produces a brood of size *B* by selfing. If there are males, each hermaphrodite produces *SB* hermaphrodite offspring by selfing and (1 *-S)B* offspring by mating (male or hermaphrodite with equal probability), with a single father drawn randomly from the population of males. Self progeny are mostly hermaphrodites, except each has probability *Him* of being male (Him is the worm community name for the rate of male production by X-chromosome nondisjunction in selfing hermaphrodites).

For simplicity, we assume that all selection is on embryo viability and that there is no cost to the drive allele. For single-driver analyses (Fig 12A and B), individuals that are dd but have Dd mothers are viable with probability *V*(i.e., if V>0 some embryos can survive the maternal-effect toxin). Everybody else has viability 1.

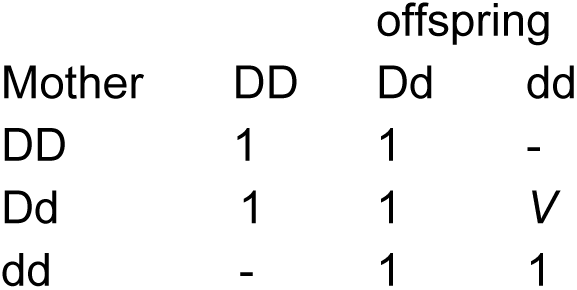

In simulations with antagonistic drivers, we assume the penetrance of the alternate drivers is identical, yielding the following viabilities:

| Mother | offspring |  |  |
| --- | --- | --- | --- |
|  | DD | Dd | dd |
| DD | 1 | 1 | - |
| Dd | $V$ | 1 | $V$ |
| dd | - | 1 | 1 |

In simulations with fixed population size, each discrete generation is sampled from the pool of viable embryos. We then track p, the frequency of the D allele, given parameters *N, S, B, V,* and *Him.* In simulations with exponential growth, *p* and *N* are both variables. For all simulations described in the text, we used *B* = 50, *V=* 0.05, and *Him =* 0.005. We then investigate the effects of population size (*N*) and selfing rate (S) on drive allele frequency *p.* Simulations started with allele frequency of 0.05 for single-driver scenarios, and with frequency 0.2 for antagonistic-driver scenarios where frequencies were tracked for only a few generations. Simulation code is available on [github](https://github.com/lukemn/tropicalis).

## Source Data Files

Figure 1 – source data 1: outcrossProbability.tsv.zip; wild isolate outcross probability trials.

Figure 1 – source data 2: malePassaging.tsv.zip; wild isolate spontaneous male frequency.

Figure 2 – source data 1: selferMaps.tsv.zip; genetic maps for *C. elegans, C. briggsae,* and *C. tropicalis.*

Figure 2 – source data 2: selferRecombinationRateDomains.tsv.zip; recombinate rate domain breakpoints for *C. elegans, C. briggsae,* and *C. tropicalis.*

Figure 2 – source data 3: genomeStats.zip; archive containing genome sequence summary statistics; genomeStats_codingBases.tsv, genomeStats_compressability.tsv, genomeStats_geneDensity.tsv, genomeStats_nuc.tsv, genomeStats_selferTreeConcordance.tsv.

Figure 3 – source data 1: caeno_o2oOrthologs.tsv.zip; genomic positions of 1:1 orthologs across *Caenorhabditis* species.

Figure 3 – source data 2: caeno_cactusAlignment.tsv.zip; *Caenorhabditis* Cactus alignment blocks, with NIC58 reference genome.

Figure 3 – source data 3: JU1373-NIC58_mummerAlignment.tsv.zip; JU1373 and NIC58 synteny.

Figure 4 – source data 1: selfer_theta_20kb.tsv.zip; Binned nucleotide diversity for *C. elegans, C. briggsae,* and *C. tropicalis.*

Figure 4 – source data 2: JU1373-NIC58.alignmentCoverage.tsv.zip; JU1373 and NIC58 identity and copy number variation (Minimap2 alignment).

Figure 4 – source data 3: theta_10bp.bed.zip; *C. tropicalis* fine-scale nucleotide diversity (10 bp scale).

Figure 5 – source data 1: isolateMetadata.tsv.zip; metadata for *C. tropicalis* wild isolates.

Figure 6 – source data 1: RILmating.tsv.zip; RIL outcross probability trials.

Figure 7 – source data 1: RILsegregationDistortion.tsv.zip; genotype tables at segregation distortion peaks on chromosomes I, III and V.

Figure 9 – source data 1: NIC58_JU1373_RIL.driveCrosses.tsv.zip; plate-level cross compatibility data for JU1373, NIC58 and RILs.

Figure 11 – source data 1: NIC58_JU1373_isolate.driveCrosses.tsv.zip; plate-level cross compatibility data for JU1373, NIC58 and wild isolates.

## Supplementary Files

Supplementary File 1: NIC58_rqtlCross.rda.zip; R/qtl cross object containing the NIC58 genetic map and associated RIL genotypes.

Supplementary File 2: tropicalisGenomes.zip; archive containing nuclear and mitochondrial genomes and annotations for NIC58 and JU1373.

Supplementary File 3: rawVariantCalls.zip; archive containing unfiltered variant calls for nuclear and mitochondrial genomes.

Supplementary File 4: filteredVariantCalls.zip; archive containing hard-filtered variant calls for the nuclear genome.

Supplementary File 5: processedVariantCalls.zip; archive containing hard-filtered variant calls for nuclear and mitochondrial genomes with no missing data.

Supplementary File 6: caeno_orthogroups.tsv.zip; all ortholog groupings for *Caenorhabditis* species.

## Acknowledgements

We thank Arielle Martel, Jia Shen, and Patrick Ammerman for assistance in the lab, and the Felix, Teotonio, and Rockman labs for discussion. We are grateful to Marie-Anne Felix for the use of the JU strains, for stimulating discussions with her and Henrique Teotonio, and for helpful comments on the preprint from Asher Cutter. This work was supported by R01GM121828 (MVR), R01GM089972 (MVR), R01ES029930 (ECA and MVR), Dean’s Undergraduate Research Fund grants (JY), the Centre National de la Recherche Scientifique (CNRS; CB), and the European Commission Marie Skłodowska-Curie Fellowship H2020-MSCA-IF-2017-798083 (LMN). *C. tropicalis* strain QG843 was collected under permit SEX/A-25-12 from the Republic of Panama. Sequencing data were generated by the Duke University Center for Genomic and Computational Biology and the New York University Center for Genomics and Systems Biology Core Facility, and this work was supported in part through the NYU IT High Performance Computing resources, services, and staff expertise.

## Reference

Andersen, E. C., Gerke, J. P., Shapiro, J. A., Crissman, J. R., Ghosh, R., Bloom, J. S., Felix, M.-A., & Kruglyak, L. (2012). Chromosome-scale selective sweeps shape *Caenorhabditis elegans* genomic diversity. Nature Genetics, 44(3), 285–290.

Anderson, J. L., Morran, L. T., & Phillips, P. C. (2010). Outcrossing and the maintenance of males within *C. elegans* populations. The Journal of Heredity, 101 *Suppl 1*, S62–S74. https://doi.org/10.1093/jhered/esq003

Andolfatto, P., Davison, D., Erezyilmaz, D., Hu, T. T., Mast, J., Sunayama-Morita, T., & Stern, D. L. (2011). Multiplexed shotgun genotyping for rapid and efficient genetic mapping. Genome Research, 21(4), 610–617.https://doi.org/10.1101/gr.115402.110

Ayad, L. A. K., & Pissis, S. P. (2017). MARS: improving multiple circular sequence alignment using refined sequences. BMC Genomics, 18(1), 1–10. https://doi.org/10.1186/s12864-016-3477-5

Baird, S. E., & Stonesifer, R. (2012). Reproductive isolation in *Caenorhabditis briggsae·.* Dysgenic interactions between maternal- and zygotic-effect loci result in a delayed development phenotype. Worm, 7(4), 189–195. https://doi.org/10.4161/worm.23535

Barriere, A., & Felix, M.-A. (2007). Temporal dynamics and linkage disequilibrium in natural *Caenorhabditis elegans* populations. Genetics, 176(2), 999–1011.

Barriere, A., Yang, S.-P., Pekarek, E., Thomas, C. G., Haag, E. S., & Ruvinsky, I. (2009). Detecting heterozygosity in shotgun genome assemblies: Lessons from obligately outcrossing nematodes. Genome Research, 19(3), 470–480. https://doi.org/10.1101/gr.081851.108

Bates, D., Machler, M., Bolker, B., & Walker, S. (2015). Fitting Linear Mixed-Effects Models Using Ime4. Journal of Statistical Software, Articles, 67(1), 1-48. https://doi.org/10.18637/jss.v067.i01

Baym, M., Kryazhimskiy, S., Lieberman, T. D., Chung, H., Desai, M. M., & Kishony, R. (2015). Inexpensive multiplexed library preparation for megabase-sized genomes. PloS One, 10(5), e0128036. https://doi.org/10.1371/journal.pone.0128036

Beeman, R. W., & Friesen, K. S. (1999). Properties and natural occurrence of maternal-effect selfish genes (’Medea^1^ factors) in the red flour beetle, *Tribolium castaneum*. Heredity, 82 (Pt 5), 529–534. https://doi.org/10.1038/sj.hdy.6885150

Beeman, R. W., Friesen, K. S., & Denell, R. E. (1992). Maternal-effect selfish genes in flour beetles. Science, 256(5053), 89–92. https://doi.org/10.1126/science.1566060

Ben-David, E., Burga, A., & Kruglyak, L. (2017). A maternal-effect selfish genetic element in *Caenorhabditis elegans*. Science, 356(6342), 1051–1055. https://doi.org/10.1126/science.aan0621

Ben-David, E., Pliota, P., Widen, S. A., Koreshova, A., Lemus-Vergara, T., Verpukhovskiy, P., Mandali, S., Braendle, C., Burga, A., & Kruglyak, L. (2020). Ubiquitous selfish toxin-antidote elements in Caenorhabditis species, https://doi.org/10.1101/2020.08.06.240564

Bernstein, M. R., & Rockman, M. V. (2016). Fine-Scale crossover rate variation on the *Caenorhabditis elegans* X chromosome. G3: Genes, Genomes, Genetics, 6(6), 1767–1776.

Bernt, M., Donath, A., Juhling, F., Externbrink, F., Florentz, C., Fritzsch, G., Putz, J., Middendorf, M., & Stadler, P. F. (2013). MITOS: improved de novo metazoan mitochondrial genome annotation. Molecular Phylogenetics and Evolution, 69(2), 313–319. https://doi.org/10.1016/j.ympev.2012.08.023

Brand, C. L., Larracuente, A. M., & Presgraves, D. C. (2015). Origin, evolution, and population genetics of the selfish Segregation Distorter gene duplication in European and African populations of *Drosophila melanogaster*. Evolution; International Journal of Organic Evolution, 69(5), 1271–1283. https://doi.org/10.1111/evo.12658

Brenner, S. (1974). The Genetics of *Caenorhabditis elegans*. Genetics, 77(1), 71–94. https://www.genetics.org/content/77/1/71

Briggs Gochnauer, M., & McCoy, E. (1954). Response of a soil nematode, *Rhabditis briggsae*, to antibiotics. The Journal of Experimental Zoology, 125(3), 377–406. https://doi.org/10.1002/jez.1401250302

Broman, K. W., Wu, H., Sen, S., & Churchill, G. A. (2003). R/qtl: QTL mapping in experimental crosses. Bioinformatics, 79(7), 889–890. https://doi.org/10.1093/bioinformatics/btg112

Bull, J. J. (2017). Lethal gene drive selects inbreeding. Evolution, Medicine, and Public Health, 2017(1), 1–16. https://doi.org/10.1093/emph/eow030

Bull, J. J., Remien, C. H., & Krone, S. M. (2019). Gene-drive-mediated extinction is thwarted by population structure and evolution of sib mating. *Evolution*, Medicine, and Public Health, 2079(1), 66–81. https://doi.org/10.1093/emph/eoz014

Bulmer, M. G., & Taylor, P. D. (1980). Sex ratio under the haystack model. Journal of Theoretical Biology, 86(1), 83–89. https://d0i.0rg/l0.1016/0022-5193(80)90066-1

Burt, A. (2003). Site-specific selfish genes as tools for the control and genetic engineering of natural populations. Proceedings. Biological Sciences/The Royal Society, 270(1518), 921–928. https://doi.org/10.1098/rspb.2002.2319

Champer, J., Liu, J., Oh, S. Y., Reeves, R., Luthra, A., Oakes, N., Clark, A. G., & Messer, P. W. (2018). Reducing resistance allele formation in CRISPR gene drive. Proceedings of the National Academy of Sciences of the United States of America, 775(21), 5522–5527. https://doi.org/10.1073/pnas.1720354115

Charlesworth, B. (2012). The effects of deleterious mutations on evolution at linked sites. Genetics, 790(1), 5–22. https://doi.org/10.1534/genetics.111.134288

Charlesworth, B., Nordborg, M., & Charlesworth, D. (1997). The effects of local selection, balanced polymorphism and background selection on equilibrium patterns of genetic diversity in subdivided populations. Genetical Research, 70(2), 155–174. https://doi.org/10.1017/S0016672397002954

Chaudhuri, J., Bose, N., Tandonnet, S., Adams, S., Zuco, G., Kache, V., Parihar, M., von Reuss, S. H., Schroeder, F. C., & Pires-daSilva, A. (2015). Mating dynamics in a nematode with three sexes and its evolutionary implications. Scientific Reports, 5(1), 1–11. https://doi.org/10.1038/srep17676

Chelo, I. M., Afonso, B., Carvalho, S., Theologidis, I., Goy, C., Pino-Querido, A., Proulx, S., & Teotonio, H. (2019). Partial selfing can reduce genetic loads while maintaining diversity during evolution. G3, 9, 2811–2821.

Cook, D. E., Zdraljevic, S., Roberts, J. P., & Andersen, E. C. (2017). CeNDR, the *Caenorhabditis elegans* natural diversity resource. Nucleic Acids Research, 45(D1), D650–D657. https://doi.org/10.1093/nar/gkw893

Cook, D. E., Zdraljevic, S., Tanny, R. E., Seo, B., Riccardi, D. D., Noble, L. M., Rockman, M. V., Alkema, M. J., Braendle, C., Kammenga, J. E., Wang, J., Kruglyak, L., Felix, M.-A., Lee, J., & Andersen, E. C. (2016). The Genetic Basis of Natural Variation in *Caenorhabditis elegans* Telomere Length. Genetics, 204, 371–383. https://doi.org/10.1534/genetics.116.191148

Courret, C., Chang, C.-H., Wei, K. H.-C., Montchamp-Moreau, C., & Larracuente, A. M. (2019). Meiotic drive mechanisms: lessons from *Drosophila*. Proceedings. Biological Sciences / The Royal Society, 286(1913), 20191430. https://doi.org/10.1098/rspb.2019.1430

Crombie, T. A., Zdraljevic, S., Cook, D. E., Tanny, R. E., Brady, S. C., Wang, Y., Evans, K. S., Hahnel, S., Lee, D., Rodriguez, B. C., Zhang, G., van der Zwagg, J., Kiontke, K., & Andersen, E. C. (2019). Deep sampling of Hawaiian *Caenorhabditis elegans* reveals high genetic diversity and admixture with global populations. eLife, 8, e50465. https://doi.org/10.7554/eLife.50465

Cutter, A. D. (2015). *Caenorhabditis* evolution in the wild. *BioEssays: News and Reviews in Molecular*, Cellular and Developmental Biology, 37(9), 983–995. https://doi.org/10.1002/bies.201500053

Cutter, A. D. (2019). Reproductive transitions in plants and animals: selfing syndrome, sexual selection and speciation. The New Phytologist, 224(3), 1080–1094. https://doi.org/10.1111/nph.16075

Cutter, A. D., Dey, A., & Murray, R. L. (2009). Evolution of the *Caenorhabditis elegans* genome. Molecular Biology and Evolution, 26(6), 1199–1234. https://doi.org/10.1093/molbev/msp048

Cutter, A. D., Felix, M.-A., Barriere, A., & Charlesworth, D. (2006). Patterns of nucleotide polymorphism distinguish temperate and tropical wild isolates of *Caenorhabditis briggsae*. Genetics, 173(A), 2021–2031. https://doi.org/10.1534/genetics.106.058651

Cutter, A. D., & Payseur, B. A. (2003). Selection at linked sites in the partial selfer *Caenorhabditis elegans*. Molecular Biology and Evolution, 20(5), 665–673. https://doi.org/10.1093/molbev/msg072

DePristo, M. A., Banks, E., Poplin, R., Garimella, K. V., Maguire, J. R., Hartl, C., Philippakis, A. A., del Angel, G., Rivas, M. A., Hanna, M., McKenna, A., Fennell, T. J., Kernytsky, A. M., Sivachenko, A. Y., Cibulskis, K., Gabriel, S. B., Altshuler, D., & Daly, M. J. (2011). A framework for variation discovery and genotyping using next-generation DNA sequencing data. Nature Genetics, 43(5), 491–498. https://doi.org/10.1038/ng.806

Dobin, A., Davis, C. A., Schlesinger, F., Drenkow, J., Zaleski, C., Jha, S., Batut, P., Chaisson, M., & Gingeras, T. R. (2013). STAR: ultrafast universal RNA-seq aligner. Bioinformatics, 29(1), 15–21. https://d0i.0rg/l0.1093/bioinformatics/bts635

Dolgin, E. S., Charlesworth, B., Baird, S. E., & Cutter, A. D. (2007). Inbreeding and Outbreeding Depression in *Caenorhabditis* Nematodes. Evolution; International Journal of Organic Evolution, 61(6), 1339–1352. https://www.jstor.org/stable/4621380

Dolgin, E. S., Felix, M.-A., & Cutter, A. D. (2008). Hakuna Nematoda: genetic and phenotypic diversity in African isolates of *Caenorhabditis elegans* and *C. briggsae*. Heredity, 100(3), 304–315. https://doi.org/10.1038/sj.hdy.6801079

Dougherty, E. C., & Nigon, V. (1949). A new species of the free-living nematode genus Rhabditis of interest in comparative physiology and genetics. The Journal of Parasitology, 35(11). https://www.scienceopen.com/document?vid=7f99cea0-62a6-4db5-9554-4cf1f10df029

Dowle, M., & Srinivasan, A. (2019). *data.table: Extension of ‘data.frame’.* https://CRAN.R-project.org/package=data.table

Drury, D. W., Dapper, A. L., Siniard, D. J., Zentner, G. E., & Wade, M. J. (2017). CRISPR/Cas9 gene drives in genetically variable and nonrandomly mating wild populations. Science Advances, 3(5), e1601910. https://doi.org/10.1126/sciadv.1601910

Dunn, P. K., & Smyth, G. K. (2016). *dglm: Double Generalized Linear Models.* https://CRAN.R-project.org/package=dglm

Duong, T. (2020). *ks: Kernel Smoothing.* https://CRAN.R-project.org/package=ks

Ellis, R. E. (2017). “The persistence of memory”—Hermaphroditism in nematodes. Molecular Reproduction and Development, 84(2), 144–157. https://doi.org/10.1002/mrd.22668

Emms, D. M., & Kelly, S. (2019). OrthoFinder: phylogenetic orthology inference for comparative genomics. Genome Biology, 20(1), 238. https://doi.org/10.1186/s13059-019-1832-y

Escobar, J. S., Auld, J. R., Correa, A. C., Alonso, J. M., Bony, Y. K., Coutellec, M.-A., Koene, J. M., Pointier, J.-P., Jarne, P., & David, P. (2011). PATTERNS OF MATING-SYSTEM EVOLUTION IN HERMAPHRODITIC ANIMALS: CORRELATIONS AMONG SELFING RATE, INBREEDING DEPRESSION, AND THE TIMING OF REPRODUCTION. Evolution; International Journal of Organic Evolution, 65(5), 1233–1253. https://doi.Org/10.1111/j.1558-5646.2011.01218.x

Evans, T. C., & Hunter, C. P. (2005). Translational control of maternal RNAs. In The C. elegans Research Community (Ed.), *WormBook.* https://doi.Org/doi/10.1895/wormbook.1.34.1

Felix, M.-A. (2020, August). Felix database. World Wde Worms. https://justbio.com/worldwideworms/

Felix, M.-A., & Braendle, C. (2010). The natural history of *Caenorhabditis elegans*. Current Biology: CB, 20(22), R965–R969. https://doi.Org/10.1016/j.cub.2010.09.050

Felix, M.-A., Braendle, C., & Cutter, A. D. (2014). A Streamlined System for Species Diagnosis in Caenorhabditis (Nematoda: Rhabditidae) with Name Designations for 15 Distinct Biological Species. PloS One, 9(4), e94723. https://doi.org/10.1371/journal.pone.0094723

Felix, M.-A., & Duveau, F. (2012). Population dynamics and habitat sharing of natural populations of Caenorhabditis elegans and C. briggsae. BMC Biology, 10, 59. https://d0i.0rg/10.1186/1741-7007-10-59

Felix, M.-A., Jovelin, R., Ferrari, C., Han, S., Cho, Y. R., Andersen, E. C., Cutter, A. D., & Braendle, C. (2013). Species richness, distribution and genetic diversity of *Caenorhabditis* nematodes in a remote tropical rainforest. BMC Evolutionary Biology, 73(1), 10. https://doi.org/10.1186/1471-2148-13-10

Ferrari, C., Salle, R., Callemeyn-Torre, N., Jovelin, R., Cutter, A. D., & Braendle, C. (2017). Ephemeral-habitat colonization and neotropical species richness of *Caenorhabditis* nematodes. BMC Ecology, 77(1), 43. https://doi.org/10.1186/s12898-017-0150-z

Fierst, J. L., Willis, J. H., Thomas, C. G., Wang, W., Reynolds, R. M., Ahearne, T. E., Cutter, A. D., & Phillips, P. C. (2015). Reproductive Mode and the Evolution of Genome Size and Structure in *Caenorhabditis* Nematodes. PLoS Genetics, 11(6), e1005323. https://doi.org/10.1371/journal.pgen.1005323

Fishman, L., & Sweigart, A. L. (2018). When Two Rights Make a Wrong: The Evolutionary Genetics of Plant Hybrid Incompatibilities. Annual Review of Plant Biology, 69, 707–731. https://d0i.0rg/l0.1146/annurev-arplant-042817-040113

Fradin, H., Kiontke, K., Zegar, C., Gutwein, M., Lucas, J., Kovtun, M., Corcoran, D. L., Baugh, L. R., Fitch, D. H. A., Piano, F., & Gunsalus, K. C. (2017). Genome Architecture and Evolution of a Unichromosomal Asexual Nematode. Current Biology: CB, 27(19), 2928–2939.e6. https://doi.Org/10.1016/j.cub.2017.08.038

Frezal, L., & Felix, M.-A. (2015). *C. elegans* outside the Petri dish. eLife, 4, e05849. https://doi.org/10.7554/eLife.05849

Frokjaer-Jensen, C., Davis, M. W., Hopkins, C. E., Newman, B. J., Thummel, J. M., Olesen, S.-P., Grunnet, M., & Jorgensen, E. M. (2008). Single-copy insertion of transgenes in *Caenorhabditis elegans*. Nature Genetics, 40(11), 1375–1383. https://doi.org/10.1038/ng.248

Garrison, E., Siren, J., Novak, A. M., Hickey, G., Eizenga, J. M., Dawson, E. T., Jones, W., Garg, S., Markello, C., Lin, M. F., Paten, B., & Durbin, R. (2018). Variation graph toolkit improves read mapping by representing genetic variation in the reference. Nature Biotechnology, 36(9), 875–879. https://doi.org/10.1038/nbt.4227

Gimond, C., Jovelin, R., Han, S., Ferrari, C., Cutter, A. D., & Braendle, C. (2013). Outbreeding Depression with Low Genetic Variation in Selfing *Caenorhabditis* Nematodes. Evolution; International Journal of Organic Evolution, 67(11), 3087–3101. https://doi.org/10.1111/evo.12203

Goodwillie, C., Kalisz, S., & Eckert, C. G. (2005). The Evolutionary Enigma of Mixed Mating Systems in Plants: Occurrence, Theoretical Explanations, and Empirical Evidence. Annual Review of Ecology, Evolution, and Systematics, 36(1), 47–79. https://doi.org/10.1146/annurev.ecolsys.36.091704.175539

Graustein, A., Gaspar, J. M., Walters, J. R., & Palopoli, M. F. (2002). Levels of DNA polymorphism vary with mating system in the nematode genus Caenorhabditis. Genetics, 161(1), 99–107. https://www.ncbi.nlm.nih.gov/pubmed/12019226

Grosmaire, M., Launay, C., Siegwald, M., Brugière, T., Estrada-Virrueta, L., Berger, D., Burny, C., Modolo, L., Blaxter, M., Meister, P., Felix, M.-A., Gouyon, P.-H., & Delattre, M. (2019). Males as somatic investment in a parthenogenetic nematode. Science, 363(6432), 1210–1213. https://doi.org/10.1126/science.aau0099

Grundt, Η. H., Kjølner, S., Borgen, L., Rieseberg, L. H., & Brochmann, C. (2006). High biological species diversity in the arctic flora. Proceedings of the National Academy of Sciences of the United States of America, 103(4), 972–975. https://doi.org/10.1073/pnas.0510270103

Haber, M., Schüngel, M., Putz, A., Muller, S., Hasert, B., & Schulenburg, H. (2005). Evolutionary history of *Caenorhabditis elegans* inferred from microsatellites: evidence for spatial and temporal genetic differentiation and the occurrence of outbreeding. Molecular Biology and Evolution, 22(1), 160–173. https://doi.org/10.1093/molbev/msh264

Harris, T. W., Arnaboldi, V., Cain, S., Chan, J., Chen, W. J., Cho, J., Davis, P., Gao, S., Grove, C. A., Kishore, R., Lee, R. Y. N., Muller, H.-M., Nakamura, C., Nuin, P., Paulini, M., Raciti, D., Rodgers, F. H., Russell, M., Schindelman, G., … Sternberg, P. W. (2020). WormBase: a modern Model Organism Information Resource. Nucleic Acids Research, 48(D1), D762–D767. https://doi.org/10.1093/nar/gkz920

Hedrick, P. W. (2013). Adaptive introgression in animals: examples and comparison to new mutation and standing variation as sources of adaptive variation. Molecular Ecology, 22(18), 4606—4618. https://doi.Org/10.1111/mec.12415

Hillers, K. J., Jantsch, V., Martinez-Perez, E., & Yanowitz, J. L. (2017). Meiosis. In The C. elegans Research Community (Ed.), WormBook. https://doi.Org/10.1895/wormbook.1.178.1

Hillier, L. W., Miller, R. D., Baird, S. E., Chinwalla, A., Fulton, L. A., Koboldt, D. C., & Waterston, R. H. (2007). Comparison of *C. elegans* and *C. briggsae* Genome Sequences Reveals Extensive Conservation of Chromosome Organization and Synteny. PLoS Biology, 5(7). https://doi.org/10.1371/journal.pbio.0050167

Hoff, K. J., Lomsadze, A., Borodovsky, M., & Stanke, M. (2019). Whole-Genome Annotation with BRAKER. In M. Kollmar (Ed.), Gene Prediction: Methods and Protocols (pp. 65–95). Springer New York, https://doi.org/10.1007/978-1-4939-9173-0_5

Hubley, R., Finn, R. D., Clements, J., Eddy, S. R., Jones, T. A., Bao, W., Smit, A. F. A., & Wheeler, T. J. (2016). The Dfam database of repetitive DNA families. Nucleic Acids Research, 44(D1), D81–D89. https://doi.org/10.1093/nar/gkv1272

Hunter, S., Apweiler, R., Attwood, T. K., Bairoch, A., Bateman, A., Binns, D., Bork, P., Das, U., Daugherty, L., Duquenne, L., Finn, R. D., Gough, J., Haft, D., Hulo, N., Kahn, D., Kelly, E., Laugraud, A., Letunic, I., Lonsdale, D., … Yeats, C. (2009). InterPro: the integrative protein signature database. Nucleic Acids Research, 37(Database issue), D211–D215. https://doi.org/10.1093/nar/gkn785

Igic, B., & Kohn, J. R. (2006). The Distribution of Plant Mating Systems: Study Bias Against Obligately Outcrossing Species. Evolution; International Journal of Organic Evolution, 60(5), 1098–1103. https://doi.Org/10.1111/j.0014-3820.2006.tb01186.x

Jarne, P., & Auld, J. R. (2006). Animals Mix It up Too: The Distribution of Self-Fertilization Among Hermaphroditic Animals. Evolution; International Journal of Organic Evolution, 60(9), 1816–1824. https://doi.Org/10.1111/j.0014-3820.2006.tb00525.x

Jovelin, R., Dey, A., & Cutter, A. D. (2013). Fifteen Years of Evolutionary Genomics in *Caenorhabditis elegans*. In eLS. American Cancer Society. https://doi.org/10.1002/9780470015902.a0022897

Kahle, D., & Wickham, H. (2013). ggmap: Spatial Visualization with ggplot2. In The R Journal (Vol. 5, Issue 1, pp. 144–161). https://journal.r-project.org/archive/2013-1/kahle-wickham.pdf

Kajitani, R., Toshimoto, K., Noguchi, H., Toyoda, A., Ogura, Y., Okuno, M., Yabana, M., Harada, M., Nagayasu, E., Maruyama, H., Kohara, Y., Fujiyama, A., Hayashi, T., & Itoh, T. (2014). Efficient de novo assembly of highly heterozygous genomes from whole-genome shotgun short reads. Genome Research, 24(8), 1384–1395. https://doi.org/10.1101/gr.170720.113

Kanzaki, N., Kiontke, K., Tanaka, R., Hirooka, Y., Schwarz, A., Muller-Reichert, T., Chaudhuri, J., & Pires-daSilva, A. (2017). Description of two three-gendered nematode species in the new genus *Auanema* (Rhabditina) that are models for reproductive mode evolution. Scientific Reports, 7(1), 11135. https://doi.org/10.1038/s41598-017-09871-1

Kanzaki, N., Tsai, I. J., Tanaka, R., Hunt, V. L., Liu, D., Tsuyama, K., Maeda, Y., Namai, S., Kumagai, R., Tracey, A., Holroyd, N., Doyle, S. R., Woodruff, G. C., Murase, K., Kitazume, H., Chai, C., Akagi, A., Panda, O., Ke, H.-M., … Kikuchi, T. (2018). Biology and genome of a newly discovered sibling species of *Caenorhabditis elegans*. Nature Communications, 9(1), 3216. https://doi.org/10.1038/s41467-018-05712-5

Kaur, T., & Rockman, M. V. (2014). Crossover heterogeneity in the absence of hotspots in *Caenorhabditis elegans*. Genetics, 796(1), 137–148.

Kiontke, K. C. (2005). The phylogenetic relationships of *Caenorhabditis* and other rhabditids. In The C. elegans Research Community (Ed.), WormBook. https://doi.Org/10.1895/wormbook.1.11.1

Kiontke, K. C., Felix, M.-A., Ailion, M., Rockman, M. V., Braendle, C., Pénigault, J.-B., & Fitch, D. H. A. (2011). A phylogeny and molecular barcodes for *Caenorhabditis*, with numerous new species from rotting fruits. BMC Evolutionary Biology, 77(1), 339. https://doi.org/10.1186/1471-2148-11-339

Kiontke, K. C., & Sudhaus, W. (2006). Ecology of *Caenorhabditis* species. In The C. elegans Research Community (Ed.), WormBook. https://doi.Org/10.1895/wormbook.1.37.1

Kolmogorov, M., Yuan, J., Lin, Y., & Pevzner, P. A. (2019). Assembly of long, error-prone reads using repeat graphs. Nature Biotechnology, 37(5), 540–546. https://doi.org/10.1038/s41587-019-0072-8

Koren, S., Walenz, B. P., Berlin, K., Miller, J. R., Bergman, N. H., & Phillippy, A. M. (2017). Canu: scalable and accurate long-read assembly via adaptive k-mer weighting and repeat separation. Genome Research, 27(5), 722–736. https://doi.org/10.1101/gr.215087.116

Koster, J., & Rahmann, S. (2012). Snakemake-a scalable bioinformatics workflow engine. Bioinformatics, 28(19), 2520–2522. https://doi.org/10.1093/bioinformatics/bts480

Lee, D., Zdraljevic, S., Stevens, L., Wang, Y., Tanny, R. E., Crombie, T. A., Cook, D. E., Webster, A. K., Chirakar, R., Ryan Baugh, L., Sterken, M. G., Braendle, C., Felix, M.-A., Rockman, M. V., & Andersen, E. C. (2020). Balancing selection maintains ancient genetic diversity in C. elegans (p. 2020.07.23.218420). https://doi.org/10.1101/2020.07.23.218420

Lemire, B. (2005). Mitochondrial genetics. In The C. elegans Research Community (Ed.), WormBook. https://doi.org/10.1895/wormbook.1.25.1

Li, H. (2011). A statistical framework for SNP calling, mutation discovery, association mapping and population genetical parameter estimation from sequencing data. Bioinformatics, 27(21), 2987–2993. https://doi.org/10.1093/bioinformatics/btr509

Li, H. (2018). Minimap2: pairwise alignment for nucleotide sequences. Bioinformatics, 34(18), 3094–3100. https://doi.org/10.1093/bioinformatics/bty191

Li, H., & Durbin, R. (2009). Fast and accurate short read alignment with Burrows-Wheeler transform. Bioinformatics, 25(14), 1754–1760. https://doi.org/10.1093/bioinformatics/btp324

Li, H., Handsaker, B., Wysoker, A., Fennell, T., Ruan, J., Homer, N., Marth, G., Abecasis, G., Durbin, R., & 1000 Genome Project Data Processing Subgroup. (2009). The Sequence Alignment/Map format and SAMtools. Bioinformatics, 25(16), 2078–2079. https://doi.org/10.1093/bioinformatics/btp352

Lindholm, A. K., Dyer, K. A., Firman, R. C., Fishman, L., Forstmeier, W., Holman, L., Johannesson, H., Knief, U., Kokko, H., Larracuente, A. M., Manser, A., Montchamp-Moreau, C., Petrosyan, V. G., Pomiankowski, A., Presgraves, D. C., Safronova, L. D., Sutter, A., Unckless, R. L., Verspoor, R. L., … Price, T. A. R. (2016). The Ecology and Evolutionary Dynamics of Meiotic Drive. Trends in Ecology & Evolution, 31(A), 315–326. https://doi.Org/10.1016/j.tree.2016.02.001

Li, R., Ren, X., Bi, Y., Ding, Q., Ho, V. W. S., & Zhao, Z. (2018). Comparative mitochondrial genomics reveals a possible role of a recent duplication of NADH dehydrogenase subunit 5 in gene regulation. DNA Research: An International Journal for Rapid Publication of Reports on Genes and Genomes, 25(6), 577–586. https://doi.org/10.1093/dnares/dsy026

Li, W., Jaroszewski, L., & Godzik, A. (2001). Clustering of highly homologous sequences to reduce the size of large protein databases. Bioinformatics, 17(3), 282–283. https://doi.Org/10.1093/bioinformatics/17.3.282

Lorenzen, M. D., Gnirke, A., Margolis, J., Garnes, J., Campbell, M., Stuart, J. J., Aggarwal, R., Richards, S., Park, Y., & Beeman, R. W. (2008). The maternal-effect, selfish genetic element Medea is associated with a composite Tc1 transposon. Proceedings of the National Academy of Sciences of the United States of America, 105(29), 10085–10089. https://doi.org/10.1073/pnas.0800444105

Lyttle, T. W. (1991). Segregation distorters. Annual Review of Genetics, 25, 511–557. https://d0i.0rg/10.1146/annurev.ge.25.120191.002455

Maheshwari, S., & Barbash, D. A. (2011). The Genetics of Hybrid Incompatibilities. Annual Review of Genetics, 45(1), 331–355. https://doi.org/10.1146/annurev-genet-110410-132514

Marcais, G., Delcher, A. L., Phillippy, A. M., Coston, R., Salzberg, S. L., & Zimin, A. (2018). MUMmer4: A fast and versatile genome alignment system. PLoS Computational Biology, 14, e1005944. https://doi.org/10.1371/journal.pcbi.1005944

Maupas, E. (1899). La mue et I’enkystement chez les nematodes. Arch Zoo! Exp gen,(3e. Serie), 7, 563–628.

Maupas, E. (1900). Modes et formes de reproduction des nematodes. Archives de Zoologie Experimental et Generate, 463–624.

Mayer, W. E., Herrmann, M., & Sommer, R. J. (2007). Phylogeny of the nematode genus *Pristionchus* and implications for biodiversity, biogeography and the evolution of hermaphroditism. BMC Evolutionary Biology, 7(1), 1–13. https://doi.org/10.1186/1471-2148-7-104

McKenna, A., Hanna, M., Banks, E., Sivachenko, A., Cibulskis, K., Kernytsky, A., Garimella, K., Altshuler, D., Gabriel, S., Daly, M., & DePristo, M. A. (2010). The Genome Analysis Toolkit: a MapReduce framework for analyzing next-generation DNA sequencing data. Genome Research, 20(9), 1297–1303.

Morran, L. T., Parmenter, M. D., & Phillips, P. C. (2009). Mutation load and rapid adaptation favour outcrossing over self-fertilization. Nature, 462(7271), 350–352.

Nei, M., & Li, W. H. (1979). Mathematical model for studying genetic variation in terms of restriction endonucleases. Proceedings of the National Academy of Sciences of the United States of America, 76(10), 5269–5273. https://doi.org/10.1073/pnas.76.10.5269

Nigon, V., & Felix, M.-A. (2017). History of research on *C. elegans* and other free-living nematodes as model organisms. In The C. elegans Research Community (Ed.), WormBook. https://doi.org/10.1895/wormbook.1.181.1

Nordborg, M., Charlesworth, B., & Charlesworth, D. (1996). Increased levels of polymorphism surrounding selectively maintained sites in highly selling species. Proceedings of the Royal Society of London. Series B: Biological Sciences, 263(1373), 1033–1039. https://doi.org/10.1098/rspb.1996.0152

Nordborg, M., & Donnelly, P. (1997). The Coalescent Process With Selfing. Genetics, 746(3), 1185–1195. https://www.genetics.Org/content/146/3/1185

Otto, S. P. (2009). The Evolutionary Enigma of Sex. The American Naturalist, 174(S1), S1–S14. https://doi.org/10.1086/599084

Paten, B., Earl, D., Nguyen, N., Diekhans, M., Zerbino, D., & Haussler, D. (2011). Cactus: Algorithms for genome multiple sequence alignment. Genome Research, 21(9), 1512–1528. https://doi.org/10.1101/gr.123356.111

Peters, L. L., & Barker, J. E. (1993). Novel inheritance of the murine severe combined anemia and thrombocytopenia (Scat) phenotype. Cell, 74(1), 135–142. https://doi.org/10.1016/0092-8674(93)90301-6

Prasad, A., Croydon-Sugarman, M. J. F., Murray, R. L., & Cutter, A. D. (2011). Temperature-Dependent Fecundity Associates with Latitude in *Caenorhabditis briggsae*. Evolution; International Journal of Organic Evolution, 65(1), 52–63. https://doi.org/10.1111/j.1558-5646.2010.01110.x

Presgraves, D. C. (2010). The molecular evolutionary basis of species formation. Nature Reviews. Genetics, 11(3), 175–180. https://doi.org/10.1038/nrg2718

Presgraves, D. C., Gerard, P. R., Cherukuri, A., & Lyttle, T. W. (2009). Large-scale selective sweep among Segregation Distorter chromosomes in African populations of *Drosophila melanogaster*. PLoS Genetics, 5(5), e1000463. https://doi.org/10.1371/journal.pgen.1000463

Price, T. A. R., Verspoor, R., & Wedell, N. (2019). Ancient gene drives: an evolutionary paradox. Proceedings. Biological Sciences / The Royal Society, 286(1917), 20192267. https://doi.org/10.1098/rspb.2019.2267

Quinlan, A. R., & Hall, I. M. (2010). BEDTools: a flexible suite of utilities for comparing genomic features. Bioinformatics, 26(6), 841–842. https://doi.org/10.1093/bioinformatics/btq033

R Core Team. (2018). R: A Language and Environment for Statistical Computing. R Foundation for Statistical Computing. https://www.R-project.org/

Richaud, A., Zhang, G., Lee, D., Lee, J., & Felix, M.-A. (2018). The Local Coexistence Pattern of Selfing Genotypes in *Caenorhabditis elegans* Natural Metapopulations. Genetics, 208(2), 807–821. https://doi.org/10.1534/genetics.117.300564

Robertson, S., & Lin, R. (2015). The Maternal-to-Zygotic Transition in *C. elegans*. In H. D. Lipshitz (Ed.), Current Topics in Developmental Biology (Vol. 113, pp. 1–42). Academic Press. https://doi.org/10.1016/bs.ctdb.2015.06.001

Rockman, M. V., & Kruglyak, L. (2009). Recombinational Landscape and Population Genomics of Caenorhabditis elegans. PLoS Genetics, 5(3), e1000419. https://doi.org/10.1371/journal.pgen.1000419

Rockman, M. V., Skrovanek, S. S., & Kruglyak, L. (2010). Selection at linked sites shapes heritable phenotypic variation in *C. elegans*. Science, 330(6002), 372–376. https://doi.org/10.1126/science.1194208

Ross, J. A., Koboldt, D. C., Staisch, J. E., Chamberlin, Η. M., Gupta, B. P., Miller, R. D., Baird, S. E., & Haag, E. S. (2011). *Caenorhabditis briggsae* Recombinant Inbred Line Genotypes Reveal Inter-Strain Incompatibility and the Evolution of Recombination. PLoS Genetics, 7(7), e1002174. https://doi.org/10.1371/journal.pgen.1002174

Ruan, J., & Li, H. (2019). *Fast and accurate long-read assembly with wtdbg2* (p. 530972). https://doi.org/10.1101/530972

Saito, T. T., & Colaiacovo, M. P. (2017). Regulation of Crossover Frequency and Distribution during Meiotic Recombination. Cold Spring Harbor Symposia on Quantitative Biology, 82, 223–234. https://doi.org/10.1101/sqb.2017.82.034132

Seemann, T. (2014). Prokka: rapid prokaryotic genome annotation. Bioinformatics, 30(14), 2068–2069. https://doi.org/10.1093/bioinformatics/btu153

Seidel, H. S., Ailion, M., Li, J., van Oudenaarden, A., Rockman, M. V., & Kruglyak, L. (2011). A Novel Sperm-Delivered Toxin Causes Late-Stage Embryo Lethality and Transmission Ratio Distortion in *C. elegans*. PLoS Biology, 9(7), e1001115. https://doi.org/10.1371/journal.pbio.1001115

Seidel, H. S., Rockman, M. V., & Kruglyak, L. (2008). Widespread genetic incompatibility in *C. elegans* maintained by balancing selection. Science, 379(5863), 589–594. https://d0i.0rg/10.1126/science.1151107

Seppey, M., Manni, M., & Zdobnov, E. M. (2019). BUSCO: Assessing Genome Assembly and Annotation Completeness. In M. Kollmar (Ed.), Gene Prediction: Methods and Protocols (pp. 227–245). Springer, New York, https://doi.org/10.1007/978-1-4939-9173-0_14

Shi, C., Runnels, A. M., & Murphy, C. T. (2017). Mating and male pheromone kill *Caenorhabditis* males through distinct mechanisms. eLife, 6. https://doi.org/10.7554/eLife.23493

Sivasundar, A., & Hey, J. (2005). Sampling from natural populations with RNAi reveals high outcrossing and population structure in *Caenorhabditis elegans*. Current Biology: CB, 75(17), 1598–1602. https://doi.Org/10.1016/j.cub.2005.08.034

Slowikowski, K. (2020). *ggrepel: Automatically Position Non-Overlapping Text Labels with “ggplot2.”* https://CRAN.R-project.org/package=ggrepel

Smit, A., Hubley, R., & Green, P. (n.d.). *RepeatMasker.* Retrieved August 6, 2020, from http://www.repeatmasker.org/

Stanke, M., Schöffmann, O., Morgenstern, B., & Waack, S. (2006). Gene prediction in eukaryotes with a generalized hidden Markov model that uses hints from external sources. BMC Bioinformatics, 1, 62. https://doi.org/10.1186/1471-2105-7-62

Stankowski, S., Chase, M. A., Fuiten, A. M., Rodrigues, M. F., Ralph, P. L., & Streisfeld, M. A. (2008). Wdespread selection and gene flow shape the genomic landscape during a radiation of monkeyflowers. PLoS Biology, 17(7), e3000391. https://doi.org/10.1371/journal.pbio.3000391

Stevens, L., Felix, M.-A., Beltran, T., Braendle, C., Caurcel, C., Fausett, S., Fitch, D., Frezal, L., Gosse, C., Kaur, T., Kiontke, K., Newton, M. D., Noble, L. M., Richaud, A., Rockman, M. V., Sudhaus, W., & Blaxter, M. (2019). Comparative genomics of 10 new *Caenorhabditis* species. Evolution Letters, 3(2), 217–236. https://doi.org/10.1002/evl3.110

Stewart, A. D., & Phillips, P. C. (2002). Selection and maintenance of androdioecy in Caenorhabditis elegans. Genetics, 160(3), 975–982.

Stiernagle, T. (2006). Maintenance of *C. elegans.* In The C. elegans Research Community (Ed.), WormBook. https://doi.org/10.1895/wormbook.1.101.1

Subirana, J. A., & Messeguer, X. (2017). Evolution of Tandem Repeat Satellite Sequences in Two Closely Related *Caenorhabditis* Species. Diminution of Satellites in Hermaphrodites. Genes, 8(12). https://doi.org/10.3390/genes8120351

Sunnucks, P., & Hales, D. F. (1996). Numerous transposed sequences of mitochondrial cytochrome oxidase l-ll in aphids of the genus *Sitobion* (Hemiptera: Aphididae). Molecular Biology and Evolution, 13(3), 510–524. https://doi.org/10.1093/oxfordjournals.molbev.a025612

Sweigart, A. L., Brandvain, Y., & Fishman, L. (2019). Making a Murderer: The Evolutionary Framing of Hybrid Gamete-Killers. Trends in Genetics: TIG, 35(4), 245–252. https://doi.Org/10.1016/j.tig.2019.01.004

Teotonio, H., Carvalho, S., Manoel, D., Roque, M., & Chelo, I. M. (2012). Evolution of Outcrossing in Experimental Populations of *Caenorhabditis elegans*. PloS One, 7(4), e35811. https://doi.org/10.1371/journal.pone.0035811

Teotonio, H., Manoel, D., & Phillips, P. C. (2006). Genetic variation for outcrossing among Caenorhabditis elegans *isolates*. Evolution; International Journal of Organic Evolution, 60(6), 1300–1305.

Teterina, A. A., Willis, J. H., & Phillips, P. C. (2020). Chromosome-Level Assembly of the *Caenorhabditis remanei* Genome Reveals Conserved Patterns of Nematode Genome Organization. Genetics, 214(4), 769–780. https://doi.org/10.1534/genetics.119.303018

Thomas, C. G., Wang, W., Jovelin, R., Ghosh, R., Lomasko, T., Trinh, Q., Kruglyak, L., Stein, L. D., & Cutter, A. D. (2015). Full-genome evolutionary histories of selfing, splitting, and selection in *Caenorhabditis*. Genome Research, 25(5), 667–678. https://doi.org/10.1101/gr.187237.114

Thompson, O. A., Snoek, L. B., Nijveen, H., Sterken, M. G., Volkers, R. J. M., Brenchley, R., Van’t Hof, A., Bevers, R. P. J., Cossins, A. R., Yanai, I., Hajnal, A., Schmid, T., Perkins, J. D., Spencer, D., Kruglyak, L., Andersen, E. C., Moerman, D. G., Hillier, L. W., Kammenga, J. E., & Waterston, R. H. (2015). Remarkably Divergent Regions Punctuate the Genome Assembly of the *Caenorhabditis elegans* Hawaiian Strain CB4856. Genetics, 200(3), 975–989.

Ting, J. J., Woodruff, G. C., Leung, G., Shin, N.-R., Cutter, A. D., & Haag, E. S. (2014). Intense Sperm-Mediated Sexual Conflict Promotes Reproductive Isolation in *Caenorhabditis* Nematodes. PLoS Biology, 12(7), e1001915. https://doi.org/10.1371/journal.pbio.1001915

van den Brand, T. (2020). *ggh4x: Hacks for “ggplot2.”*

Vaser, R., & Sikic, M. (2019). *Yet another de novo genome assembler (p.* 656306). https://doi.org/10.1101/656306

Vasimuddin, Md, Misra, S., Li, H., & Aluru, S. (2019). Efficient Architecture-Aware Acceleration of BWA-MEM for Multicore Systems. In *arXiv[cs.DC]*, arXiv. http://arxiv.org/abs/1907.12931

Wade, M. J., & Beeman, R. W. (1994). The population dynamics of maternal-effect selfish genes. Genetics, 138(4), 1309–1314. https://www.ncbi.nlm.nih.gov/pubmed/7896109

Walker, B. J., Abeel, T., Shea, T., Priest, M., Abouelliel, A., Sakthikumar, S., Cuomo, C. A., Zeng, Q., Wortman, J., Young, S. K., & Earl, A. M. (2014). Pilon: an integrated tool for comprehensive microbial variant detection and genome assembly improvement. PloS One, 9(11), e112963. https://doi.org/10.1371/journal.pone.0112963

Weichenhan, D., Kunze, B., Traut, W., & Winking, H. (1998). Restoration of the Mendelian transmission ratio by a deletion in the mouse chromosome 1 HSR. Genetical Research, 71(2), 119–125. https://doi.org/10.1017/s0016672398003206

Weichenhan, D., Traut, W., Kunze, B., & Winking, H. (1996). Distortion of Mendelian recovery ratio for a mouse HSR is caused by maternal and zygotic effects. Genetical Research, 68(2), 125-129. https://doi.org/10.1017/s0016672300034017

Wei, Q., Zhao, Y., Guo, Y., Stomel, J., Stires, R., & Ellis, R. E. (2014). Co-option of alternate sperm activation programs in the evolution of self-fertile nematodes. Nature Communications, 5, 5888. https://doi.org/10.1038/ncomms6888

Wickham, Η. (2016). ggplot2: Elegant Graphics for Data Analysis. Springer-Verlag New York. https://ggplot2.tidyverse.org

Wickham, H., Francis, R., Henry, L., & Muller, K. (2020). *dplyr: A Grammar of Data Manipulation.* https://CRAN.R-project.org/package=dplyr

Wickham, H., & Henry, L. (2020). *tidyr: Tidy Messy Data.* https://CRAN.R-project.org/package=tidyr

Wck, R. R., Judd, L. M., Gorrie, C. L., & Holt, K. E. (2017). Unicycler: Resolving bacterial genome assemblies from short and long sequencing reads. PLoS Computational Biology, 13(6), e1005595. https://doi.org/10.1371/journal.pcbi.1005595

Wcky, C., Villeneuve, A. M., Lauper, N., Codourey, L., Tobler, H., & Muller, F. (1996). Telomeric repeats (TTAGGC)n are sufficient for chromosome capping function in *Caenorhabditis elegans*. Proceedings of the National Academy of Sciences of the United States of America, 93(17), 8983–8988. https://doi.org/10.1073/pnas.93.17.8983

Wilson, D. S., & Colwell, R. K. (1981). Evolution of Sex Ratio in Structured Demes. Evolution; International Journal of Organic Evolution, 35(5), 882–897. https://doi.org/10.2307/2407858

Winking, H., Weith, A., Boldyreff, B., Moriwaki, K., Fredga, K., & Traut, W. (1991). Polymorphic HSRs in chromosome 1 of the two semispecies *Mus musculus musculus* and *M. m. domesticus* have a common origin in an ancestral population. Chromosoma, 100(3), 147–151. https://doi.org/10.1007/BF00337242

Woodruff, G. C., & Teterina, A. A. (2020). Degradation of the repetitive genomic landscape in a close relative of *C. elegans*. Molecular Biology and Evolution. https://doi.org/10.1093/molbev/msaa107

Yin, D., Schwarz, E. M., Thomas, C. G., Felde, R. L., Korf, I. F., Cutter, A. D., Schartner, C. M., Ralston, E. J., Meyer, B. J., & Haag, E. S. (2018). Rapid genome shrinkage in a self-fertile nematode reveals sperm competition proteins. Science, 359(6371), 55–61. https://doi.org/10.1126/science.aao0827

Yoshimura, J., Ichikawa, K., Shoura, M. J., Artiles, K. L., Gabdank, I., Wahba, L., Smith, C. L., Edgley, M. L., Rougvie, A. E., Fire, A. Z., Morishita, S., & Schwarz, E. M. (2019). Recompleting the *Caenorhabditis* elegans genome. Genome Research. https://doi.org/10.1101/gr.244830.118

Zeileis, A., Leisch, F., Hornik, K., & Kleiber, C. (2002). strucchange: An R Package for Testing for Structural Change in Linear Regression Models. *Journal of Statistical Software*, Articles, 7(2), 1–38. https://doi.org/10.18637/jss.v007.i02

Zhao, Y., Tan, C.-H., Krauchunas, A., Scharf, A., Dietrich, N., Warnhoff, K., Yuan, Z., Druzhinina, M., Gu, S. G., Miao, L., Singson, A., Ellis, R. E., & Kornfeld, K. (2018). The zinc transporter ZIPT-7.1 regulates sperm activation in nematodes. PLoS Biology, 16(6), e2005069. https://doi.org/10.1371/journal.pbio.2005069

